# Signature reversion of three disease-associated gene signatures prioritizes cancer drug repurposing candidates

**DOI:** 10.1101/2023.03.10.532074

**Authors:** Jennifer L. Fisher, Elizabeth J. Wilk, Vishal H. Oza, Timothy C. Howton, Victoria Flanary, Amanda D. Clark, Anita B. Hjelmeland, Brittany N. Lasseigne

## Abstract

Drug repurposing is promising because approving a drug for a new indication requires fewer resources than approving a new drug. Signature reversion detects drug perturbations most inversely related to the disease-associated gene signature to identify drugs that may reverse that signature. We assessed the performance and biological relevance of three approaches for constructing disease-associated gene signatures (i.e, limma, DESeq2, and MultiPLIER) and prioritized the resulting drug repurposing candidates for four low-survival human cancers. Our results were enriched for candidates that had been used in clinical trials or performed well in the PRISM drug screen. Additionally, we found that pamidronate and nimodipine, drugs predicted to be efficacious against the brain tumor glioblastoma (GBM), inhibited the growth of a GBM cell line and cells isolated from a patient derived xenograft (PDX). Our results demonstrate that by applying multiple disease-associated gene signature methods, we prioritized several drug repurposing candidates for low-survival cancers.

**Graphical Abstract:** 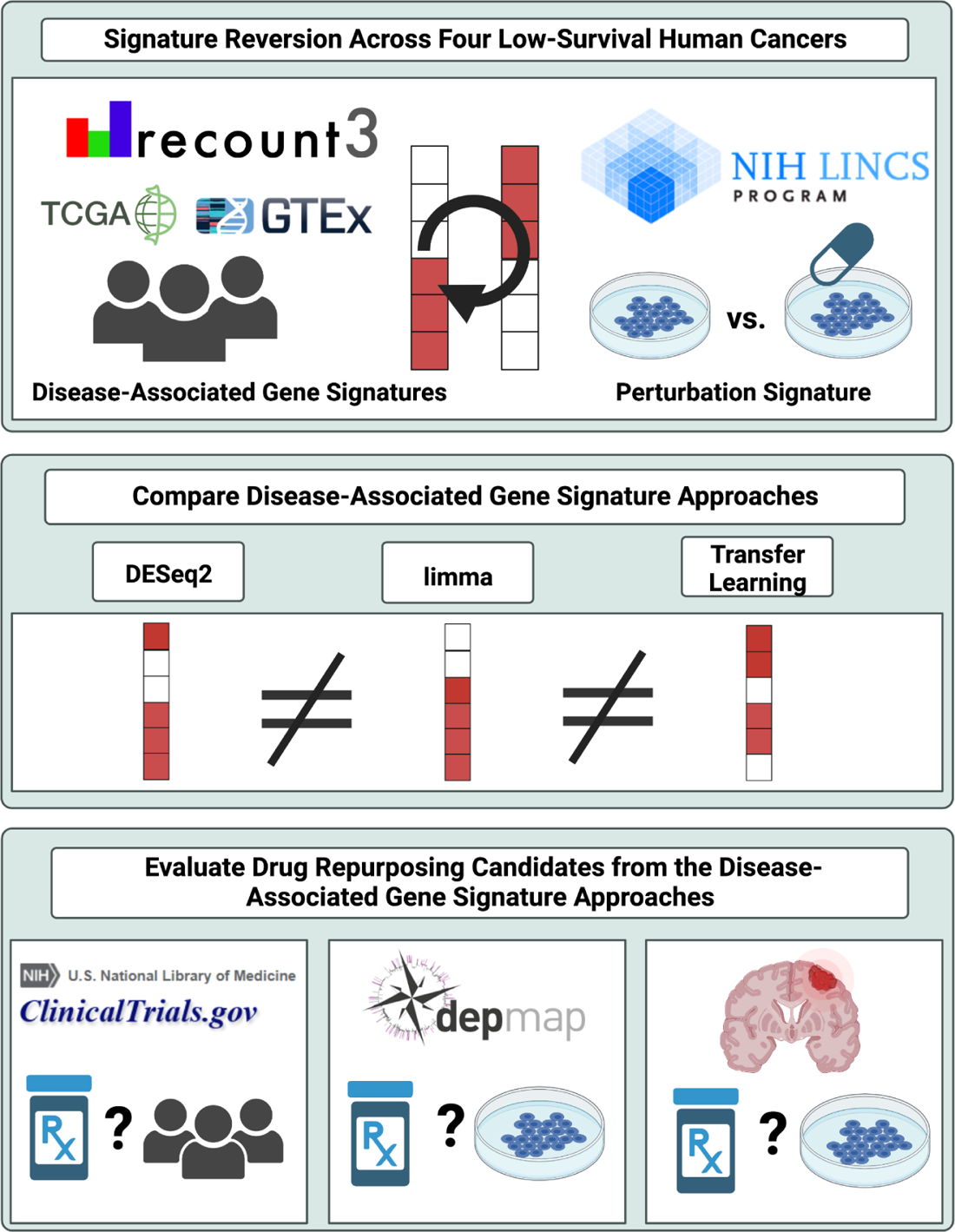

## Introduction

New therapeutic options are critical for improving cancer survival rates.^1^ While novel drug discovery can be effective, developing these new drugs costs billions of US dollars on average and can take upwards of 13 years for FDA approval.^2, 3^ Additionally, 95% of phase I clinical trial oncology drug candidates are ultimately not approved due to toxicity or inefficacy.^2, 4^ However, drug repurposing (identifying novel indications for previously approved drugs) is a promising alternative to drug discovery. As the candidates have already passed toxicity testing in previous clinical trials, drug repurposing reduces both the time and cost needed for approval for a new indication.^5, 6^ This approach successfully identified oncology and non-oncology drugs for novel cancer applications. For example, imatinib, a tyrosine kinase inhibitor for chronic myeloid leukemia, has been repurposed for treating gastrointestinal stromal tumors, and rapamycin, originally used as an immunosuppressant for kidney transplants, was repurposed for the treatment of renal cell carcinoma. ^2, 7, 8^

Identifying prioritized drugs for future cancer clinical trials requires identifying both the drug target(s) associated with the disease process and the drug predicted to perturb those target(s). Towards those goals, several computational drug repurposing methods were developed that leverage optimized algorithms and high-performance computing systems to prioritize novel drug repurposing candidates more quickly and with less expense than exhaustive large-scale experimental approaches like phenotypic drug screens.^5^ One such approach is signature reversion, a computational drug repurposing method used to prioritize drug repurposing candidates for the treatment of several cancers that were successfully validated in cell culture systems or mouse xenograft models.^9–12^ Signature reversion identifies drugs predicted to reverse disease-associated gene signatures (i.e., gene expression differences between disease and control tissue) by determining which cell line drug perturbation signatures (i.e., gene expression differences before and after drug treatment) are most inverse from the disease-associated gene signature.^13–15^ While previous studies have investigated how signature reversion methods perform^9, 11, 16–18^, understanding the impact of different approaches for selecting a disease-associated gene signature on downstream signature reversion is critical for further candidate prioritization and interpretation to advance drug repurposing studies.^11^

Here we applied and evaluated three approaches for selecting a disease-associated gene signature for downstream signature reversion (**Supplemental Figure 1**). Two approaches involved developing a disease-associated gene signature from differential gene expression analysis using limma and DESeq2, respectively.^19, 20^ Limma and DESeq2 consistently perform well in differential gene expression analysis evaluations, incorporate covariates to reduce technical influences such as batch effects^21, 22^, and are widely used in cancer signature reversion drug repurposing applications.^9–12^ For the third approach, we developed disease-associated gene signatures built on transfer learning latent variables. In this case, the transfer learning approach transfers biologically meaningful linear combinations of gene expression patterns (known as latent variables) from large databases to the tumor and control gene expression profiles.^23^ To evaluate these methods in a cancer drug repurposing context, we focused on cancers with gene expression profiles available from The Cancer Genome Atlas (TCGA) that also had the lowest survival. These four cancers [pancreatic adenocarcinoma (PAAD, 5-year survival < 8%)^24^, liver hepatocellular carcinoma (LIHC, 5-year survival <18%)^25^, glioblastoma (GBM, 5-year survival <4%)^26^, and lung adenocarcinoma (LUAD, 5-year survival = 5% for metastatic cases)] require novel drug candidates desperately.^27, 28^

We demonstrated that these three approaches for selecting a disease-associated gene signature for downstream signature reversion each captured enough unique biology to prioritize mostly different drug repurposing candidates. By comparing the identified drug candidates to drugs that have already progressed to clinical trials and the PRISM drug screen for each cancer, we validated these approaches for generating hypotheses for future in vitro and in vivo experiments. As a proof of concept, we further demonstrated that two novel drug repurposing candidates, which we predicted would be beneficial for GBM treatment (nimodipine and pamidronate), inhibited GBM growth in vitro. In addition to providing prioritized drug repurposing candidates for PAAD, LIHC, and LUAD, we conclude that by incorporating multiple approaches for selecting a disease-associated gene signature for downstream signature reversion, additional viable cancer drug repurposing candidates can be identified.

## Results

### Selection and evaluation of disease-associated gene signatures for each low survival cancer

In order to identify drug repurposing candidates, we first applied and evaluated three approaches for selecting a disease-associated gene signature for downstream signature reversion. For each cancer (**Supplemental File 1**), we identified disease-associated gene signatures by comparing TCGA tumor to TCGA and GTEx non-cancer tissue gene expression profiles with two differential gene expression methods (limma and DESeq2) and a transfer learning approach, MultiPLIER.^21–23^ Briefly, the DESeq2 algorithm modeled the gene expression counts as a negative binomial distribution and generated generalized linear models to determine differentially expressed genes while limma’s approach uses gene-wise linear models.^19, 20^ These differential gene expression approaches provided adjusted p-values and log2 fold changes for gene expression between conditions that have been used to develop disease-associated gene signatures.^9–12^ To develop a disease-associated gene signature via a transfer learning approach, we first validated the transfer of information via the gene labels of the transfer learning approach (see ‘Differential latent variable analysis’ in <u>Methods</u>) and then identified the latent variables that are different between cancer and non-tumor tissue via latent variable-wise linear models with limma.^19, 23^ From these latent variables, we determined the genes that contributed the most to the latent variables (i.e., have the highest weights in the latent variable linear gene expression equation) in order to develop the disease-associated gene signature.

We next compared the disease-associated gene signature sets identified by DESeq2, limma, and transfer learning within each cancer. First, we calculated the Spearman correlation between log2 fold change and the adjusted p-value from DESeq2 and limma. We found that all of the correlations were significant (linear regression model p-values < 2.2e-16) (**Supplemental Figure 2**). While the log2 fold changes had a correlation coefficient between 0.69 and 0.94 for each cancer, the correlation coefficient for the adjusted p-values was between 0.55 and 0.72 (**Figure 1A**). This suggests that the differences between the genes included in the DESeq2 and limma disease-associated signatures were not due to differences in log2 fold change. Instead, gene inclusion was driven by differences in the adjusted p-value methods (**Figure 1A**). Second, to compare the DESeq2 and limma disease-associated gene signatures with the disease-associated gene signature identified by transfer learning, we labeled the transfer learning genes in the volcano plots of the DESeq2 and limma differential expression analyses. The volcano plots demonstrated that the transfer learning genes did not have larger absolute log2 fold changes compared to genes identified as significantly differentially expressed by either DESeq2 or limma (**Figure 1B-I**). This underscores that the genes with the highest weights in the differential expressed latent variables identified by transfer learning were not the top differentially expressed genes identified by high absolute log2 fold change. Also, several transfer learning genes had non-significant adjusted p-values in the DESeq2 analysis (i.e., ∼31% in GBM, ∼16% in LIHC, ∼9% in LUAD, ∼73% in PAAD) or in the limma analysis (i.e., ∼45% in GBM, ∼5% in LIHC, ∼11% in LUAD, and ∼61% in PAAD). In total, these results highlight that each approach for selecting a disease-associated gene signature identified gene sets that were largely non-overlapping. In fact, the overlap in the disease-associated gene signatures was 0.8%, 0%, 0%, and 1% for GMB, LIHC, LUAD, and LUAD, respectively (**Supplemental Figure 3**).

**Figure 1:**
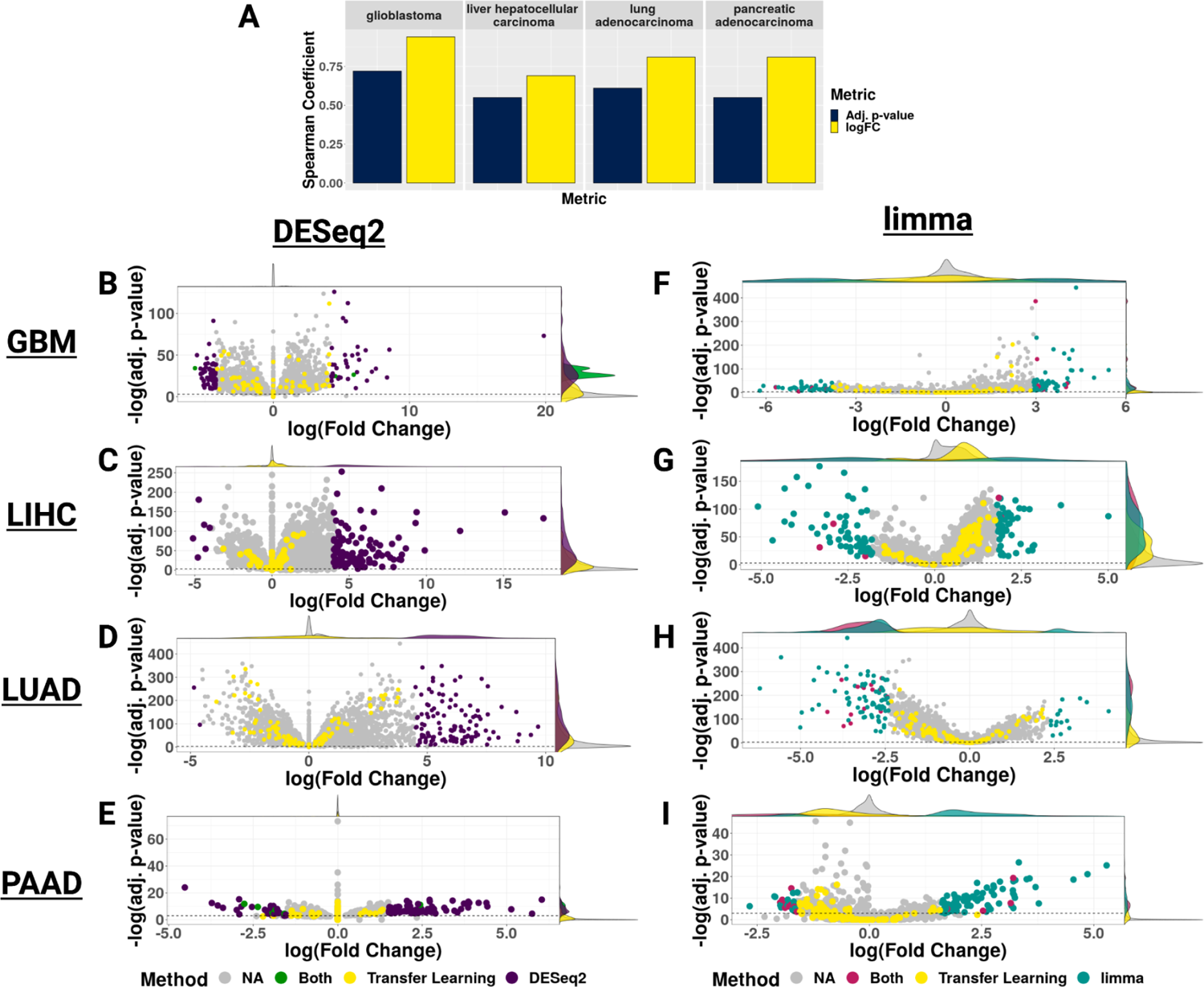
Comparison of gene inclusion in disease-associated gene signatures for downstream signature reversion. **A)** Bar plot of the Spearman correlation of adjusted p-values and log fold changes between the disease-associated differential gene expression signatures across each cancer. Volcano plots of DESeq2 calculated log fold change and adjusted p-values to compare with transfer learning disease-associated gene signature genes identified across each cancer (purple = DESeq2 disease-associated signature genes, yellow = transfer learning disease-associated genes, green = genes shared by both the DESeq2 and transfer learning disease-associated gene signatures, gray = genes not in the DESeq2 or transfer learning disease-associated gene signatures) for **B)** GBM, **C)** LIHC, **D)** LUAD, and **E)** PAAD. Volcano plots of limma calculated log fold change and adjusted p-values to compare with transfer learning disease-associated gene signature genes identified across each cancer (teal = limma disease-associated signature genes, yellow = transfer learning disease-associated genes, green = genes shared by both the limma and transfer learning disease-associated gene signatures, gray = genes not in the limma or transfer learning disease-associated gene signatures) for **F)** GBM, **G)** LIHC, **H)** LUAD, and **I)** PAAD.

Next, we used the STRING database to construct a protein-protein interaction (PPI) network to assess the degree (the number of immediate neighbors in a network), betweenness (quantification of the node importance in information flow), and eigenvector centrality (a measure of the node’s degree along the degree of neighboring nodes) of the genes in each of the disease-associated signatures.^29^ Previous studies have shown that FDA-approved or Phase 4 drug targets have higher network centrality than other genes in PPI networks because of higher degree, higher betweenness, higher bridge centrality (a metric that describes nodes that connect modular subregions of a network), lower average shortest path and lower topological coefficient (a measure for the extent to which a node shares neighbors with other nodes) network centrality metrics.^30–33^ Therefore, we wanted to compare the PPI network properties of the unique genes selected for each cancer’s disease-associated signature. We found that the unique genes identified in the transfer learning disease-associated signature have a significantly higher degree, betweenness, and eigenvector centrality than the unique genes identified by either DESeq2 or limma in GBM, LUAD, and LIHC (Kruskal-Wallis and two-tailed Wilcox tests followed by a Bonferroni p-value adjustment, adjusted p-values <0.05) (**Supplemental Figures 4-7**). However, in PAAD, both limma’s and transfer learning’s unique disease-associated genes were not significantly different from one another for all three centrality metrics (**Supplemental Figure 7**). While PPI network degree, betweenness, and eigenvector centrality metrics for the unique genes identified by DESeq2 were the lowest, followed by those from limma, and then transfer learning, these comparisons were not always statistically significant. Our results highlight that transfer learning may identify disease-associated genes with a higher PPI degree, betweenness, and eigenvector centrality than DESeq2 or limma. We further investigated the centrality metrics (i.e., degree, betweenness, and eigenvector centrality) of the highest-weighted genes in all of the latent variables used for transfer learning (n=385). We found that the 10 highest weighted genes for each latent variable (i.e. the most important genes for defining that latent variable) had significantly higher degree, betweenness, and eigenvector centrality than genes that were not in the 10 highest weighted genes for any latent variable (**Supplemental Figure 8**). Therefore, by using the top-weighted genes from the latent variables that we identified as significantly different between cancer and non-disease control samples, the transfer learning approach might have been biased to select disease-associated gene signatures with higher centrality genes (based on degree, betweenness, and eigenvector centrality from the disease-associated gene signatures and the top 10 highest weighted genes from all the 385 latent variables) than in the disease-associated signatures constructed from the DESeq2 or limma differential expression analyses.

We next asked if the three approaches identified distinct genes from common pathways. Therefore, we applied functional enrichment analysis of gene sets from Gene Ontology (GO)^34^, KEGG^35^, Reactome^36^, WikiPathways^37^, TRANSFAC^38^, miRTarBase^39^, Human Protein Atlas^40^, and The Comprehensive Resource of Mammalian Protein Complexes (CORUM)^41^ to identify any pathways or gene sets enriched in each disease-associated gene signature.^42^ We observed that 10% or less of pathway terms were shared across all three disease-associated gene signatures by cancer (**Supplemental Figures 9-12**). The largest was a ∼9% overlap in both the up- and down-regulated pathways for the GBM disease-associated gene signatures (**Supplemental Figure 9**). To determine how similarly enriched biological process terms were between the three approaches within a cancer^43^, we applied GO semantic similarity. We divided enriched GO terms into subgroups (GO_Group) based on semantic similarity and a common parent term in the GO graph to group similar GO terms together. Based on this hierarchical clustering of GO terms, we found similarities between the disease-associated gene signatures within a cancer (**Figure 2**). However, we also found enrichment of biological processes previously implicated in disease progression or therapeutic relevance that were not shared between all the disease-associated gene signatures. For example, in the GBM up-regulated enriched GO terms (**Figure 2A**), we found that all three approaches had terms associated with cell adhesion, extracellular matrix organization, and positive regulation of cell population proliferation, pathways which are all known to be perturbed in cancer. However, terms were also associated only with a specific approach’s gene set. For example, pro-inflammatory response in microglia has previously been associated with GBM progression by single-cell studies.^44^ Several pro-inflammatory gene sets were enriched in the limma disease-associated gene signature, as were terms related to angiogenesis (i.e., blood vessel development, circulatory system development, vasculature development). Angiogenesis is a common molecular feature of GBM and the therapeutic target of bevacizumab, which is FDA approved for recurrent GBMs.^45^ However, the down-regulated GO terms enriched for each of the three disease-associated gene sets were more similar (**Figure 2B**). All three analyses also identified pathways associated with dysfunctional brain tissue (e.g., chemical synapse transmission, regulation of membrane potential, and transmembrane transport) and nervous system development, further highlighting signal de-differentiation that has been associated with therapy resistance (**Figure 2B**).^46^

**Figure 2:**
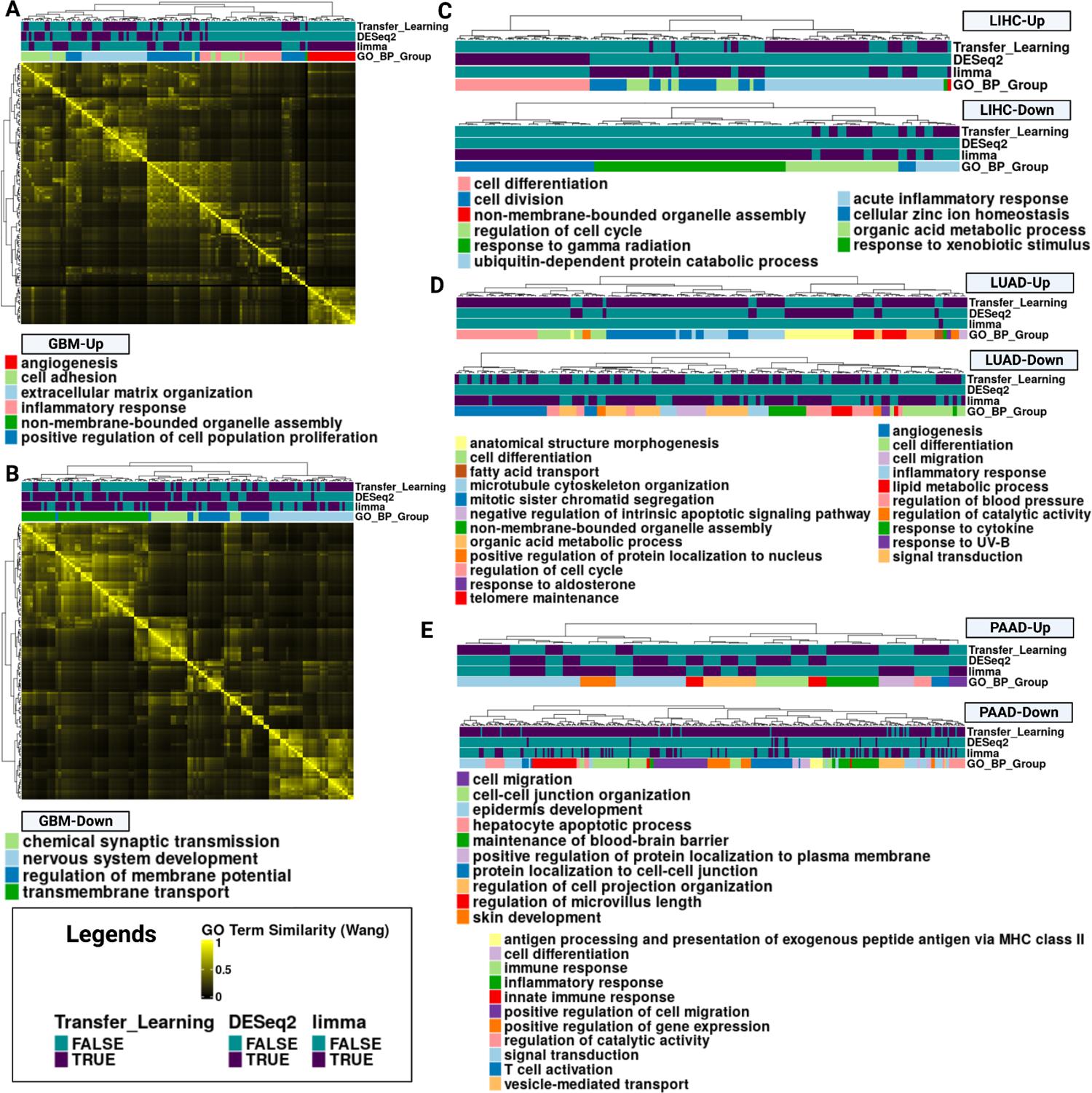
Functional enrichment analysis of disease-associated gene signatures used for signature reversion. **A)** Heatmap of the Gene Ontology (GO) term semantic similarity (Wang method) of the up-regulated enriched GO Biological Process terms from the GBM disease-associated gene signatures. Each term is associated with a disease-associated gene signature if the row for that method is purple. In addition, the Gene Ontology Biological Process terms were grouped together based on common parent terms, and the different groups are indicated in the GO_BP_Group. **B)** A heatmap of the GO term semantic similarity (Wang method) of down-regulated enriched Gene Ontology Biological Process terms for GBM. **C-H)** Dendrogram from the up and down-regulated GO Biological Process terms for LIHC, LUAD, and PAAD cancer, respectively. Up-regulated terms are plotted on top and down-regulated terms on the bottom. The legend for the up-regulated GO term groups are listed at the top and left. The down-regulated terms are listed at the bottom and the right.

While we identified similar GO terms represented in the disease-associated gene signatures across cancers based on the GO similarity clustering, there were also distinct GO term subgroups identified by only one disease-associated gene signature. We were interested in investigating the GO term groups that differed between the disease-associated gene signatures, as these may drive the differences in the drug candidate sets. For example, we found that the LIHC disease-associated gene signature from the transfer learning approach contained up-regulated GO terms unique for the cell differentiation subgroup compared to the other two signatures (**Figure 2C**). For LIHC down-regulated GO terms, we found that the disease-associated genes from the transfer learning methodology were enriched for terms related to the response to xenobiotic stimulus (**Figure 2C**). The disease-associated gene signature from the limma and transfer learning approach consisted of enriched GO terms related to 3 other GO term subgroups (**Figure 2C**). In LUAD (**Figure 2D**), we found the following distinct GO term subgroups from the respective disease-associated gene signatures: upregulation of anatomical structure morphogenesis (DESeq2), down-regulation of regulation of response to UV-B (limma), and down-regulation of lipid metabolic processes, as well as up-regulation of 5 additional GO term subgroups (Transfer Learning). For PAAD, up-regulated enriched GO terms included: regulation of the microvillus length (DESeq2), positive regulation of the protein localization to the plasma membrane, cell migration, & skin development (limma), and maintenance of the blood-brain barrier, hepatocyte apoptotic process, & protein localization to cell-cell junction (Transfer Learning) (**Figure 2E**). Interestingly, the only two distinct GO term subgroups for down-regulated PAAD disease-associated gene signatures were enriched in the transfer learning signature: antigen processing and presentation of exogenous peptide antigen via MHC class II and cell differentiation (**Figure 2E**). Overall, based on the similarity clustering of the enriched GO terms, these results highlight that the disease-associated signatures were enriched for similar and distinct GO terms. Many of these distinct GO term groups are associated with disease progression or response to therapy.^47–53^ Thus, each approach for defining disease-associated gene signature identified different aspects of cancer biology to target via signature reversion.

### Identifying prioritized drug repurposing candidates for each cancer

To perform signature reversion, we applied the LINCS methodology in the SignatureSearch R package to identify drug repurposing candidates for each of the four cancers using each of the three disease-associated signatures.^13, 54^ We considered a drug as a potential candidate if it had a negative normalized connectivity score (i.e., the perturbation signature was inverse to the disease-associated gene signature), a false discovery rate (FDR) less than 0.05, and a Tau value (describes the overlap between the signatures) less than −80. With the limma disease-associated gene signature, we identified 15 FDA-approved candidates for GBM, 12 candidates for LIHC, 92 candidates for LUAD, and 39 candidates for PAAD. With the DESeq2 disease-associated gene signature, we identified 13 FDA-approved candidates for GBM, 6 candidates for LIHC, 56 candidates for LUAD, and 30 candidates for PAAD. With the transfer learning disease-associated gene signature, we prioritized 18 FDA-approved candidates for GBM, 11 candidates for LIHC, 95 candidates for LUAD, and 29 candidates for PAAD (**Figure 3 & Supplemental Figure 16-19**).

**Figure 3:**
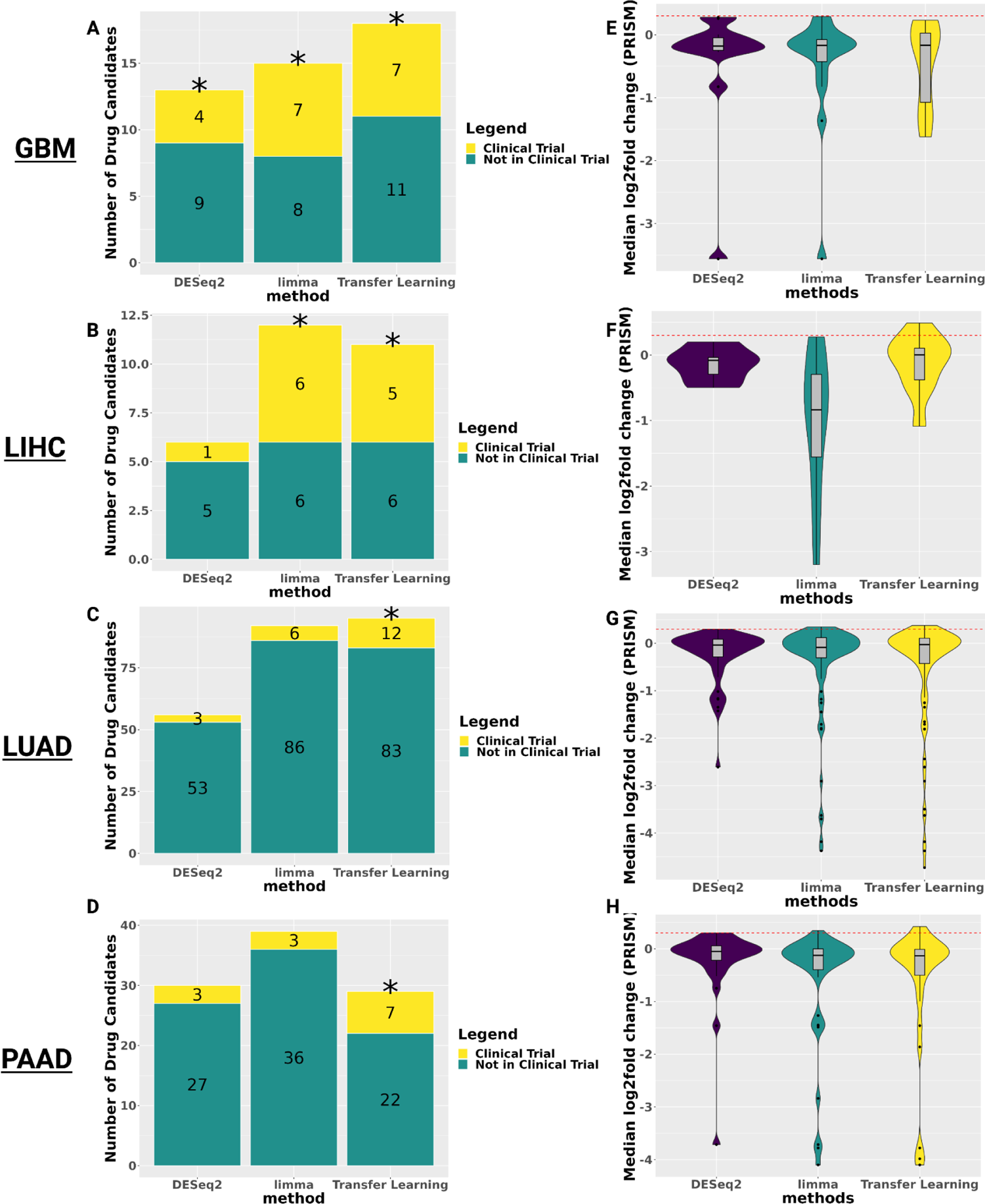
Performance of drug candidates predicated by signature reversion for each disease-associated gene signatures. The number of predicted drugs annotated by whether it has been in a clinical trial for **A)** GBM, **B)** LIHC, **C)** LUAD, or **D)** PAAD, respectively. Violin plot of the median log2 fold change of cancer-specific cell lines for candidates identified by the disease-associated gene signatures where the red horizontal line indicates the 0.3 log fold change threshold indicating cell line sensitivity to the drug candidates in the PRISM primary screen for E) GBM, F) LIHC, G) LUAD, and H) PAAD.

**Figure 4:**
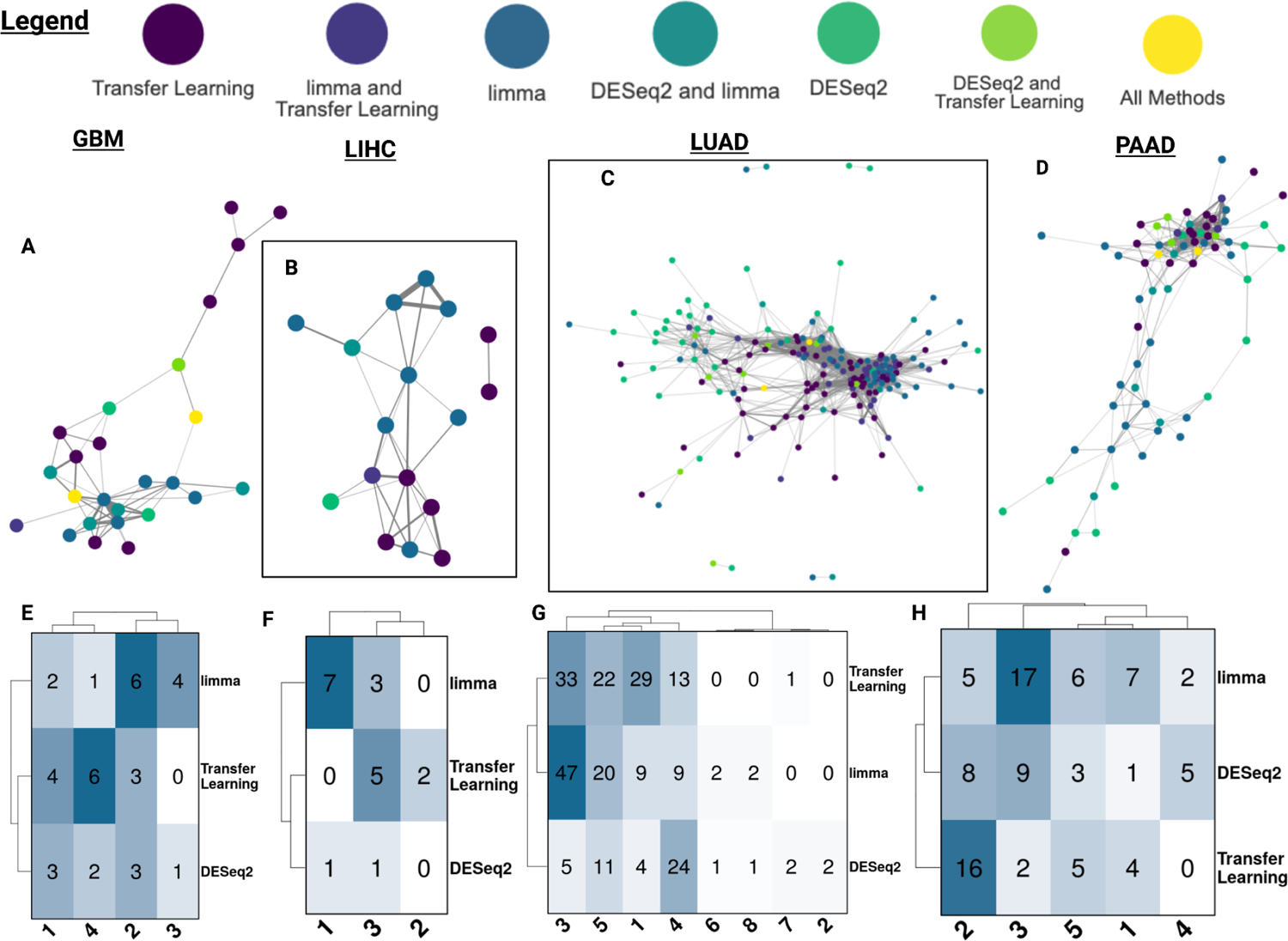
Drug-drug similarity network analysis for prioritized candidates. **A-D)** Drug-drug similarity networks based on cosine similarity of LINCS perturbation profiles of candidates, where each node is a candidate colored by the disease-associated gene signature used to identify that candidate, and the top 90% of edges are displayed and weighted by cosine similarity for **A)** GBM candidates (GI1 profiles), **B)** LIHC cancer candidates (HEPG2 profiles), **C)** LUAD candidates (A529 LINCS profiles), and **D)** PAAD candidates (YPAC profiles). Heatmaps of the composition of the Leiden communities in the drug-drug similarity networks for **E)** GBM candidates (GI1 profiles), **F)** LIHC candidates (HEPG2 profiles), **G)** LUAD candidates (A529 LINCS profiles), and **H)** PAAD candidates (YPAC profiles).

We next assessed if the drug repurposing candidates identified for each disease-associated signature and cancer have been in clinical trials or assessed by PRISM (a pooled drug screen of 930 cancer cell lines treated with 21,000 drugs to identify which inhibit cancer growth) for that cancer (**Figure 3** & **Supplemental Figure 16-19**). We performed permutation testing of a matched number of random sets of drugs for each as a comparison (**Supplemental File 2)**. The limma disease-associated signature drugs were enriched for drugs identified as sensitive by the PRISM drug screen for all four cancers but only enriched for drugs in clinical trials compared to the random sets for 2 of the 4 cancers (i.e., GBM and LIHC). Transfer learning disease-associated signature drugs were enriched for drugs in clinical trials compared to the random sets for all four cancers, but only enriched for drugs identified as sensitive by the PRISM drug screen in 2 of the 4 cancers (i.e., GBM and PAAD). Finally, DESeq2 disease-associated signature drugs were only significant compared to random sets of drugs in clinical trials for GBM. However, in the case of the PRISM drug screen, DESeq2 disease-associated signature drugs were enriched for sensitive drugs for all four cancers. Across the cancers, there were similar performances for the positive controls between the methods, but a lack of overlapping candidates (discussed further in the next section) suggests that each of the three disease-associated gene signature approaches identified different signatures resulting in different but plausible candidates.

### Comparison by cancer of prioritized drug targets and mechanisms of action

For each of the disease-associated gene signatures, most of the drug repurposing candidates identified through signature reversion were unique to a particular analysis: 50%, 79%, 61%, and 65% of the candidates were unique to disease-associated gene signature analysis in GBM, LIHC, LUAD, and PAAD, respectively. Between the sets of candidates per cancer, the maximum overlap between them was 3 in GBM (∼9% of all the candidates, **Supplemental Figure 20**). The mechanism of action (MOA) captured by each set also varied (**Supplemental Figure 21-24**). This might be due to broad MOA (e.g., tyrosine kinase inhibitors) or incomplete knowledge about drug MOA and targets [e.g, cabozantinib, which was known to inhibit MET proto-oncogene (MET) and vascular endothelial growth factor receptor 2 (VEGFR-2), but later found to also target RET proto-oncogene (RET)].^55^ Interestingly, each analysis found a diverse group of drugs with several different MOAs. GBM and LUAD had more MOA overlap compared to LIHC and PAAD. In LUAD, we identified drug repurposing candidates from all three disease-associated gene signature reversion results with the glucocorticoid receptor agonist MOA. This MOA has been shown to induce cell dormancy and stop the growth of cell lines.^56^ In LIHC, we found mTOR inhibitors in both the limma and transfer learning disease-associated gene signature reversion candidates. These inhibitors have been previously tested in the treatment of liver cancer and are thought to impact several liver cancer cellular phenotypes such as inflammation, angiogenesis, and metabolism.^57^ Lastly, in PAAD, the drug formoterol (a β adrenergic receptor antagonist identified by DESeq2 and transfer learning disease-associated gene signatures) and phentolamine (a ɑ-adrenergic receptor antagonist identified by limma disease-associated gene signature) share a common MOA, adrenergic receptor antagonist. Within the literature, ɑ-adrenergic receptor antagonists have been shown to affect apoptosis and proliferation^58^, and β-adrenergic receptor antagonists have suppressed cell invasion.^59^

We also investigated the overlap of the drug targets annotated from LINCS, DrugBank, CLUE, and STITCH databases for the identified candidates and found less than 15% overlapped across all three analyses for a cancer (**Supplemental Figure 25**). GBM had the highest number of overlapping drug targets (∼14.7% of drug targets), while LIHC had the lowest (∼5% of drug targets). As expected, several of these shared drug targets for each cancer are cytochrome P450 (CYP450) enzymes, essential for drug metabolism (**Supplemental Figure 26-28**).^60^ In addition, to determine if the identified candidate set differences between the three analyses within a cancer were due to cut-offs, we conducted a Spearman’s correlation between the normalized connectivity score (NCS) and false discovery rate (FDR) for each candidate in each analysis within the four cancers (**Supplemental Figure 30-34**). These correlations (NCS range:-0.027 to 0.59; FDR range:-0.12 to 0.38) suggest that the disease-associated gene signature reversion analyses (i.e., DESeq2, limma, and Transfer Learning) result in different rankings of drug candidates (**Supplemental Figure 34**). Overall, signature reversion of disease-associated gene signatures from three different approaches identified sets of drug candidates with different MOAs and drug targets.

**Figure 5:**
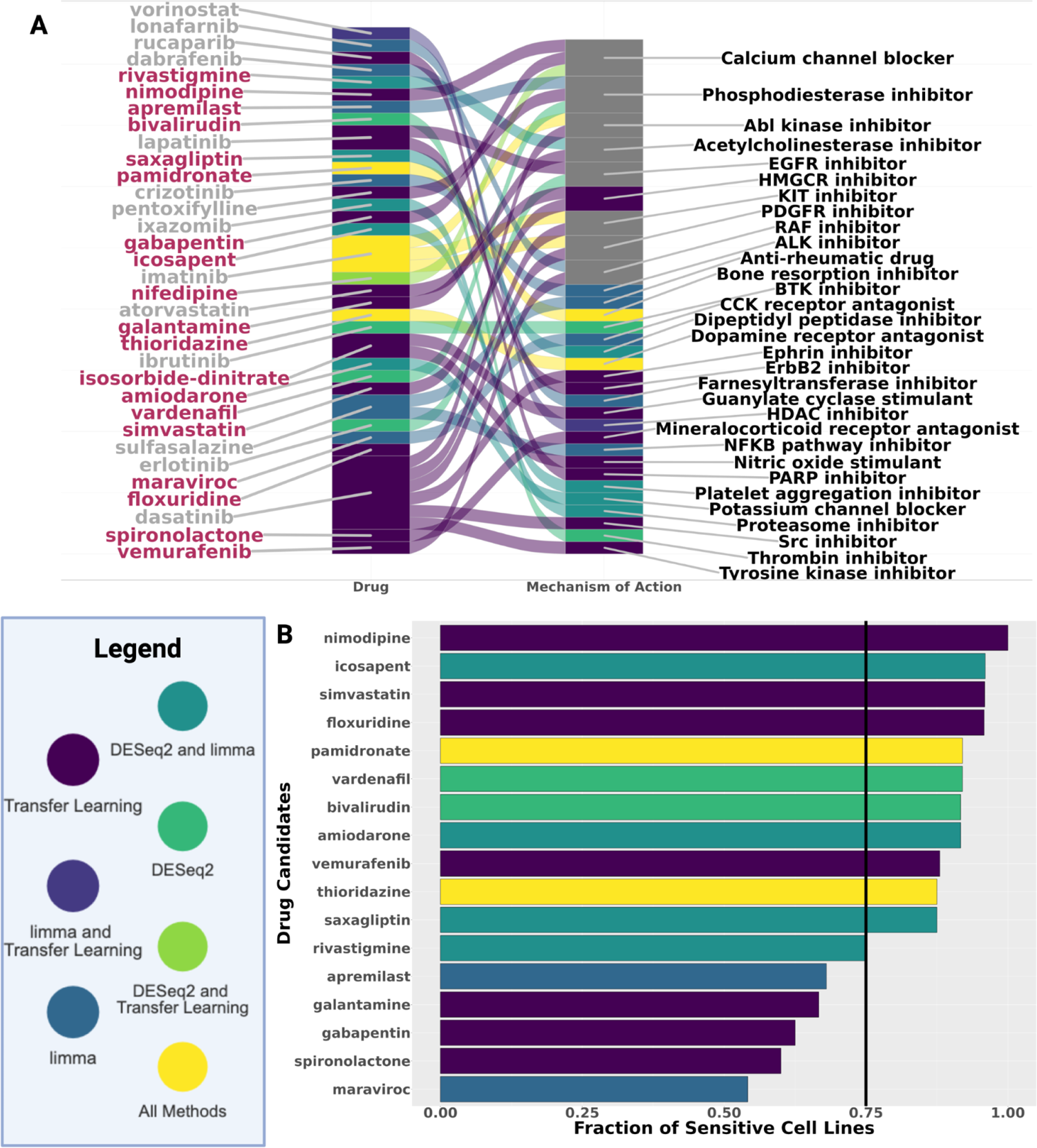
Prioritized GBM drug repurposing candidates perturb multiple MOAs and impact GBM cell line growth. **A)** Alluvial plot of the mechanism of action for the drug candidates from DESeq2, limma, and transfer learning disease-associated gene signature reversion analyses with drugs ordered by the median rank across analyses and MOAs ordered by the number of candidates with that MOA and then by alphabetical order (gray = candidates that have been in a clinical trial for GBM, maroon = candidates have not been in a clinical trial for GBM). **B)** The fraction of sensitive cell lines based on the PRISM primary screen for the top drug repurposing candidates that were not in a previous clinical trial for GBM. Black line at 0.75 indicates the fraction of GBM cell lines sensitive to the standard treatment, TMZ. For both plots, drug candidates are colored by the disease-associated gene signature used to identify it as a candidate.

Next, we wanted to determine if the candidates identified through signature reversion of different disease-associated gene signatures perturb the same genes. We used the perturbation profiles (LINCS level 5 modified z-scores) from the cancer-specific cell lines (the same ones used in the signature reversion step of our study), to determine the gene expression changes induced by each drug repurposing candidate. We applied cosine similarity to these perturbation profiles between candidates and constructed a drug-drug perturbation similarity network for each cancer. These drug-drug networks allowed us to compare and evaluate the different perturbation profiles (i.e., the gene expression changes before and after treatment) between the various candidates. In our analysis, we found drug repurposing candidates for each cancer generally clustered by the disease-associated gene signature approach used to identify them (**Figure 4A-D**). This indicates that repurposing candidates identified from the same disease-associated gene signature reversion analysis tend to perturb more similar gene sets than candidates identified from other disease-associated gene signature reversion analyses. Thus, they are not perturbing the same genes across the three analyses. To further support this observation, we determined Leiden communities for each cancer drug-drug network and compared the composition of these communities (**Figure 4E-H**). Drugs in the same Leiden community are more connected and thus more similar to each other compared to candidates in other communities. In GBM, which had the largest overlap of drug candidates across analyses, there were distinct communities with a higher number of candidates from one analysis, such as community 4 (transfer learning) and community 3 (limma) (**Figure 4E**). These distinct communities with a large portion of candidates from one analysis were also seen in the LIHC, LUAD, and PAAD drug-drug networks (**Figure 4F-H**). Across the different cancers, this highlights that drug candidate cell line gene expression perturbation profiles differ between the three disease-associated gene signature reversion candidate sets.

Because previous studies show that drugs with similar structures tend to have similar efficacy and safety profiles^61^, we also created drug-drug similarity networks based on drug structures using the Tanimoto coefficient between drug structures (**Supplemental Figure 35**). We found a similar trend as the drug-drug perturbation similarity network that repurposing candidates identified by signature reversion of the same disease-associated gene signature clustered closer to each other than repurposing candidates identified by signature reversion of the other disease-associated gene signatures. However, these structure similarity network clusters had more candidates from different disease-associated gene signature candidate sets in the same communities than the drug-drug perturbation similarity networks (**Supplemental Figure 35**). This might be because several candidates did not have available drug structures (i.e., SMILES structures from customCMPdb R package LINCS data set). Overall, both the structure similarity and perturbation drug-drug networks further support that the candidates identified by each disease-associated gene signature approach differ. Combined with our findings that drug candidates identified by reversion of each of the three disease-associated gene signatures are similarly represented in clinical trials and drug responses in the PRISM cell lines, these disease-associated signature methods identified disparate, but potentially similarly effective drug repurposing candidates.

### Additional validation of prioritized GBM drug repurposing candidates

We further prioritized the validation of repurposing candidates for GBM because it is the rarest of the selected cancers studied here and has only one standard of care chemotherapy, temozolomide.^62, 63^ In addition, our prioritized drug candidates included more drugs in GBM clinical trials and more GBM cell lines sensitive to treatment in the PRISM drug screen than selecting random drugs. This cancer also had the most overlap between candidates (3 out of 32) across the three disease-associated gene signature reversion analyses (**Figure 5A**). We first filtered for drug repurposing candidates that have not been in a clinical trial for GBM and have more PRISM cell lines sensitive to treatment with our candidate than temozolomide (**Figure 5B**). This resulted in 11 candidates from the three disease-associated gene signature reversion results (2 from DESeq2, and 4 from transfer learning, including 3 shared between DESeq2 and limma and 2 shared between all three). From those, simvastatin, floxuridine, vardenafil, amiodarone, and thioridazine had already been investigated for the treatment of GBM with *in vitro* and/or *in vivo* studies.^64–70^ We did not further consider bivalirudin because there was evidence in DrugBank that this drug was unlikely to pass the blood-brain barrier. Vemurafenib was also excluded from further testing because of a basket clinical trial for gliomas with BRAFV600 mutation.^71, 72^ Therefore, we had 4 remaining candidates: pamidronate, nimodipine, icosapent, and saxagliptin. We found compelling evidence for each of these with respect to their drug targets, perturbed pathways, and other *in vitro* drug testing.^73–76^ Thus, we conducted cell viability experiments to determine if each of the four candidates was capable of decreasing the growth of the U251 cell line, a GBM cell line not included in the PRISM drug screen. In an initial screen of U251 cells with concentrations ranging from 1-100uM, pamidronate and nimodipine caused a decrease in cell viability. Icosapent and Saxagliptin were tested once with three technical replicates and did not show a decrease in cell growth (**Supplemental Figures 36 & 37).** We therefore selected pamidronate and nimodipine for further evaluation.

For both pamidronate and nimodipine, we conducted two independent cell growth drug screens of GBM cells propagated under brain tumor initiating cell conditions from either a standard cell line (U251) or a patient derived xenograft (JX39) (**Figure 6 A-D & Supplemental Figure 38**). WIn our study, we found that 40uM pamidronate was always sufficient to significantly inhibit the growth of GBM U251 and JX39of GBM cells propagated under brain tumor initiating cell conditions from either a standard cell line (U251) or a patient derived xenograft (JX39) (**Figure 6 A–B, & Supplemental Figure 38**). As mentioned previously, while pamidronate has been tested in cell line systems, to our knowledge, these are novel findings indicating GBM PDX-derived cells were sensitive to pamidronate. As PDX systems have been shown to better reflect patient molecular profiles and phenotypes like drug response than cell lines due to the genetic drift of cell lines during passaging and long-term growth in vitro environments ^77, 78^, this lends further support to pamidronate as a prioritized GBM drug repurposing candidate. For nimodipine In our cell growth assay, we found that 40uM nimodipine was consistently sufficient to significantly inhibited the growth of both U251 and JX39 GBM cells across all three concentrations we tested (i.e., 20 uM, 30uM, 40 uM) (**Figure 6C, D, & Supplemental Figure 38**).

**Figure 6:**
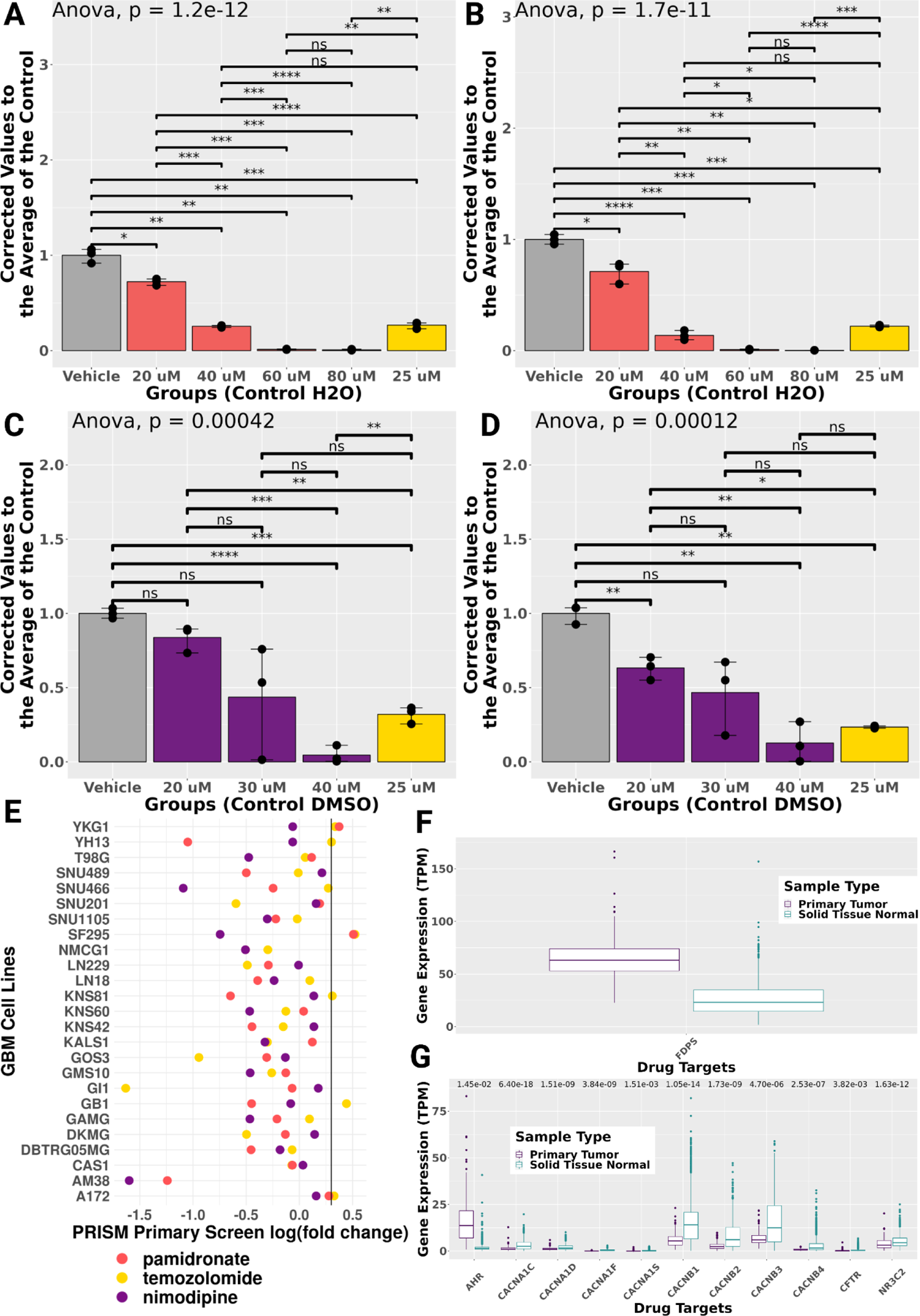
Pamidronate and nimodipine inhibit GBM cell growth in U251 and JX39 PDX in vitro models. CellTiter-Glo growth assay results for pamidronate in **A)** U251 and **B)** JX39, respectively. CellTiter-Glo growth assay results for nimodipine in **C)** U251 and **D)** JX39, respectively. The yellow bar is the results from temozolomide treated cells. *P < 0.05, **P < 0.01, ***P < 0.001, ****P < 0.0001 ANOVA with Bonferroni corrected t-tests for pairwise comparisons. **E)** Dot plot of PRISM primary screen results by GBM cell lines. Boxplot of drug target expression in tumor tissue and control brain tissue for **F)** pamidronate and **G)** nimodipine, respectively. Significant p-values from DESeq2 are plotted above the drug target expression.

Pamidronate, one of the top candidates for all the methods (ranked via theNCS as 2nd, 5th, and 10th from the DESeq2, transfer learning, and limma disease-associated gene signatures, respectively), is a bone resorption inhibitor used for the treatment of moderate to severe hypercalcemia of malignancy, Paget’s disease, osteolytic metastases, and osteoporosis (**Figure 5A**).^72, 79^ Pamidronate induces apoptosis of hematopoietic tumor cells by inhibiting farnesyl diphosphate and geranylgeranyl diphosphate.^72^ First, we compared the treatment response of GBM cell lines treated with pamidronate to the standard of care, TMZ, in the PRISM cell line data set. While this dataset only contains cell line growth results and TMZ therapy is usually combined with radiotherapy in clinic, we found that 12 of the 24 GBM cell lines treated with both drugs had a lower log2 fold change in cell growth with the pamidronate treatment, and across all GBM cell lines (n=25) only 2 were identified as not sensitive to pamidronate treatment (log2 fold change >0.3) (**Figure 6E**). Further functional enrichment analysis of the GI1 GBM cell line gene expression perturbation profile used in our signature reversion analysis identified several pathways associated with GBM. This included pathways associated with WNT signaling, development, and proliferation in the upregulated gene set, while acute inflammatory response and metabolism to fat-soluble and retinoids were enriched in the downregulated gene set (**Supplemental File 3**). In addition, while not identified as significantly differentially expressed by DESeq2 or limma, we observed increased gene expression of pamidronate’s known drug target, *FDPS*, in primary tumor tissue (**Figure 6F**). This suggests that pamidronate might particularly target tumor tissue itself. *FDPS* codes for an enzyme that has been implicated in the stem-cell-like characteristics of GBM that have been previously associated with GBM treatment resistance.^80^ Pamidronate has also been previously shown to enhance paclitaxel-induced apoptotic death in the U87MG GBM cell line.^81^

The other candidate, nimodipine, was the highest-ranking drug repurposing candidate from the transfer learning methodology for GBM that was not in a previous GBM clinical trial. This calcium channel blocker is used as an adjunct to improve neurologic outcomes following subarachnoid hemorrhage from a ruptured intracranial berry aneurysm (**Figure 5A**).^72^ In previous studies, this drug was shown to be more effective in causing cancer cell death in combination with other anticancer drugs.^82, 83^ In the PRISM drug screen, we found that 16 of the 24 GBM cell lines treated with both nimodipine and TMZ were more sensitive to nimodipine (**Figure 6E**). Additionally, this drug screen found all the GBM cell lines tested sensitive to nimodipine, while only 75% of the cell lines were sensitive to TMZ. Furthermore, our DESeq2 differential gene expression analysis confirms that all 11 of the drug targets of nimodipine are differentially expressed at the gene level between GBM and control brain tissue samples (**Figure 6G**). The functional enrichment analysis of the GI1 perturbation profile of this drug further suggests that it perturbs pathways associated with metabolism, cellular processes, and the cytoskeleton (**Supplemental File 3**).Furthermore, we also conducted one cell viability experiment with an NHA astrocyte cell line after treatment with either pamidronate or nimodipine (**Supplemental Figure 39**). As both candidates inhibited cell line growth in this one experiment, this underscores the need for further evaluation to determine both effectiveness and safety. However, our study highlights novel drug repurposing candidates for the treatment of GBM prioritized with the methods described here.

## Discussion

While previous cancer studies have applied signature reversion to identify drug repurposing candidates^11^, we sought to investigate how different approaches to developing the disease-associated gene signature for signature reversion impacts downstream drug candidate selection. In four low-survival cancers, we demonstrate that signature reversion identified drug repurposing candidates enriched for drugs in clinical trials or performed well in PRISM’s cancer cell line drug screen.^84, 85^ The candidates identified from the two differential expression (DESeq2 and limma) and transfer learning (MultiPLIER) disease-associated signature approaches were not commonly shared, and candidates prioritized from the same disease-associated gene signature were more similar to each other than candidates identified with other disease-associated gene signature approaches. This observation indicates that these disease-associated gene signature approaches identify unique biology that researchers can further leverage to prioritize drug repurposing candidates. Our deeper investigation of the disease-associated gene signatures underscored how different the disease-associated gene signature methods are by determining the gene composition and pathway enrichment of the disease-associated gene signatures. Overall, the main finding of our study is that researchers should apply these different disease-associated gene signature approaches in tandem to increase the number of prioritized drug repurposing candidates for downstream *in vitro* and *in vivo* testing.

In addition to being among the lowest survival cancers, some of the selected cancers also have unique disease features or data availability that we leveraged in our study. As previously mentioned, GBM is the rarest of the cancers in this study, which limited data availability, while the more common LUAD had the most drug candidates tested in a cancer-specific cell line in the LINCS database (e.g., n= 6,742 for the A529 LUAD cell line compared to 685 candidates for the GI1 GBM cell line). PAAD bulk tumor tissue RNA-seq profiles are known to be more similar to non-tissue profiles than most cancers because PAAD tumors are mostly composed of cancer-associated fibroblasts and a small population of cancer cells.^86^ Our principal component analysis highlights this: the pancreatic tumor samples cluster with non-cancerous pancreatic tissue samples. Despite the influence these factors may have on the disease-associated gene signatures, signature reversion, and the subsequent prioritized repurposing candidates, we find that most of the prioritized candidates had been in clinical trials and/or performed well in PRISM drug screen. Therefore, we are optimistic that the approaches we present here are robust and that our prioritized candidates warrant further investigation by the research community.

Many prior studies support the repurposing of the drug candidates we identified. Itraconazole, an antifungal medication we predicted from the LIHC transfer learning disease-associated signature, has been shown to inhibit LIHC cell growth and promote apoptosis via several pathways (e.g., WNT, ROS, AKT/mTOR/S6K) and other death receptor pathways.^87^ In another example, from the limma disease-associated gene signature for LUAD we prioritized cladribine, a chemotherapy indicated for leukemia. There is prior evidence that the A529 cell line (the reference cell line we used for our signature reversion analysis) was sensitive to cladribine and that this drug enhanced apoptotic cell death due to overexpression of DNase γ.^88^ While this might indicate that specifically, the A529 cell line might be sensitive to this candidate, all of the other PRISM LUAD cell lines were also sensitive to cladribine. Additionally, we predicted albendazole, an antihelminthic used to treat neurocysticercosis (an infection of the nervous system caused by pork tapeworms) and other worm infections, via the DESeq2 disease-associated gene signature for the treatment of PAAD. This drug has been shown to induce apoptosis and reduce proliferation and migration in pancreatic cell lines and an in vivo nude mouse xenograft model.^89^ As expected, many candidates we identified are currently approved for another type of cancer (e.g., cladribine is used for leukemia treatment but predicted here for application in lung cancer). However, we also found many repurposing candidates currently approved for conditions that are not neoplasms (e.g., nimodipine candidates for GBM). The PRISM study also observed the identification of non-neoplastic drugs for cancer applications.^84^ In addition, some drugs have drug targets for different organisms, such as fungi and tapeworms (i.e., itraconazole and albendazole). This observation suggests that these drugs have off-target or cancer context-specific drug mechanisms that are not entirely understood, a potentially interesting research direction.

We also demonstrated the potential use of transfer learning for identifying a disease-associated gene signature for signature reversion. Transfer learning has previously been used to identify disease-associated latent variables in several rare disease applications, including medulloblastoma, a rare brain tumor, and in other drug repurposing studies.^90, 91^ Here, we compared its performance to other disease-associated gene signature methods across multiple cancers. Overall, the transfer learning disease-associated gene signature approach prioritized candidates in all four cancers that were either previously in clinical trials or demonstrated cancer cell line sensitivity in the PRISM database. Additionally, the top two predicted GBM drug repurposing candidates inhibited GBM cell growth in the in vitro GBM cell culture systems we tested. This approach may also be beneficial for identifying drug repurposing candidates for other heterogeneous complex diseases.

This study addressed the critical question in signature reversion drug repurposing of how different disease-associated gene signature approaches impact downstream drug repurposing candidate selection for four low-survival cancers. Here we demonstrate the utility of transfer learning-based disease-associated gene signature approaches to signature reversion and provide prioritized candidates for four of the lowest survival cancers. By investigating the predicted drug candidates and the disease-associated gene signatures for each approach, we found that each disease-associated signature approach identified unique genes, pathways, and drug repurposing candidates. We validated our candidates using publicly available cancer cell line drug screen and clinical trial data and further validated top GBM drug repurposing candidates in cell line and xenograft systems. We also provide a valuable resource of prioritized candidates for future efficacy and safety studies. In conclusion, our results underscore how using each disease-associated gene signature in tandem may identify additional prioritized drug repurposing candidates for further investigation.

### Limtations of the Study

While overall, we did not find that signature reversion of one disease-associated gene signature approach outperforms the others, we did observe a specific challenge for developing a disease-associated gene signature via the DESeq2 methodology. To generate the DESeq2 disease-associated gene signatures, we used an absolute shrinkage log2 fold change cut-off to identify a list of ∼100 genes. For the LIHC and LUAD data sets, we observed that with DESeq2, there was more variance between 0 and the highest positive fold change compared to the variance between the lowest fold change and 0 for differential gene expression (**Figure 1C & D**). This resulted in a disease-associated gene signature that contained predominantly upregulated genes in cancer compared to non-tumor tissue. This may have resulted in the observed lower performance of the DESeq2 disease-associated gene signature drug repurposing candidates.

In addition, we focused on cancer-specific cell line perturbation profiles for each cancer as multiple studies, including the LINCS project data release, found perturbation profiles of the same cell line with different drugs had more similarity than the same drug across different cell lines.^11, 54^ This influence of the molecular context of cell lines on drug response is important because it affects the prediction of drug repurposing candidates and our orthogonal validation analyses. For example, we hypothesize that the drug structure drug-drug similarity networks did not cluster the candidates the same as the drug perturbation drug-drug similarity network analysis due to the effect of cell line context on the perturbation profiles. In addition, while the literature provides supporting evidence for our results, as evidenced in the above paragraph, additional context may be important. For example, EGFR has been a drug target for many therapies in clinical trials for GBM, including those we predicted by reversion of the transfer learning disease-associated gene signature, erlotinib and lapatinib.^92^ However, as many EGFR inhibitors have failed in GBM clinical trials, EGFR inhibition alone may not be a suitable target for GBM.^92^ However, there is evidence that EGFR inhibitors might be therapeutic for some GBM patients and not others, underscoring the importance of disease heterogeneity and personalized medicine efforts to pair the right drug with the right patient at the right time. In addition, within GBM tumors, some tumor cells are sensitive to EGFR inhibitors, while some cells are not.^92^ Thus, approaches for determining patient treatment by tumor subtypes or novel drug design to target multiple mechanisms across clonal populations are needed to combat tumor heterogeneity.^92^ Ideally, future drug repurposing studies will be powered to segregate tumor profiles and LINCS-like perturbation profiles to match these different subtypes or biomarkers for predicting therapies as we were limited here by the cell lines in the LINCS dataset, the dosing protocol (i.e., 24-hour and 10uM dosing protocols), and single-agent studies. These challenges highlight the most critical limitation of signature reversion: dependence on context-specific perturbation signatures. Future development of perturbation signature resources and methods has the potential to improve this approach.

## Methods

### Key resources table

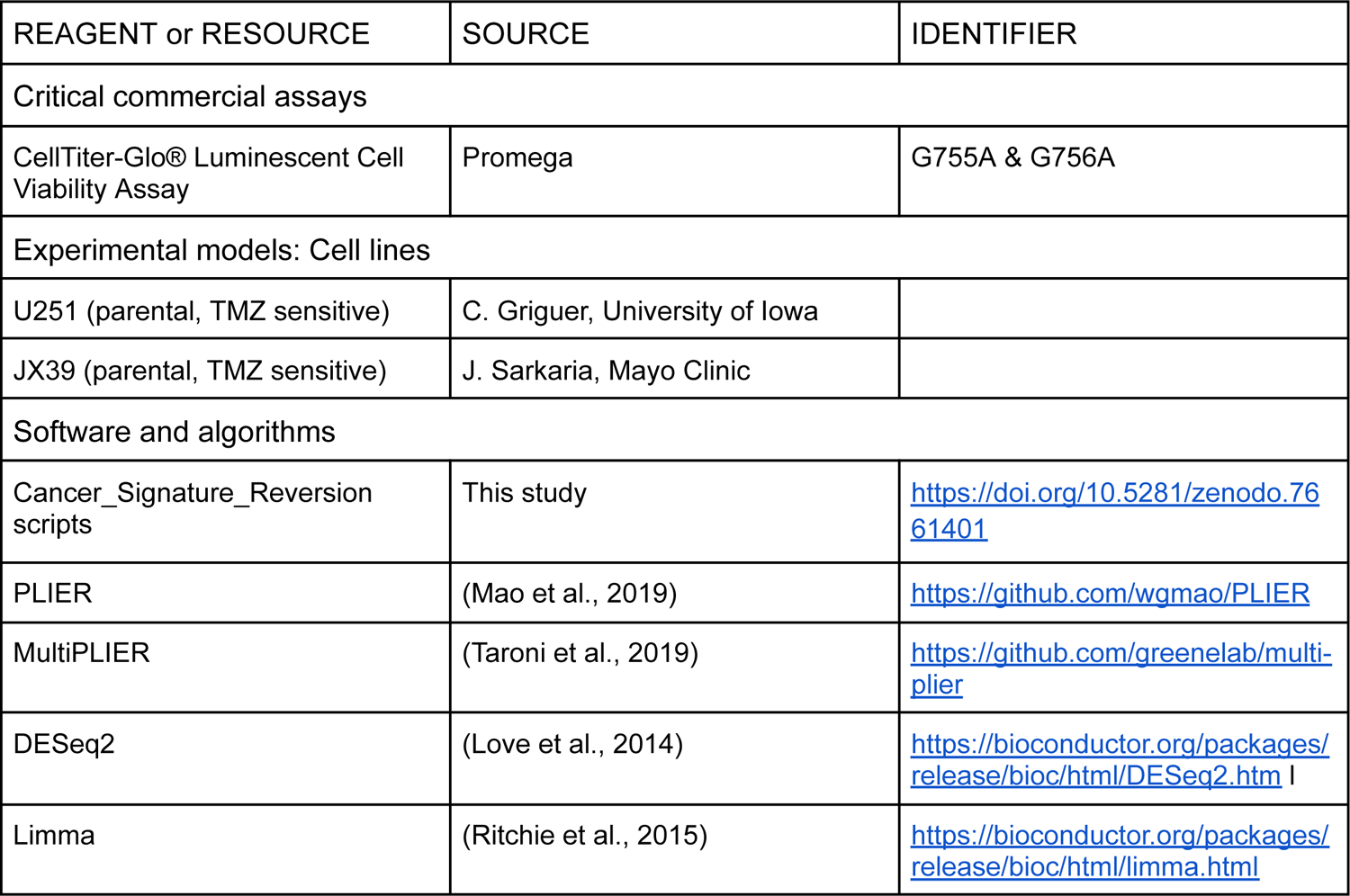
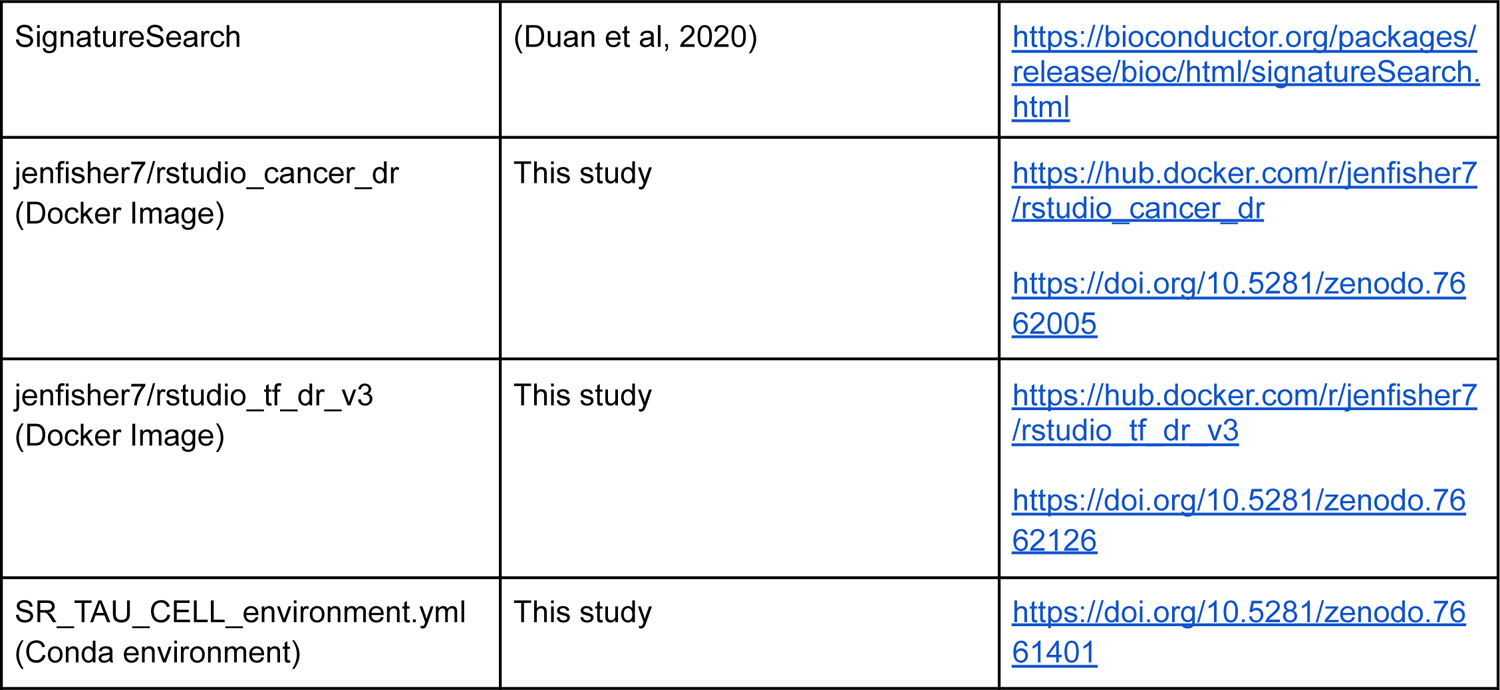

#### Scripts, Dockers, and Conda environment

The scripts for this project are available on Zenodo at https://doi.org/10.5281/zenodo.7661401. In addition to the scripts here, the Docker images used for this analysis are publicly available on Docker Hub (jenfisher7/rstudio_tf_dr_v3 & jenfisher7/rstudio_cancer_dr) and Zenodo (https://doi.org/10.5281/zenodo.7662126 & https://doi.org/10.5281/zenodo.7662005). For the Tau calculations, a conda environment was used (SR_TAU_CELL_environment.yml). Detailed computer and package version information are included in **Supplemental File 4**.

#### Data download

For this study, we downloaded RNA-Seq count profiles from the Recount3 database using the Recount3 R package.^93^ This database contains RNA-Seq samples from tumor samples via The Cancer Genome Altas (TCGA) and control samples from TCGA and the Genotype-Tissue Expression (GTEx) project (Access date for GBM/BRAIN, LIHC/LIVER, LUAD/LUNG, and PAAD/PANCREAS Recount3 project, respectively: Dec. 2021, May 2022, May 2022, and May 2022). We used the rest of the human samples from the Sequence Read Archive (SRA) that were also processed and stored by Recount3 for the dimension reduction step for transfer learning (n = 316,443) (Access date: Dec. 2021). The numbers of tumor and control non-tumor samples for each cancer are listed in **Supplemental File 1**. For the GBM comparison, we filtered the TCGA samples to include only IDH-wildtype due to the reclassification of brain tumors in 2021.^94^ In addition, because GBM can originate from almost any brain region, we also used all GTEx samples from any brain region.^95^ A limitation of this is that the brain regions are known to vary in gene expression. Unfortunately, the GBM TCGA data does not include information on tumor location in their database; therefore, we could not use batch correction or regression to reduce the influence of the brain region for the GBM analyses.

The Library of Integrated Network-based Cellular Signatures (LINCS) program created a database of gene expression profiles of cell lines before and after being exposed to perturbing agents such as small molecules (1,131 cell lines and 41,847 drugs).^54^ We downloaded the Expanded CMAP LINCS Resource 2020’s modified z-score (level 5) for compound data (aka. small molecules) from https://clue.io/data/CMap2020#LINCS2020 (Access date: Jan. 2022). The LINCS level 5 data has been uniformly processed to control for plate variation and had technical replicates combined. In addition, we accessed the metadata for the compounds, genes, cell lines, and signatures. For this study, we used signatures for the 10 uM at 24 hours because it has more drug signatures than other concentrations and time points. In addition, for this study, we focused on the perturbation profiles of cancer cell lines derived from the particular cancer we are studying (i.e., GI1 for GBM, HEPG2 for LIHC, A529 for LUAD, and YAPC for PAAD).

We downloaded the Drugs@FDA database in March 2022 from https://www.fda.gov/drugs/drug-approvals-and-databases/drugsfda-data-files (Accessed date: March. 2022). From the marketing status and products tables, we generated a table including the method of drug delivery, dosage, active ingredients, and FDA approval (fda_product_info_df.csv). In addition, we used ClinicalTrials.gov to determine drugs in clinical trials for each of the different cancers (accessed date: GBM-Feb. 2022; LUAD-June 2022; PAAD-June 2022; LIHC-June 2022). For each cancer, we searched by cancer name to collect all of the clinical trial information.

The Profiling Relative Inhibition Simultaneously in Mixtures (PRISM) project conducted a pooled drug screen that treated 930 cancer cell lines with 21,000 drugs to identify which inhibit cancer growth.^84^ We downloaded the primary and secondary screen data from the Cancer Dependency Map (DepMap) portal (Accessed date: Feb. 2022).^96^ We limited the analysis to a single cell line per cancer type since previous studies have shown that perturbation profiles of the same cell line across treatments were more similar than those derived from applying the same drug across cell lines.^11, 54^

We downloaded a human protein-protein interaction network from the STRING database (accessed date: March 2022; version 11.5).^29^ This database contains known and predicted protein-protein interactions, including direct and indirect associations.^29^ We divided the protein-protein interactions in the network by 1000 and filtered for interactions greater than 0.7, as a score greater than 0.7 indicates high confidence in that protein-protein interaction. We converted the protein Ensembl ID to the HGNC gene symbol for each protein using the provided STRING annotations. Additionally, we accessed the drug structure data files for drug candidates via customCMPdb’s loadSDFwithName function for the LINCS dataset (Accessed date: July. 2022).^97^

#### Principal component analysis

We conducted principal component analysis (PCA) for each of the selected cancers and the GTEx control tissue. We then used the variance stabilizing transformation (VST) from DESeq2 to transform the raw count data from Recount3 data (download described in the previous section)^22^ and used it for the PCA with the prcomp function in base R. The PC plots were visually inspected for clustering of samples based on sex, age, RIN score, and ischemic time. While the GTEx data did vary according to ischemic time and RIN score, as the TCGA dataset did not contain this information, they were not included as covariates in subsequent models.

### Transfer learning for disease-associated gene signature identification

#### Dimension reduction and transfer learning

We normalized the gene expression profiles from the Recount3 database as transcript per million (TPM) normalized for the transfer learning approach. To decompose the TPM normalized gene expression profiles from Recount3 into latent variables, we applied the dimension reduction method Pathway-Level Information Extractor (PLIER).^23, 98^ We removed the TCGA and GTEx samples from the downloaded Recount3 database (n = 316,443 samples remaining) to remove their influence on the resulting latent variables. This resulted in 385 latent variables. The PLIER algorithm dimension reduction approach features the L1 and L2 penalties in the loss function resulting in 1) latent variables reflective of annotated pathways (e.g., from Reactome) and 2) most genes having a zero weight (i.e., only a fraction of genes have a high and significant gene weight).^98^ We then transferred these learned latent variables from Recount3 to the TPM normalized gene expression profiles in TCGA and GTEx via matrix multiplication using MultiPLIER ^23^.

#### Differential latent variable analysis

We wanted to validate the transfer of information via the gene labels of the transfer learning approach. We hypothesized that if the gene labels transfer information and the gene labels were switched, the transfer learning approach would not be able to identify significant latent variables. To test this hypothesis, we compared the GTEx brain frontal cortex and cerebellar hemisphere region samples from Recount3 by switching a fraction of gene labels starting at 10 up to 100% at 10% intervals. We performed each label switch step 50 times. We then calculated the average number of latent variables significant for each interval (adj. p-value < 0.05; fold change of 0.05) (**Supplemental Figure 40**) and used a linear regression model to determine if the percent of gene labels switched influenced the number of latent variables (R^2^ = 0.906 & p-value = 1.381e-05). This confirmed that when gene labels are randomly switched, the number of significant latent variables from the transfer learning approach decreases. This analysis indicates that information about the genes is indeed transferred in transfer learning.

To determine the disease-associated gene signature for the signature reversion drug repurposing, we conducted a differential latent variable analysis via latent variable-wise linear regression models and identified which latent variables are different between disease and control (limma).^19^ As we used data from TCGA and GTEx for the tumor and control samples, we included the database (TCGA/GTEx) as a covariate to reduce the influence of database-specific technical noise. We applied a Bonferroni-adjusted p-value of <0.05 as a cutoff for whether a latent variable was significant. In addition, we used the absolute fold change of approximately 3 standard deviations away from the mean to identify the most different latent variables between tumor and non-tumor samples for each cancer. For example, if 3 standard deviation from the mean resulted in an upper cut-off of 0.27 and a lower cut-off of −0.23, an absolute cut-off of 0.25 was used. Then, the genes that contribute the most to the latent variables (i.e., had the highest weights in the latent variable linear gene expression equation) for each cancer were identified. The number of top genes for each cancer varied based on how many of those genes overlapped with genes in other latent variables and the genes assayed in LINCS. The target disease-associated gene signature had approximately 100 genes. **Table 1** notes the total genes for each cancer. Finally, we used the gene expression fold change from the DESeq2 analysis between disease and control tissues to determine if the genes were up or down-regulated in the disease.^20^ These up and down-regulated genes were the disease-associated gene signatures in the gene expression signature reversion method.

**Table 1:** Disease-associated gene signatures across all the cancers. Table of the number of up-regulated (Up) and down-regulated (Down) genes included in the disease-associated gene expression signature.

| Method | GBM | PAAD | LUAD | LIHC |
| --- | --- | --- | --- | --- |
| DESeq2 | Up: 37<br>Down: 66 | Up: 68<br>Down: 33 | Up: 115<br>Down: 2 | Up: 115<br>Down: 6 |
| Limma | Up: 58<br>Down: 60 | Up: 85<br>Down: 26 | Up: 16<br>Down: 92 | Up: 49<br>Down: 57 |
| Transfer Learning | Up: 52<br>Down: 54 | Up: 25<br>Down: 95 | Up: 47<br>Down: 69 | Up: 62<br>Down: 26 |

### Differential gene expression analyses for disease-associated gene signature identification

#### Limma

For each of the cancers, we normalized the raw counts from Recount3 for GTEx and TCGA as transcript per million normalized and log-transformed (log(TPM+1)). To analyze differentially expressed genes between disease and control, our limma design matrix included the experimental groups and the originating database (i.e. GTEx or TCGA).^19^ We then used the Bonferroni method for multiple hypothesis testing. Based on the limma differential expression results, we selected genes with an adjusted p-value less than 0.05 and the highest absolute log2 fold change. However, for the disease-associated gene expression signature, we selected 90-120 genes with the highest and lowest log-fold changes (based on the study by Yang et al^11^) that overlapped with genes in the LINCS database. If the gene did not overlap, we removed it, and used the next gene in the highest or lowest fold change in the signature. The number of genes in the disease-associated gene signature for each method is listed in **Table 1**.

#### DESeq2

For each cancer, we used the raw counts from gene expression Recount3 for the GTEx and TCGA samples for the DESeq2 differential expression analysis identifying differential expressed genes between each cancer and non-disease sample set. The DESeq2 design formula also included the experimental groups and the database with other DESeq2 parameters at default values.^20^ Based on the DESeq2 results, we considered genes with an adjusted p-value less than 0.05 and highest absolute log2 fold change as significantly differentially expressed. Again, we used the top ∼90-120 highest and lowest log-fold changed genes for the disease-associated gene expression signature.

### Signature reversion

For this study, we used the enrichment approach applying a bi-directional weighted Kolmogorov-Smirnov statistic via the signatureSearch R package ^13^ because it has been shown to perform better at identifying similar drugs based on mechanism of action (MOA) and drug structure similarity than other enrichment approaches.^13^ The LINCS signature reversion method signatureSearch identifies drug candidates with a gene expression perturbation profile in LINCS most inverse to the disease profile (in this case, the disease-associated gene expression signatures).^13^ This approach ranks drug candidates by the normalized bi-directional weighted Kolmogorov-Smirnov statistic (to determine how similar disease and drug profiles are to each other) normalized by dividing the statistic by the signed mean of all the statistics from the subset of signatures corresponding to the same cell line and drug in the LINCS reference. We then filtered drug candidates based on the false discovery rate to correct for multiple hypothesis testing and on the Tau values that were determined based on the calculation conducted by the signatureSearch package.^13^ Tau values describe the uniqueness of the overlap between the perturbation signature and disease-associated gene signature. A Tau value of −100 means that the negative connectivity score (i.e., a more inverse signature) from the comparison between X perturbation signature and the disease-associated gene signature was more negative than that same X perturbation signature to other perturbation signatures in the reference database (here, the LINCS 2020 database). For example, the negative normalized connectivity score (NCS) from the signature reversion analysis for perturbation X is less than NCS scores from X perturbation signature to other perturbation signatures in the LINCS database. A Tau value of 100 indicates that the positive connectivity score (i.e., a closer matching signature) from the comparison between the perturbation and disease-associated gene signature was more positive than that same perturbation signature to other perturbation signatures in the LINCS 2020 database. Tau values close to 0 suggest that the overlap between disease-associated gene signatures and perturbation signatures is not a unique overlap for the perturbation. Overall, the Tau value helps identify drugs uniquely inverse to the disease-associated gene signature.^54^ In addition, we focused on the signature reversion results for the specific cancer cell line of interest (i.e., GI1 for GBM, HEPG2 for LIHC, A529 for LUAD, and YAPC for PAAD).

### Drug target analyses

The signatureSearch signature reversion results include information about the candidate’s drug targets. From this information, we plotted drug target gene expression (TPM) and the DESeq2 adjusted p-values to determine if there was a significant difference in gene expression of those targets between tumor and non-tumor control samples.^20^ In addition, we applied the signatureSearch R package drug set enrichment analysis (dsea_hyperG hypergeometric test^13^) to the FDA-approved drug candidates identified for each cancer with each disease-associated gene expression signature and their drug targets.

### Clinical trial evaluation

To evaluate if signature reversion of disease-associated gene expression signatures from a particular differential expression approach was able to identify more drug candidates that were already in clinical trials for a specific cancer, we performed permutation testing. We randomly selected the same number of FDA-approved drugs 1,000 as were identified as significant for each case, and determined what fraction of randomly selected drugs were in clinical trials for that specific cancer. We performed a one-tailed Wilcox test to determine if the fraction of FDA-approved drugs selected for each case was higher than for the randomly selected drugs.

### Drug candidate performance in the PRISM drug screen

We used the PRISM primary and secondary pooled drug screens to further evaluate if the identified candidates could reduce cell growth of cancer cell lines derived from the same cancer as the drug identified for repurposing.^84^ For the primary screen, PRISM has calculated the median of log fold change median fluorescence intensity between replicates of a cell line treated with a drug. They considered a cell line sensitive to a treatment if the median-collapsed fold-change was less than 0.3. We also assessed if a given approach for identifying differentially expressed genes for the disease-associated gene expression signatures for signature reversion results in a set of drug repurposing candidates with a larger fraction of candidates that cancer cell lines are sensitive to (for the specific cancer in question) than an equal size set of randomly selected drugs 10,000 times and then perform a one-tail Wilcox test.

### Comparing genes included in the disease-associated gene signature from different approaches

To investigate how different the log fold change and adjusted p-values from DESeq2 and limma were, we calculated Spearman correlations. DESeq2 uses the apeglm method for effect size shrinkage in their fold change calculations to reduce large estimates of log2 fold change caused by high variance or low expression of certain genes that are not representative of the true difference between disease and control.^99^ Limma, on the other hand, assumes that the input data is log2-transformed and then calculates the mean gene expression difference between disease and control by gene.^19^ Because of these differences in algorithms and statistical models within each approach, the log2 fold changes and adjusted p-values for each method are not the same. To compare transfer learning disease-associated signature genes with DESeq2 and limma disease-associated signature genes, we constructed volcano plots of the log-transformed fold change and the adjusted p-value for both DESeq2 and limma. On these plots, we labeled genes by the disease-associated gene signature they are a member of.

### Function enrichment of disease-associated gene signature genes

We used the R package gprofiler2 for functional enrichment analysis to determine pathways and other annotated gene sets enriched in the up-regulated and down-regulated gene sets used for signature reversion. ^42^ We used the Biological Process GO terms in a Gene Ontology semantic similarity analysis. In this analysis, we compared the enriched GO terms from all disease-associated gene signature sets for a cancer by semantic similarity based on the method proposed by Wang 2007.^100^ This approach considers the location of GO terms in the GO graph and their relationship to parent and children terms via the GOSemSim R package.^43, 100^ We plotted the results in a heatmap and clustered via the wald.D2 algorithm using ComplexHeatmap.^101^ We used functions from rrvgo (getGoTerm, loadOrgdb, getGoSize, reduceSimMatrix) to create GO term subgroups from the enriched pathways based on the parent term in the GO graph and the Wang semantic similarity.^102^

### Function enrichment analysis of LINCS profiles and annotation of top candidates

We analyzed the perturbation profiles (level 5 LINCS data) with gprofiler2 functional enrichment analysis to identify which pathways or transcription factors were perturbed by the drug candidates.^42^ For each drug, the gene cut-off for the functional enrichment analysis was the LINCS’s level 5 modified z-score greater than 2 for up-regulated gene set and less than −2 for the down-regulated gene set. These modified z-scores were calculated by the LINCS consortium by aggregating across the replicates within the level 4 data plate-control normalized Z scores. We investigated identified drug candidate targets, mechanisms of action, and blood-brain permeability with DrugBank, PubTator, and PubMed.^72, 103, 104^

### Cosine similarity between drug candidate LINCS perturbation profiles

For each cancer, we calculated the cosine similarity between perturbation profiles in the LINCS level 5 data for the specific cell type we used as the reference for signature reversion.^105^ We clustered these values in a heatmap^101^ by Euclidean distance and the ward.D2 algorithm and then labeled each by the disease-associated gene signature reversion analyses used to identify the drug candidate.

We developed a drug-drug similarity network with the highest 10% of cosine similarity weights for each cancer with the igraph and visnetwork R packages.^106, 107^ In these networks, each node is a candidate where the color indicates the disease-associated gene signature reversion analysis that identified that drug candidate, the edges are weighted based on the cosine similarity, and edge thickness corresponds to the strength of similarity between drug candidates. We applied the Leiden algorithm from igraph to identify drug candidate communities.^106^

### Comparing signature reversion results between disease-associated gene signature reversion analyses

We wanted to compare the signature reversion results between each disease-associated gene signature reversion analysis by cancer. Therefore, we conducted Spearman correlation and linear regression to determine if there is a correlation between the normalized connectivity scores and false discovery rates for the three analyses per cancer. For this analysis, we focused on the signature reversion results for the specific cancer cell line of interest (i.e., GI1 for GBM, HEPG2 for LIHC, A529 for LUAD, and YAPC for PAAD)

### Drug structure similarity analysis

For each cancer, we calculated the Tanimoto Coefficient between drug structures from customCMPdb via the ChemmineR package.^97, 108^ We clustered the Tanimoto Coefficients in a heatmap by Euclidean distance and ward.D2 clustering and labeled drug candidates by which disease-associated gene signature reversion analysis identified the drug candidate. We also constructed drug-drug similarity networks based on drug structure similarity (Tanimoto Coefficient).^106, 107^ In these networks, each node is a drug candidate colored by the disease-associated gene signature reversion analysis that identified it, and the edges are weighted based on the Tanimoto Coefficient where their thickness correlates with the Tanimoto Coefficient. Therefore, the thicker the edge the stronger the similarity between candidates.

### Network centrality analysis

For this analysis, we considered a gene high weight if it was one of the 10 most highly weighted genes contributing to a latent variable. We determined this threshold by sampling the top genes and evaluating the gene overlap between different latent variables. We determined that there was a progressively larger overlap between top genes of different latent variables as the gene set was increased (**Supplemental Figure 7A**). For each gene in the protein-protein interaction network from the STRING database, we calculated the degree, betweenness, and Eigenvalue of centrality^29, 106^ and performed two-tailed Wilcox tests for each network metric followed by a Bonferroni p-value adjustment. We log2-transformed the centrality metrics for plotting and visualization. We compared the distribution of the log2-transformed metrics of the unique genes included in the disease-associated gene signature by each of the three approaches (i.e., DESeq2, limma, transfer learning).

We did not consider overlapping genes to maintain the assumption of independence for both the Kruskal-Wallis and pairwise Wilcox tests. Then we conducted a Kruskal-Wallis test to determine if there were differences between the three sets within a cancer. If the Kruskal-Wallis test was significant (ɑ=0.05), then we applied a pairwise Wilcox test with Benjamini Hochberg p-value adjustment.

### GBM in vitro cell viability assay

We conducted one initial cell viability drug screen with 3 replicates of pamidronate, nimodipine, saxagliptin, and icosapent in the U251 and NHA cell lines using CellTiter-Glo (**Supplemental Figure 36-37**). Then, we did a secondary cell viability durg screen with 3 replicates of pamidronate, nimodipine, and temozolomide in U251, JX39 (**Figure 6 A-D),** and NHA cell systems (**Supplemental Figure 39**). We performed a third cell viability drug screen with 6 replicates of pamidronate, nimodipine, and temozolomide in both U251 and JX39 cell culture systems (**Supplemental Figure 38**).

For the cell viability assays, pamidronate was resuspended in water to 100mM which was then serially diluted in water to produce 1000X stocks for use in drug treatments. Water was used as the vehicle control for all experiments with pamidronate. Nimodipine, saxagliptin, and icosapent were resuspended in DMSO to 50mM which was then serially diluted in DMSO to produce 1000X stocks for use in drug treatments. DMSO was used as the vehicle control for all experiments with nimodipine, saxagliptin, and icosapent.

To conduct the cell viability assay, 1000 U251 GBM cells or 2000 JX39 GBM cells isolated from patient-derived xenograft were plated per 96 well in DMEM/F12 in brain tumor-initiating cell maintenance conditions, treated with the indicated concentrations of drug or vehicle control the day after plating, and cell viability measured with CellTiter-Glo after five or seven days respectively, similar to our published reports.^109, 110^ 2000 Immortalized human astrocytes (NHAs; astrocytes immortalized with hTERT and E7) were plated in astrocyte media, treated with the indicated concentrations of drug or vehicle control the day after plating, and cell viability was measured with CellTiter-Glo after five similar to prior publications.^110, 111^

## Supporting information

FILE1_sample_counts

FILE2_clinical_trial_cancer_summary

FILE3_top_candidate_multi_sheet_pathways

FILE4_computer_system_and_packages_info_dockers_conda

## Acknowledgments

We would like to thank all the members of the Lasseigne Lab for their feedback on this manuscript. JLF, VHO, ABH, and BNL are supported by R03OD030604 (to BNL); JLF and BNL are supported by an ACS-IRG (to BNL); Victoria Flanary is supported by 5T32GM008361-31; JLF, EJW, VHO, and TCH are supported by R00HG009678 (to BNL); JLF, TCH, VHO, and BNL are supported by UAB funds to the Lasseigne Lab.

## Authors Contributions

Jennifer L. Fisher: Conceptualization, Methodology, Software, Formal Analysis, Investigation, Data Curation, Writing - Original Draft, Writing - Review & Editing, Visualization. Elizabeth J. Wilk: Methodology, Validation, Writing - Review & Editing. Vishal H. Oza: Conceptualization, Methodology, Writing - Review & Editing. Timothy C. Howton: Writing - Review & Editing, Project administration. Victoria Flanary: Validation, Writing - Review & Editing. Amanda D. Clark: Validation, Writing - Review & Editing. Anita B. Hjelmeland: Investigation, Formal Analysis, Resources, Writing - Review & Editing. Brittany N. Lasseigne: Conceptualization, Methodology, Resources, Writing - Review & Editing, Supervision, Project administration, Funding acquisition.

## Supplemental Files

**Supplemental File 1:** Table of the number of samples for the project. File name: FILE1_sample_counts.xlsx The numbers of tumor and control non-tumor samples for each cancer and control tissue are listed in

**Supplemental File 1** by database (TCGA/GTEX).

**Supplemental File 2:** Table of the permutation testing of random drug selection results for all methods across cancers. File name: FILE2_220608_clinical_trial_cancer_summary.csv We performed permutation testing of a matched number of random sets of drugs for each as a comparison for the clinical trials (**Supplemental File 2)**. For each cancer and disease-associated gene signature method, we note the number of drug candidates from signature reversion that have been in previous clinical trials (CT), the number that were not in previous clinical trials (Not_CT), and the permutation one-tailed Wilcox test results of significance (significant). We also included the PRISM permutation one-tailed Wilcox test results of significance in the PRISM column (PRISM).

**Supplemental File 3:** Pathway results for the top candidates for GBM. File name: FILE3_top_candidate_multi_sheet_pathways.xlsx

**Supplemental File 3** contains gprofiler2 functional enrichment of the perturbation profiles (level 5 LINCS data) of pamidronate, nimodipine, saxagliptin, and icosapent. The set column was added to the gprofiler2 output to note if enriched gene sets were from the up or down perturbed genes. The columns query, significant, p_value, term_size, query_size, intersection_size, precision, recall, term_id, source, term_name, effective_domain_size, and source_order are from the gprofiler2 output. Below is the column information from the gprofiler vignette:^112^

- query - the name of the input query which by default is the order of the query with the prefix “query_.” This can be changed by using a named list input.
- significant - indicator for statistically significant results
- p_value - hypergeometric p-value after correction for multiple testing
- term_size - number of genes that are annotated to the term
- query_size - number of genes that were included in the query. This might be different from the size of the original list if:
- any genes were mapped to multiple Ensembl gene IDs
- any genes failed to be mapped to Ensembl gene IDs
- the parameter ordered_query = TRUE and the optimal cutoff for the term was found before the end of the query
- the domain_scope was set to “annotated” or “custom”
- intersection_size - the number of genes in the input query that are annotated to the corresponding term
- precision - the proportion of genes in the input list that are annotated to the function (defined as intersection_size/query_size)
- recall - the proportion of functionally annotated genes that the query recovers (defined as intersection_size/term_size)
- term_id - unique term identifier (e.g GO:0005005)
- source - the abbreviation of the data source for the term (e.g. GO:BP)
- term_name - the short name of the function
- effective_domain_size - the total number of genes “in the universe” used for the hypergeometric test
- source_order - numeric order for the term within its data source (this is important for drawing the results)

**Supplemental File 4:** Computing system and package version information. File name: FILE4_computer_system_and_packages_info_dockers_conda.pdf Detailed computer and package version information are included in **Supplemental File 4**.

## Supplemental Figures

**Supplemental Figure 1:**
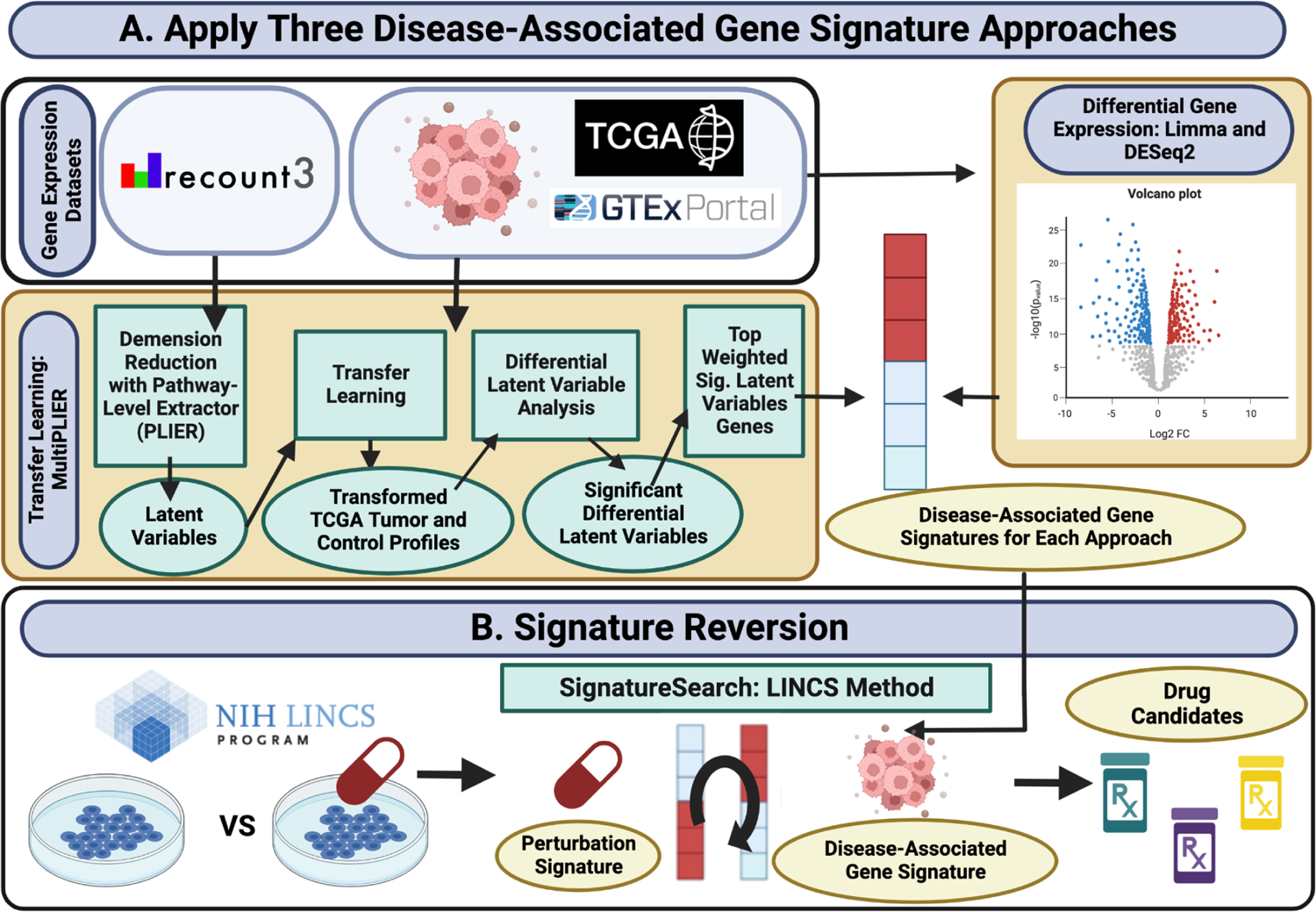
Detailed overview of the signature reversion with three disease-associated gene signatures. **A)** The process for making the disease-associated gene signatures for the three approaches. **B)** The signature reversion methodology.

**Supplemental Figure 2:**
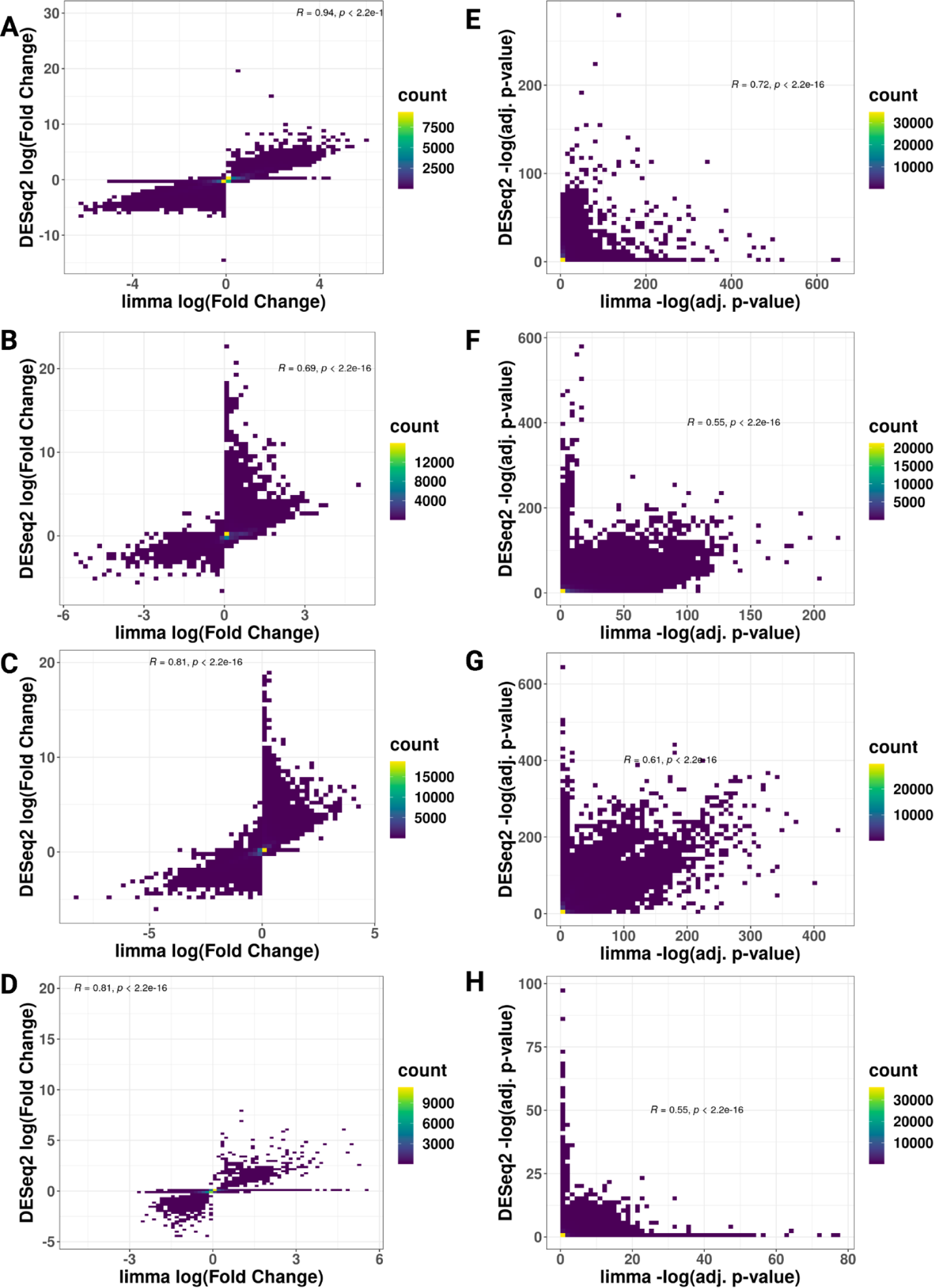
Scatter plots for limma and DESeq2 log fold change and adjusted p-value comparisons. **A-D)** Scatter plot of the limma log fold change and DESeq2 log fold change for GBM, LIHC, LUAD, and PAAD, respectively. **E-H)** Scatter plot of the limma adjusted p-value and DESeq2 adjusted p-value for GBM, LIHC, LUAD, and PAAD, respectively. Spearman correlation and p-value from linear regression models are also plotted on each panel.

**Supplemental Figure 3:**
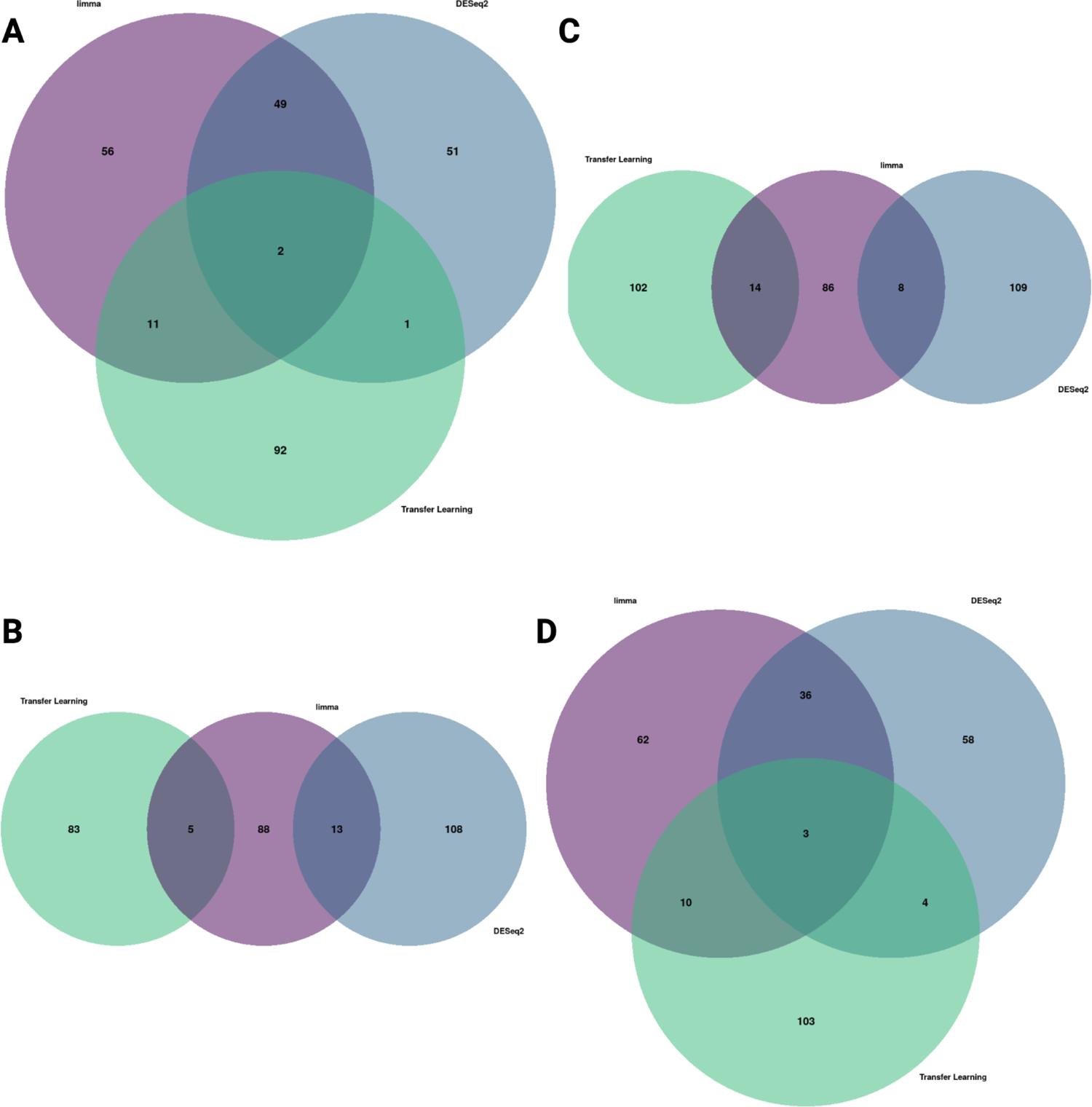
Overlap of disease-associated gene signatures between Methods. **A-D)** Venn diagrams of the overlap between disease-associated gene signatures for GBM, LIHC, LUAD, and PAAD, respectively.

**Supplemental Figure 4:**
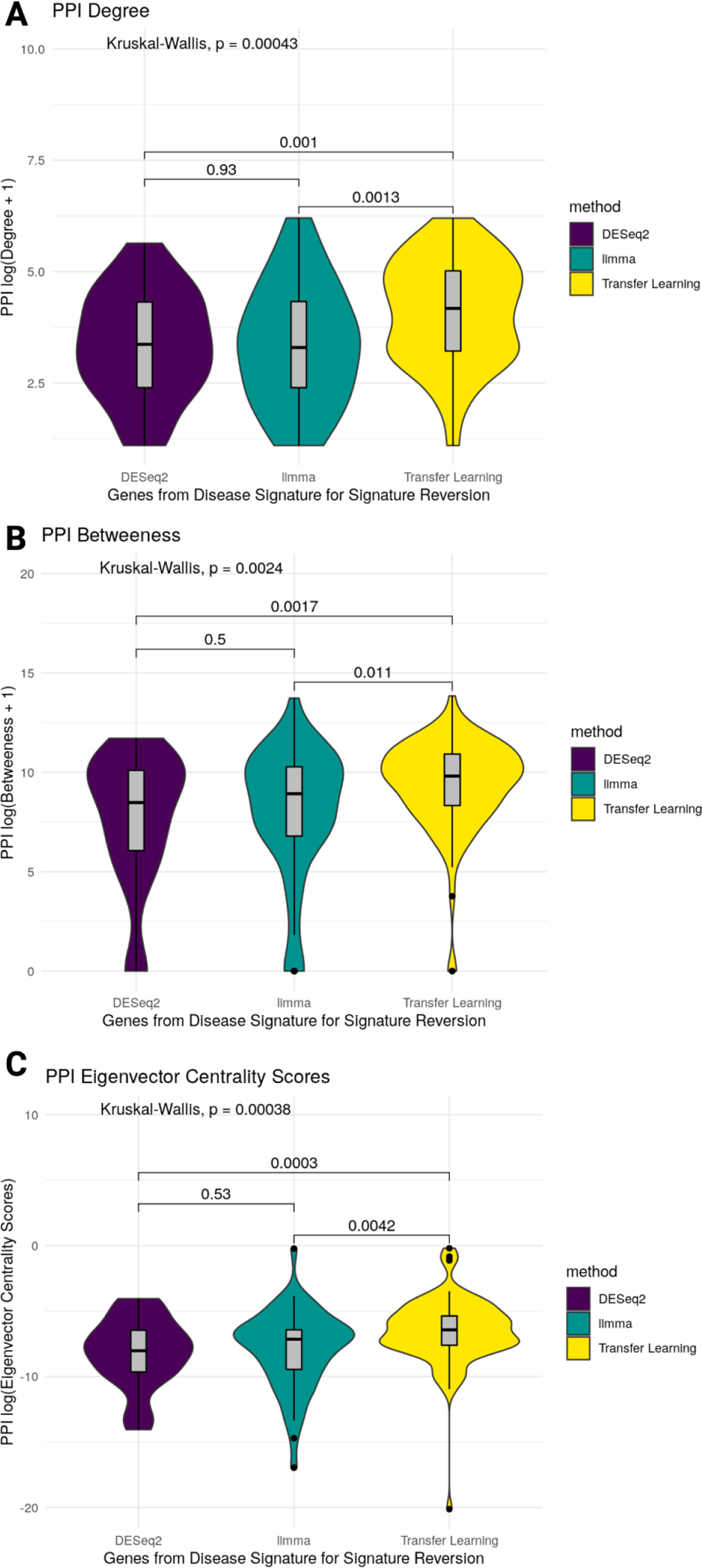
Protein-protein interaction network centrality metrics for GBM disease-associated signature genes. Violin plots for **A)** degree, **B)** betweenness, and **C)** eigenvector centrality. All tests for this figure are Kruskal-Wallis and Wilcox tests with a Bonferroni multiple hypothesis adjustment.

**Supplemental Figure 5:**
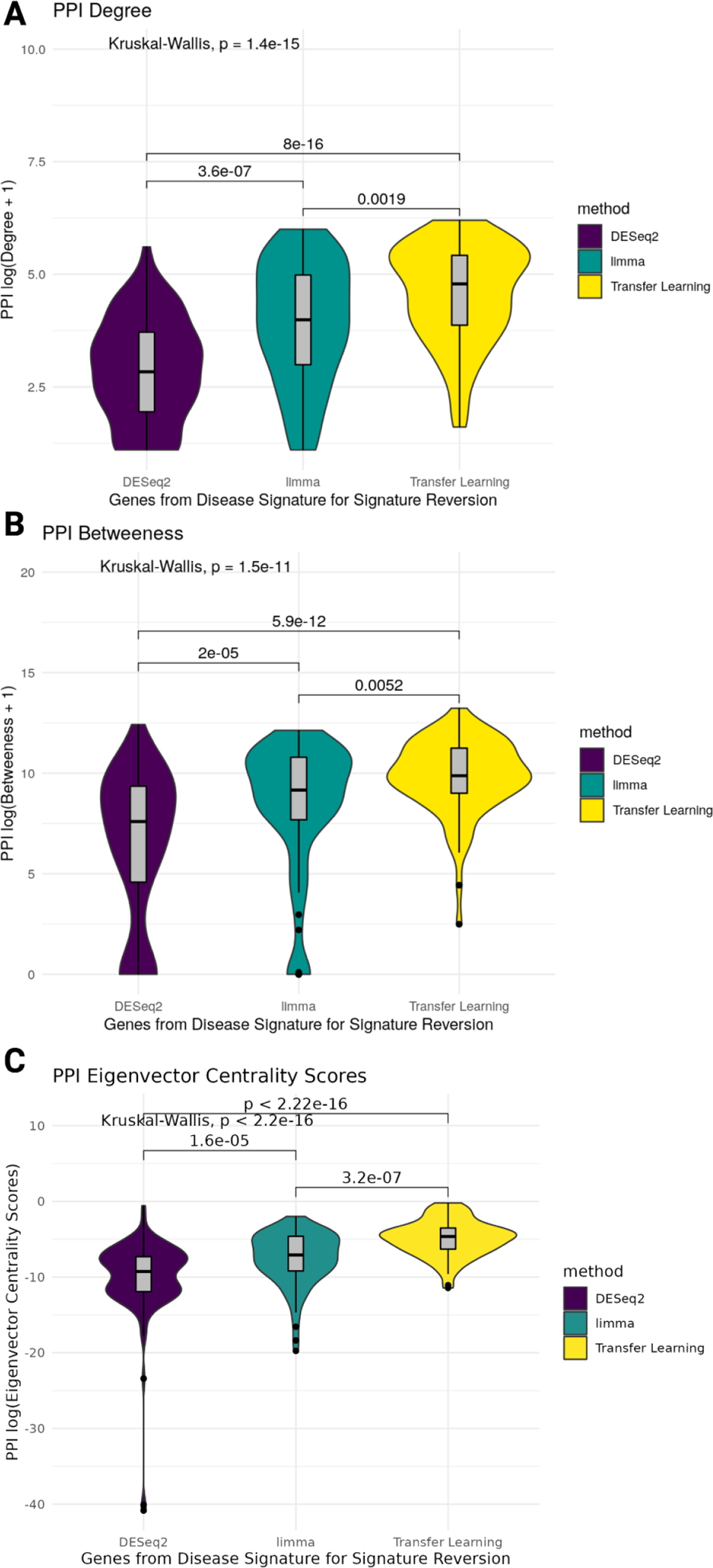
Protein-protein interaction network centrality metrics for LIHC disease-associated signature genes. Violin plots for **A)** degree, **B)** betweenness, and **C)** eigenvector centrality. All tests for this figure are Kruskal-Wallis and Wilcox tests with a Bonferroni multiple hypothesis adjustment.

**Supplemental Figure 6:**
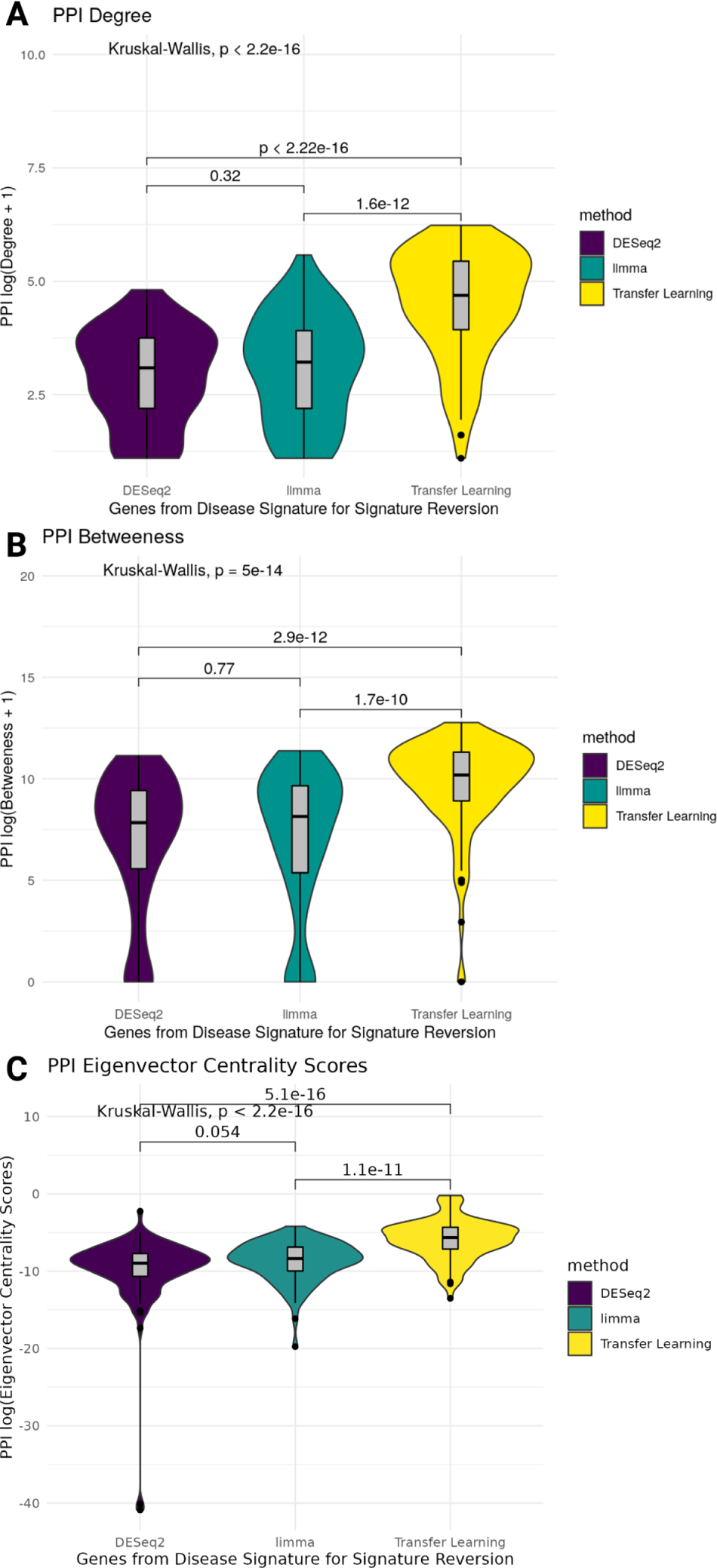
Protein-protein interaction network centrality metrics for LUAD disease-associated signature genes. Violin plots for **A)** degree, **B)** betweenness, and **C)** eigenvector centrality. All tests for this figure are Kruskal-Wallis and Wilcox tests with a Bonferroni multiple hypothesis adjustment.

**Supplemental Figure 7:**
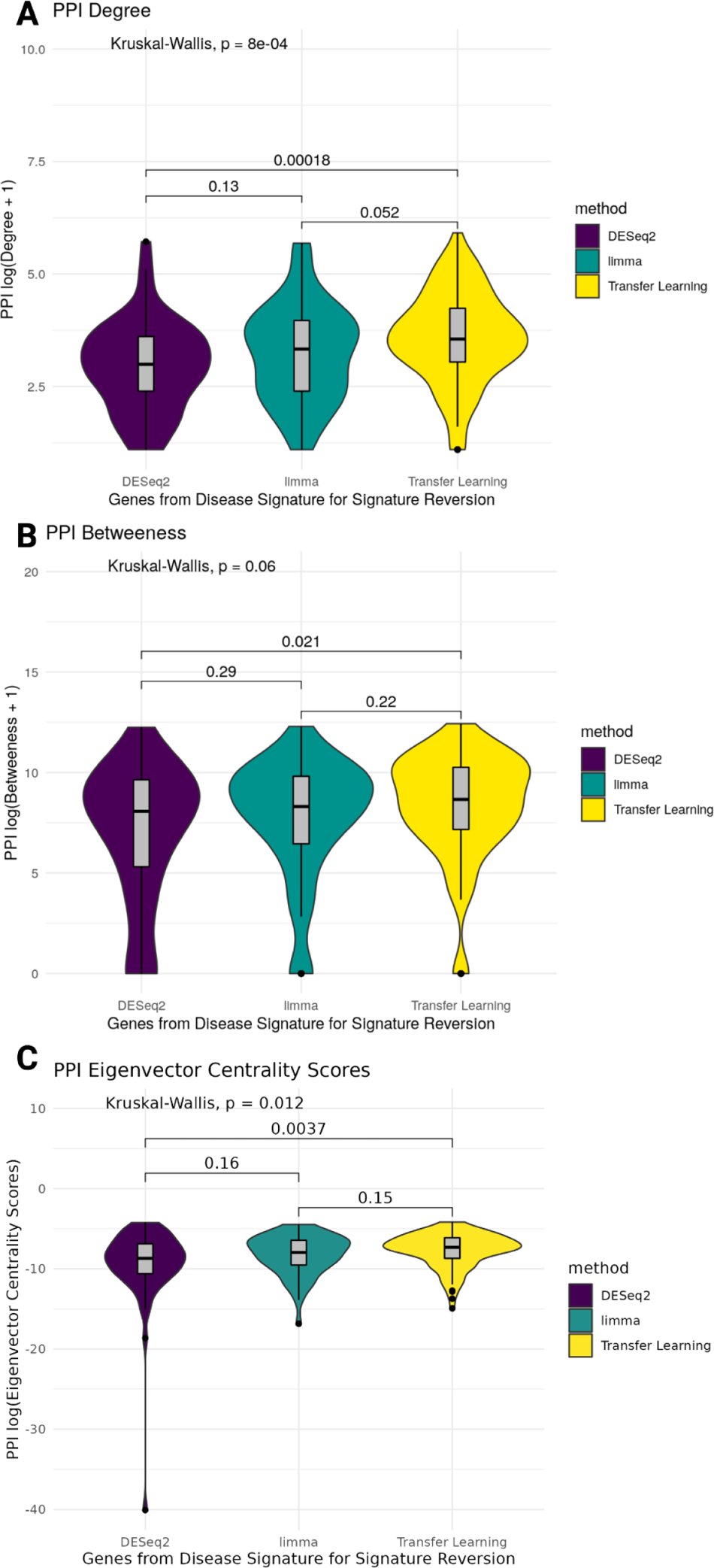
Protein-protein interaction network centrality metrics for PAAD disease-associated signature genes. Violin plots for **A)** degree, **B)** betweenness, and **C)** eigenvector centrality. All tests for this figure are Kruskal-Wallis and Wilcox tests with a Bonferroni multiple hypothesis adjustment.

**Supplemental Figure 8:**
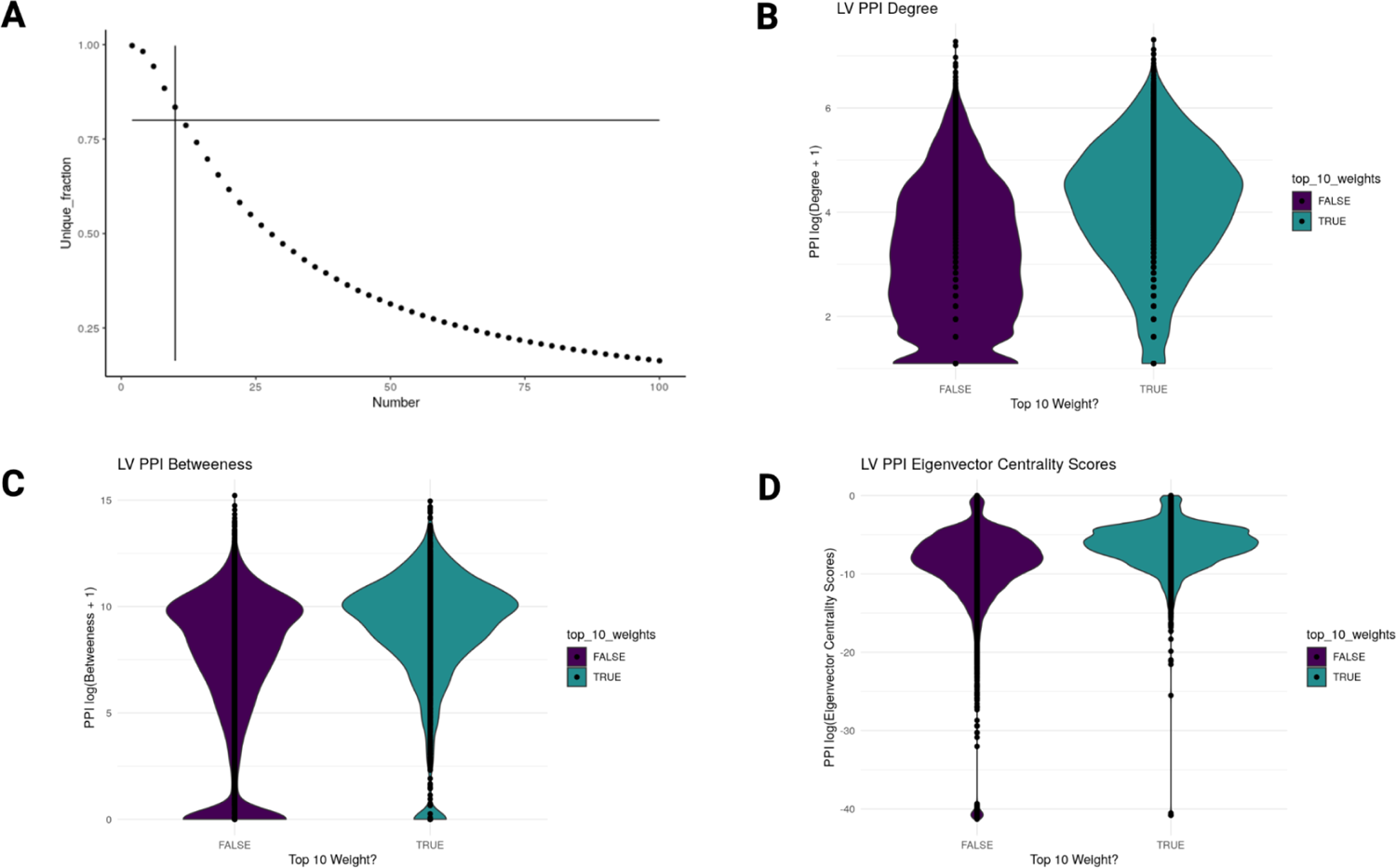
Protein-protein interaction network centrality metrics for top latent variable genes. **A)** Ratio of the number of genes in the top most weighted genes of all latent variables to all of the genes included in the top most weighted genes of all latent variables (y-axis) plotted by a given number of top most weighted genes (x-axis). The vertical line is at 10 (i.e., 10 genes each for the 385 latent variables) and the horizontal line at 0.80 was the cut-off for inclusion in the network centrality analysis. **B)** PPI degree violin plot (adj. P-value = 0). **C)** PPI betweenness (adj. P-value = 1.883475e-176 **D)** PPI eigenvector centrality (adj. P-value = 1.231573e-231). All statistical tests for this figure are Wilcox tests with a Bonferroni multiple hypothesis adjustment.

**Supplemental Figure 9:**
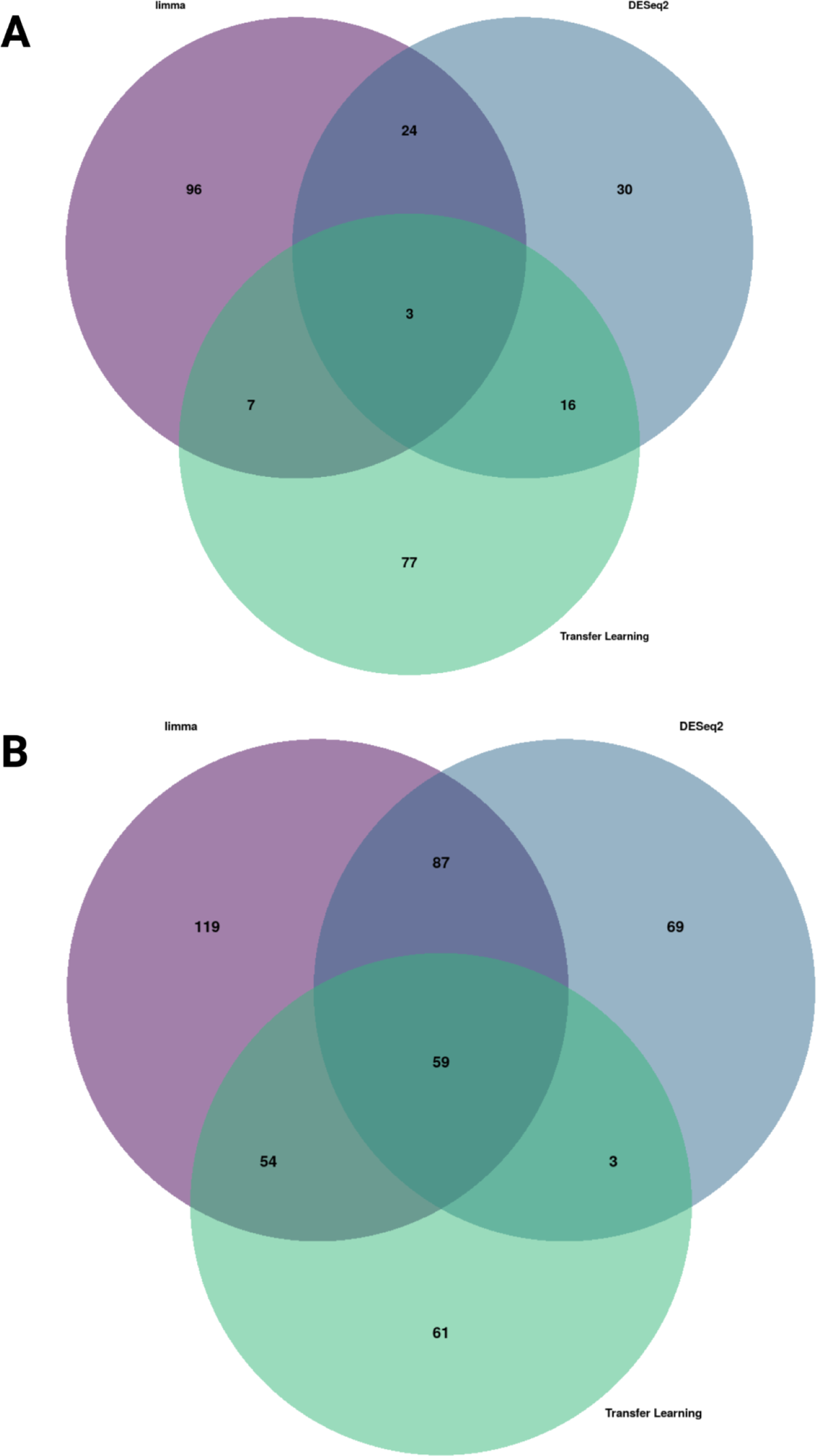
Enriched gene set overlap Venn diagrams for GBM. **A)** Up and **B)** down-regulated gene set overlap from the functional enrichment results for each disease-associated gene signature.

**Supplemental Figure 10:**
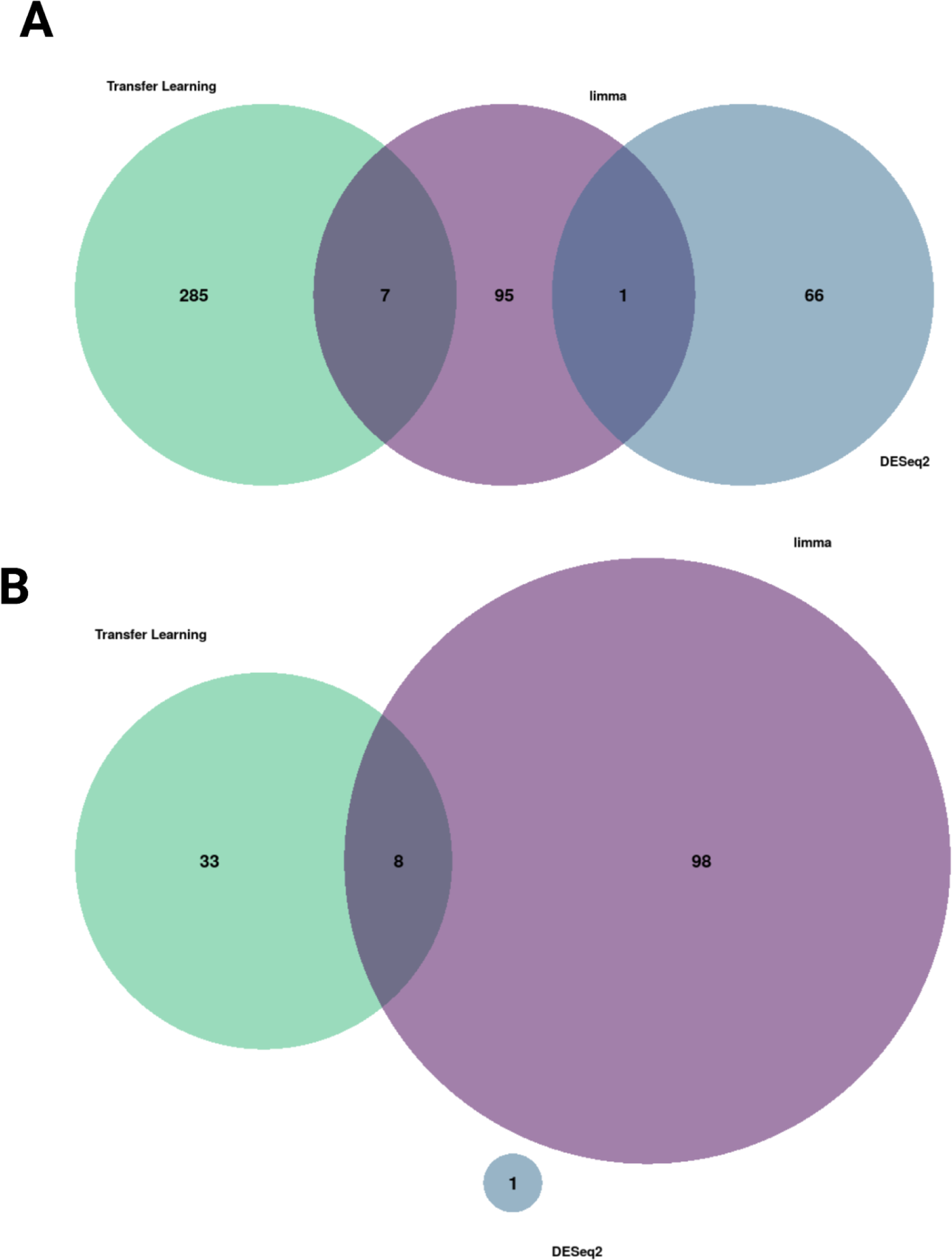
Enriched gene set overlap Venn diagrams for LIHC. A) **Up and B)** down-regulated gene set overlap from the functional enrichment results for each disease-associated gene signature.

**Supplemental Figure 11:**
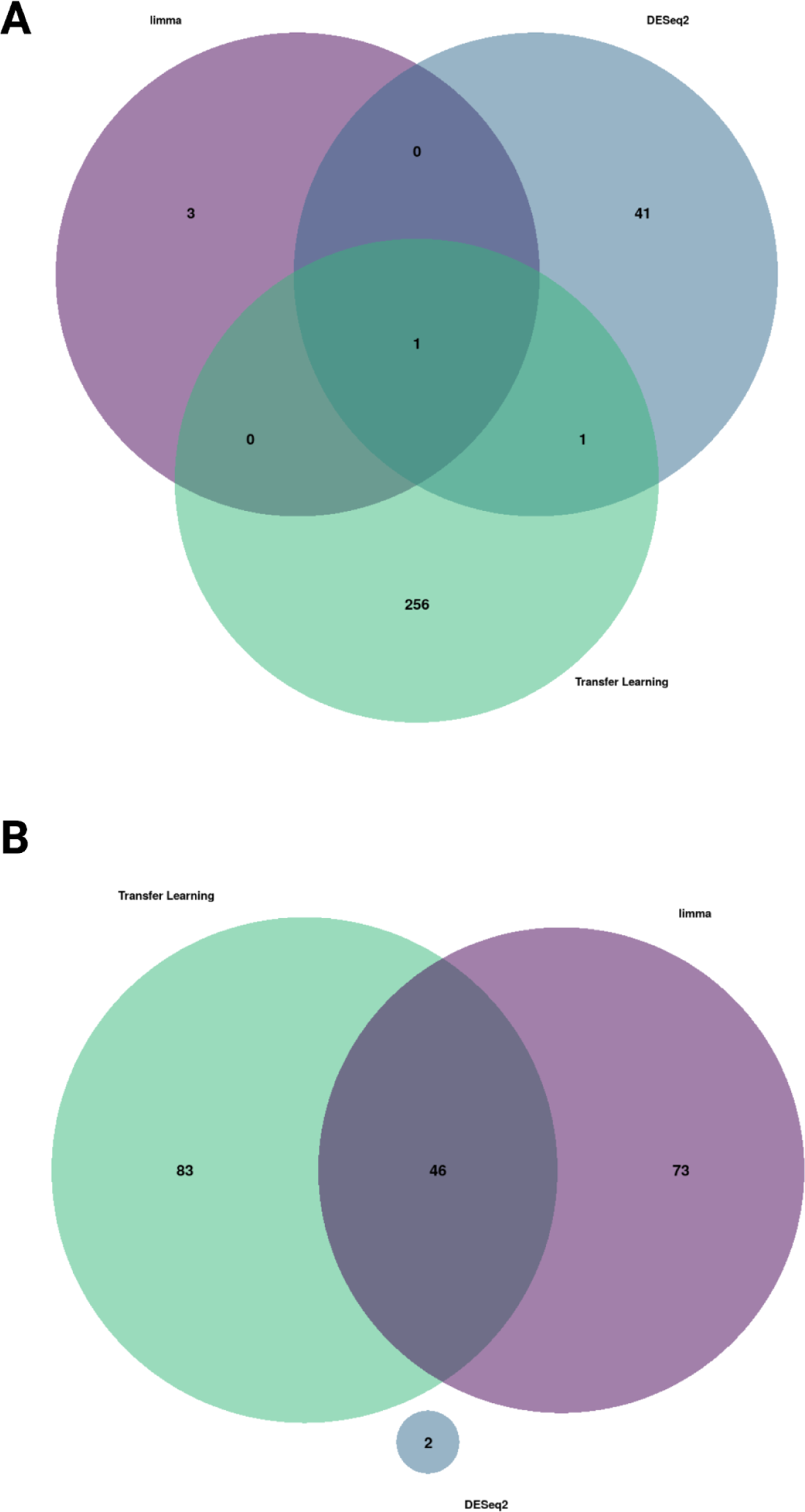
Enriched gene set overlap Venn diagrams for LUAD. A) **Up and B)** down-regulated gene set overlap from the functional enrichment results for each disease-associated gene signature.

**Supplemental Figure 12:**
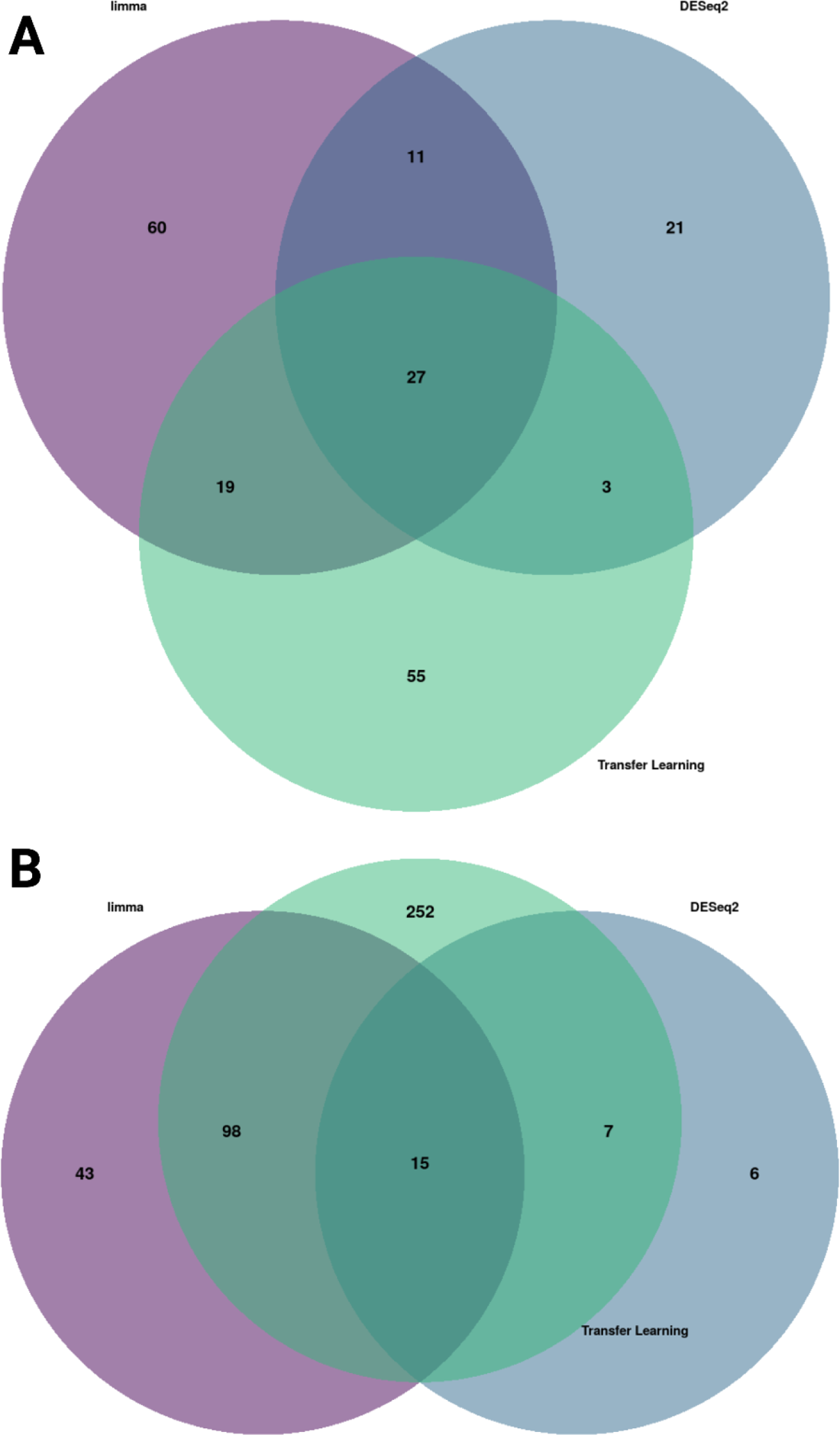
Enriched gene set overlap Venn diagrams for PAAD. A) **Up and B)** down-regulated gene set overlap from the functional enrichment results for each disease-associated gene signature.

**Supplemental Figure 13:**
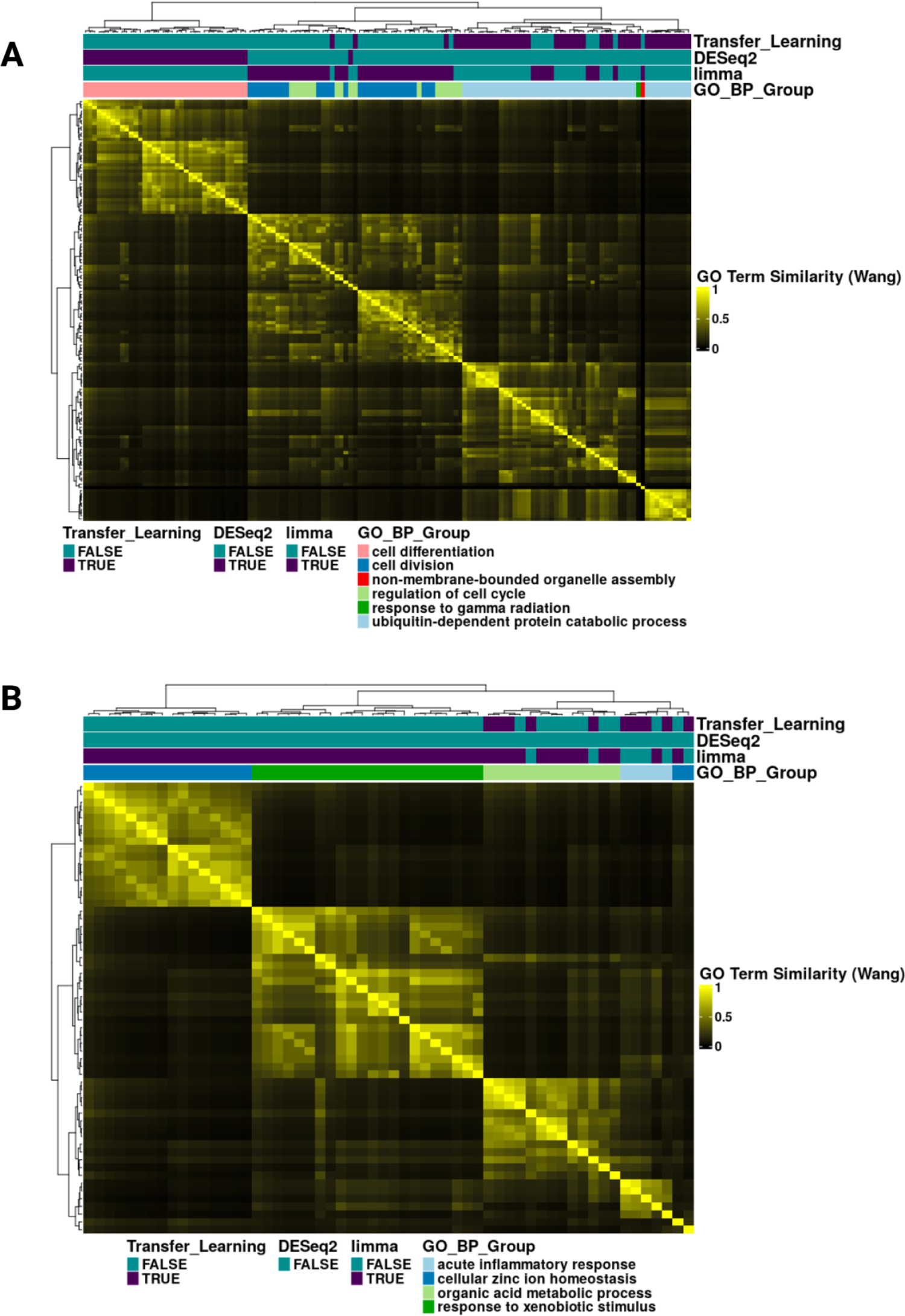
Heatmaps for LIHC GO_BP terms. **A)** Heatmap of the Gene Ontology (GO) term semantic similarity (Wang method) of the up-regulated enriched GO Biological Process terms from the LIHC disease-associated gene signatures. Each term is associated with a disease-associated gene signature if the row for that method is purple. In addition, the Gene Ontology Biological Process terms were grouped together based on common parent terms, and the different parent term groups are indicated in the GO_BP_Group. **B)** A heatmap of the GO term semantic similarity (Wang method) of down-regulated enriched Gene Ontology Biological Process terms for LIHC.

**Supplemental Figure 14:**
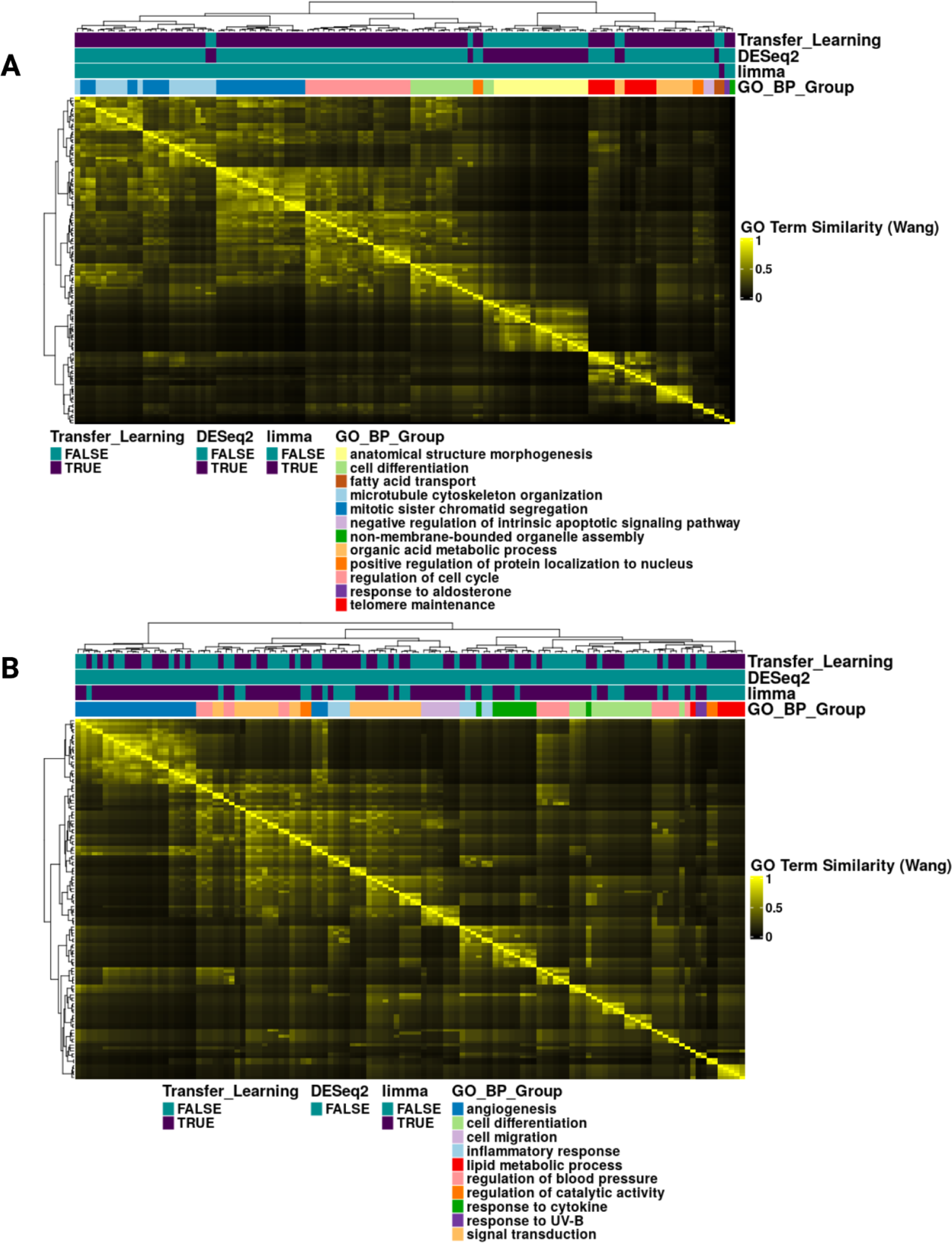
GO heatmaps for LUAD. **A)** Heatmap of the Gene Ontology (GO) term semantic similarity (Wang method) of the up-regulated enriched GO Biological Process terms from the LUAD disease-associated gene signatures. Each term is associated with a disease-associated gene signature if the row for that method is purple. In addition, the Gene Ontology Biological Process terms were grouped together based on common parent terms, and the different groups are indicated in the GO_BP_Group. **B)** A heatmap of the GO term semantic similarity (Wang method) of down-regulated enriched Gene Ontology Biological Process terms for LUAD.

**Supplemental Figure 15:**
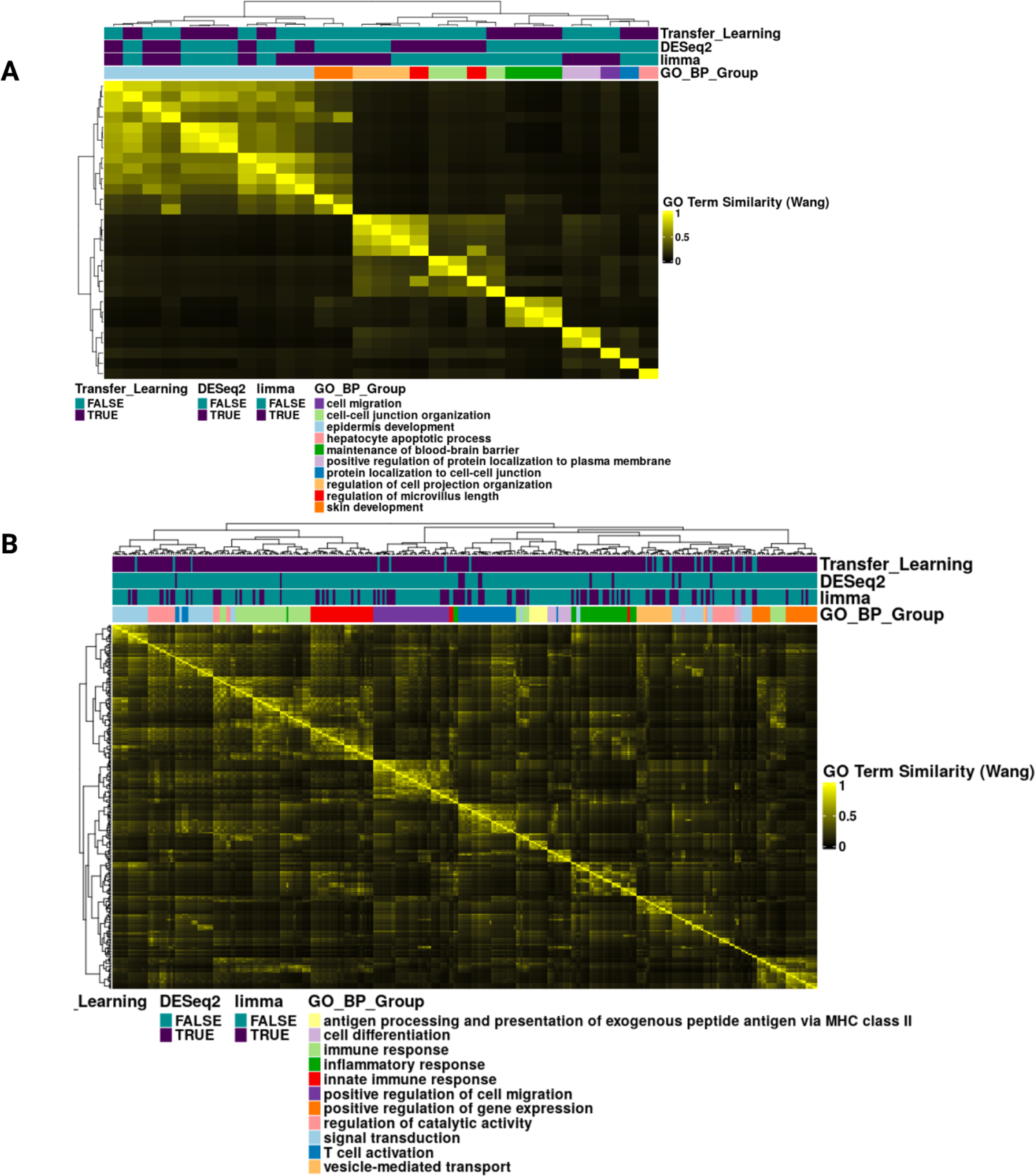
GO heatmaps for PAAD. **A)** Heatmap of the Gene Ontology (GO) term semantic similarity (Wang method) of the up-regulated enriched GO Biological Process terms from the PAAD disease-associated gene signatures. Each term is associated with a disease-associated gene signature if the row for that method is purple. In addition, the Gene Ontology Biological Process terms were grouped together based on common parent terms, and the different groups are indicated in the GO_BP_Group. **B)** A heatmap of the GO term semantic similarity (Wang method) of down-regulated enriched Gene Ontology Biological Process terms for PAAD.

**Supplemental Figure 16:**
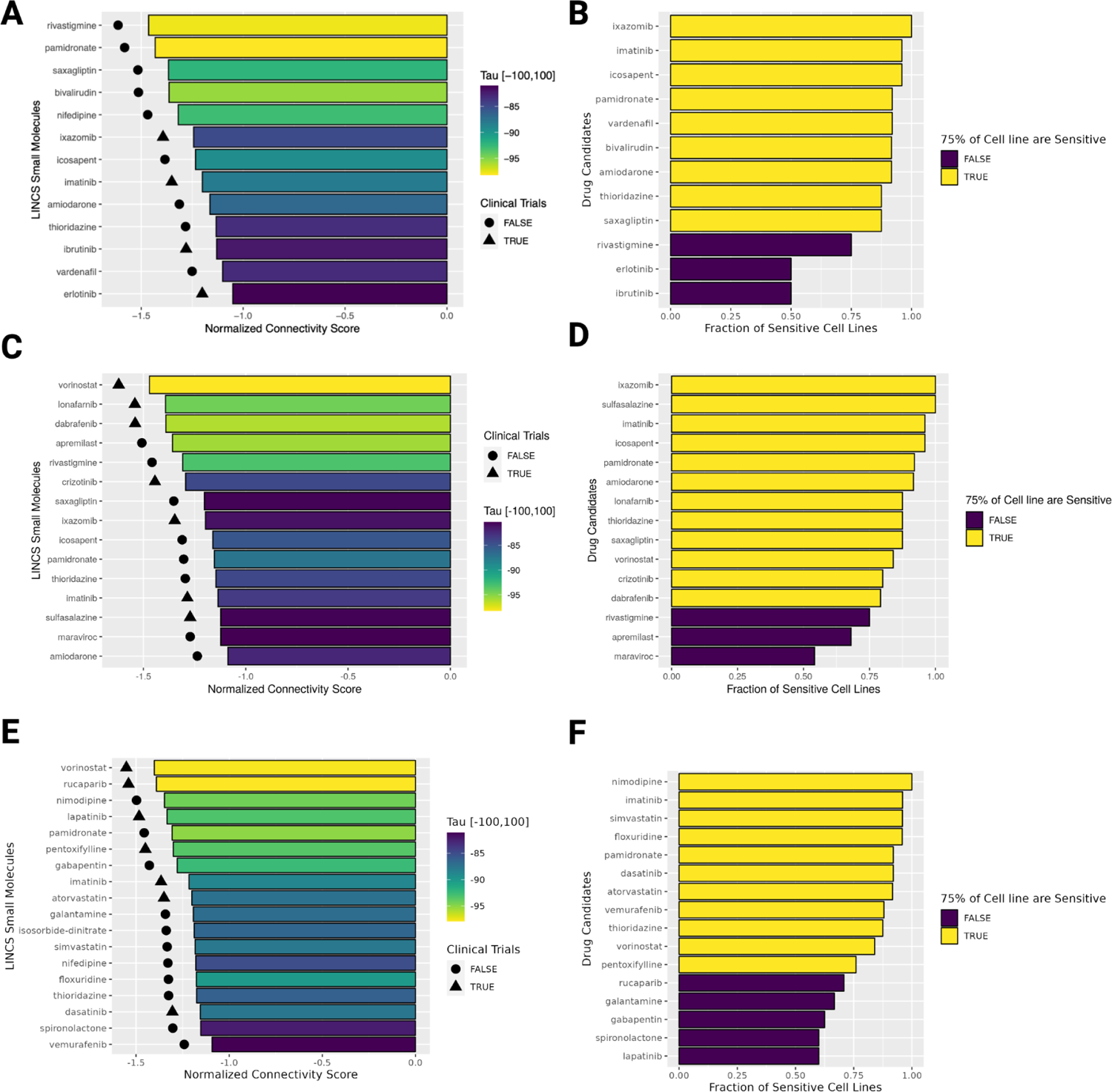
DESeq2, limma, and transfer learning signature reversion results for GBM. **A)** Barplot of the top drug repurposing candidates ordered by Normalized Connectivity Scores and with a −80 Tau cutoff for the DESeq2 disease-associated signature. **B)** Barplot of the fraction of sensitive cell lines in the PRISM dataset for DESeq2 disease-associated signature drug candidates. **C)** Barplot of the top drug repurposing candidates identified from the limma disease-associated signature. **D)** Barplot of the fraction of sensitive cell lines in the PRISM dataset for limma disease-associated signature drug candidates. **E)** Barplot of the top drug repurposing candidates identified from the transfer learning disease-associated signature. **F)** Barplot of the fraction of sensitive cell lines in the PRISM dataset for transfer learning disease-associated signature drug candidates.

**Supplemental Figure 17:**
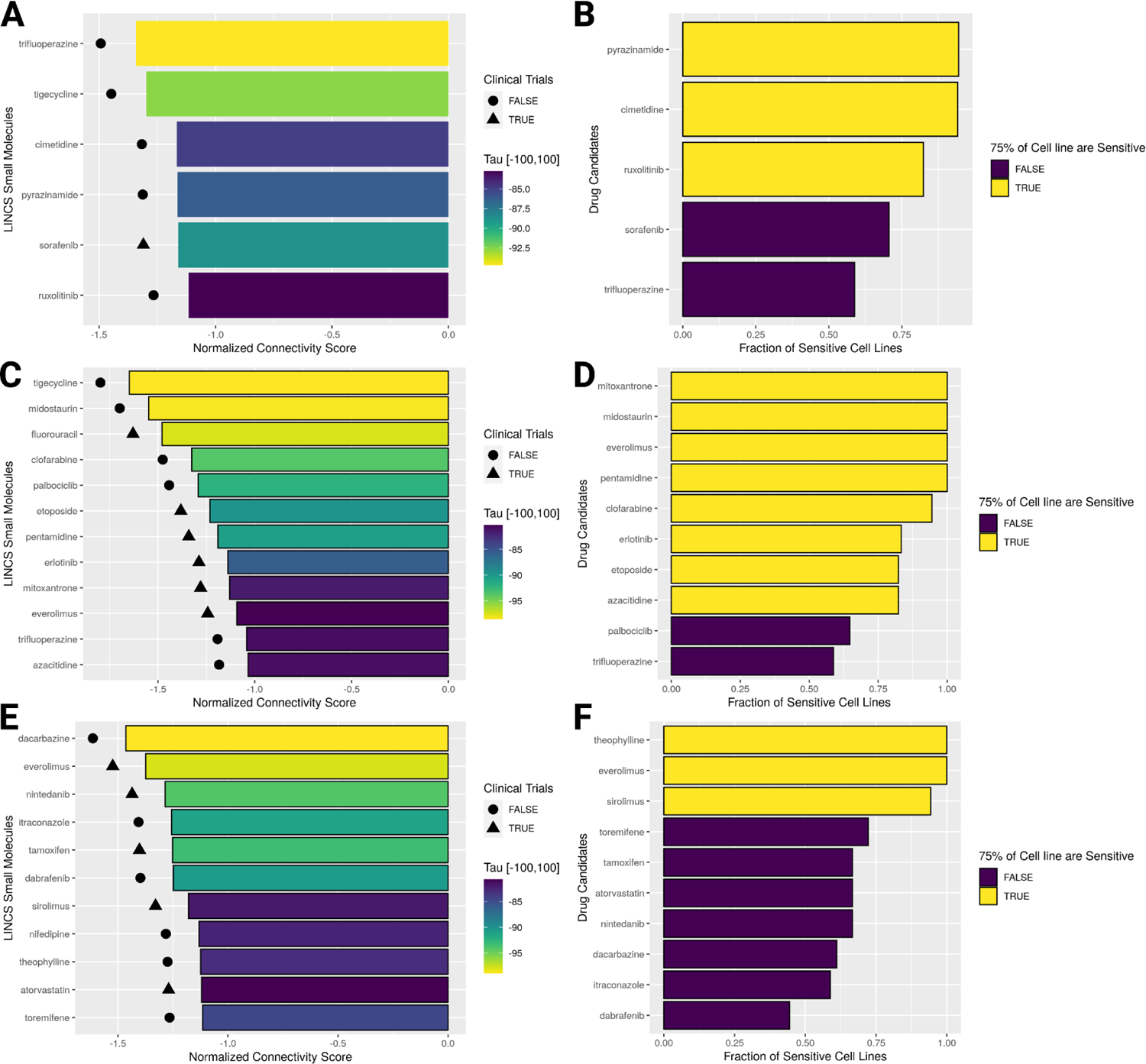
DESeq2, limma, and transfer learning signature reversion results for LIHC. **A)** Barplot of the top drug repurposing candidates ordered by Normalized Connectivity Scores and with a −80 Tau cutoff for the DESeq2 disease-associated signature. **B)** Barplot of the fraction of sensitive cell lines in the PRISM dataset for DESeq2 disease-associated signature drug candidates. **C)** Barplot of the top drug repurposing candidates identified from the limma disease-associated signature. **D)** Barplot of the fraction of sensitive cell lines in the PRISM dataset for limma disease-associated signature drug candidates. **E)** Barplot of the top drug repurposing candidates identified from the transfer learning disease-associated signature. **F)** Barplot of the fraction of sensitive cell lines in the PRISM dataset for transfer learning disease-associated signature drug candidates.

**Supplemental Figure 18:**
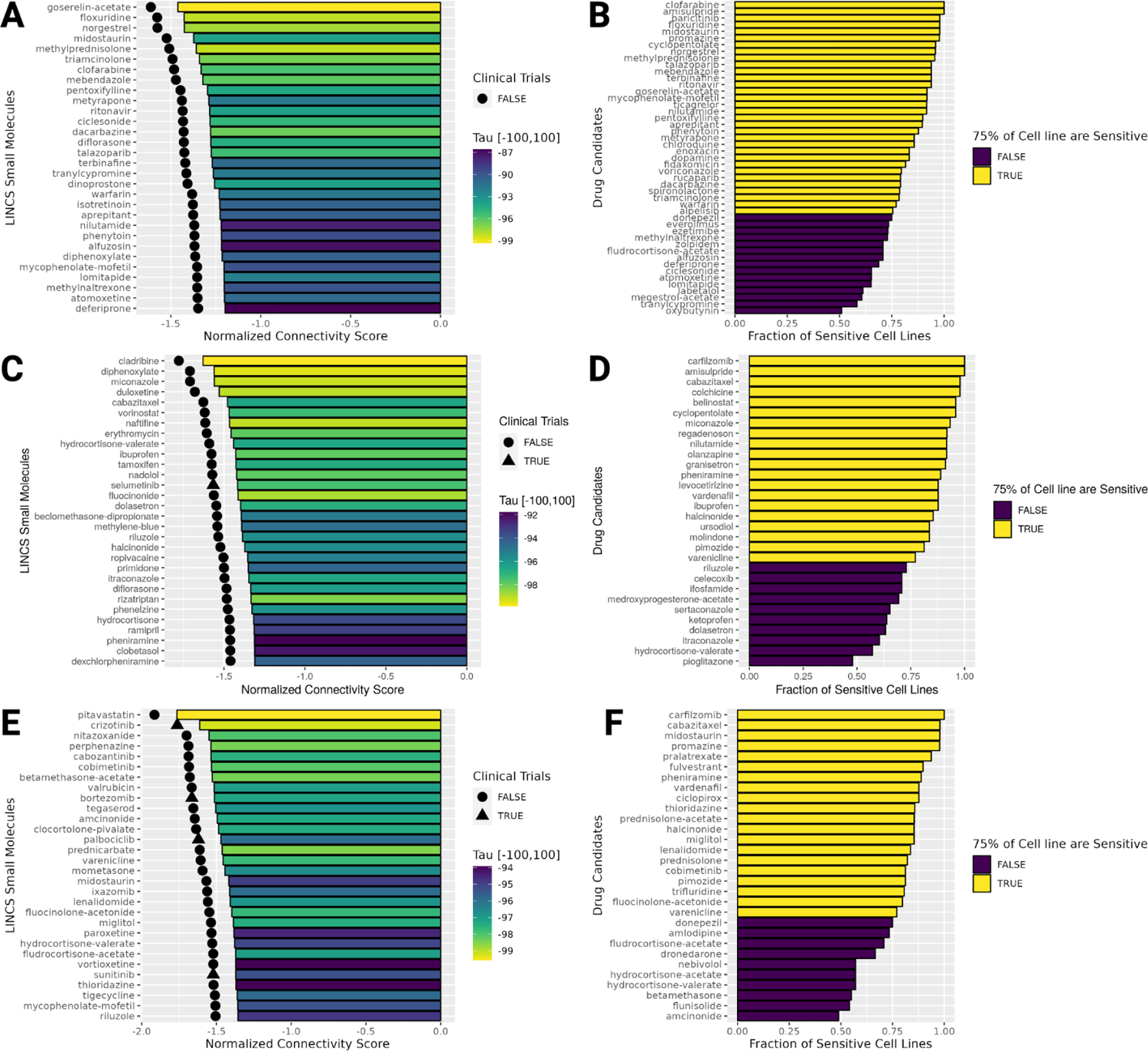
DESeq2, limma, and transfer learning signature reversion results for LUAD. **A)** Barplot of the top drug repurposing candidates ordered by Normalized Connectivity Scores and with a −80 Tau cutoff for the DESeq2 disease-associated signature. **B)** Barplot of the fraction of sensitive cell lines in the PRISM dataset for DESeq2 disease-associated signature drug candidates. **C)** Barplot of the top drug repurposing candidates identified from the limma disease-associated signature. **D)** Barplot of the fraction of sensitive cell lines in the PRISM dataset for limma disease-associated signature drug candidates. **E)** Barplot of the top drug repurposing candidates identified from the transfer learning disease-associated signature. **F)** Barplot of the fraction of sensitive cell lines in the PRISM dataset for transfer learning disease-associated signature drug candidates.

**Supplemental Figure 19:**
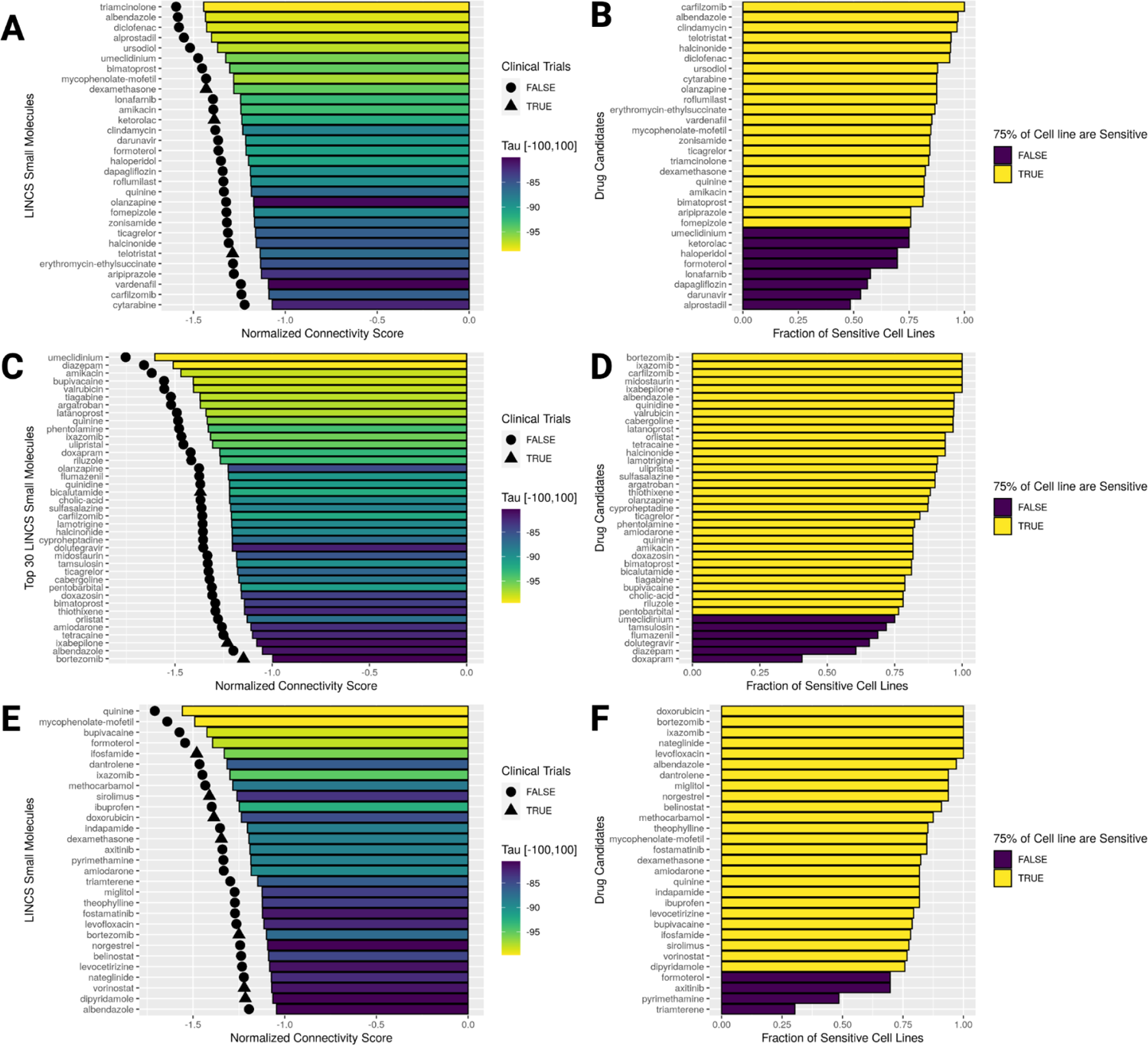
DESeq2, limma, and transfer learning signature reversion results for PAAD. **A)** Barplot of the top drug repurposing candidates ordered by Normalized Connectivity Scores and with a −80 Tau cutoff for the DESeq2 disease-associated signature. **B)** Barplot of the fraction of sensitive cell lines in the PRISM dataset for DESeq2 disease-associated signature drug candidates. **C)** Barplot of the top drug repurposing candidates identified from the limma disease-associated signature. **D)** Barplot of the fraction of sensitive cell lines in the PRISM dataset for limma disease-associated signature drug candidates. **E)** Barplot of the top drug repurposing candidates identified from the transfer learning disease-associated signature. **F)** Barplot of the fraction of sensitive cell lines in the PRISM dataset for transfer learning disease-associated signature drug candidates.

**Supplemental Figure 20:**
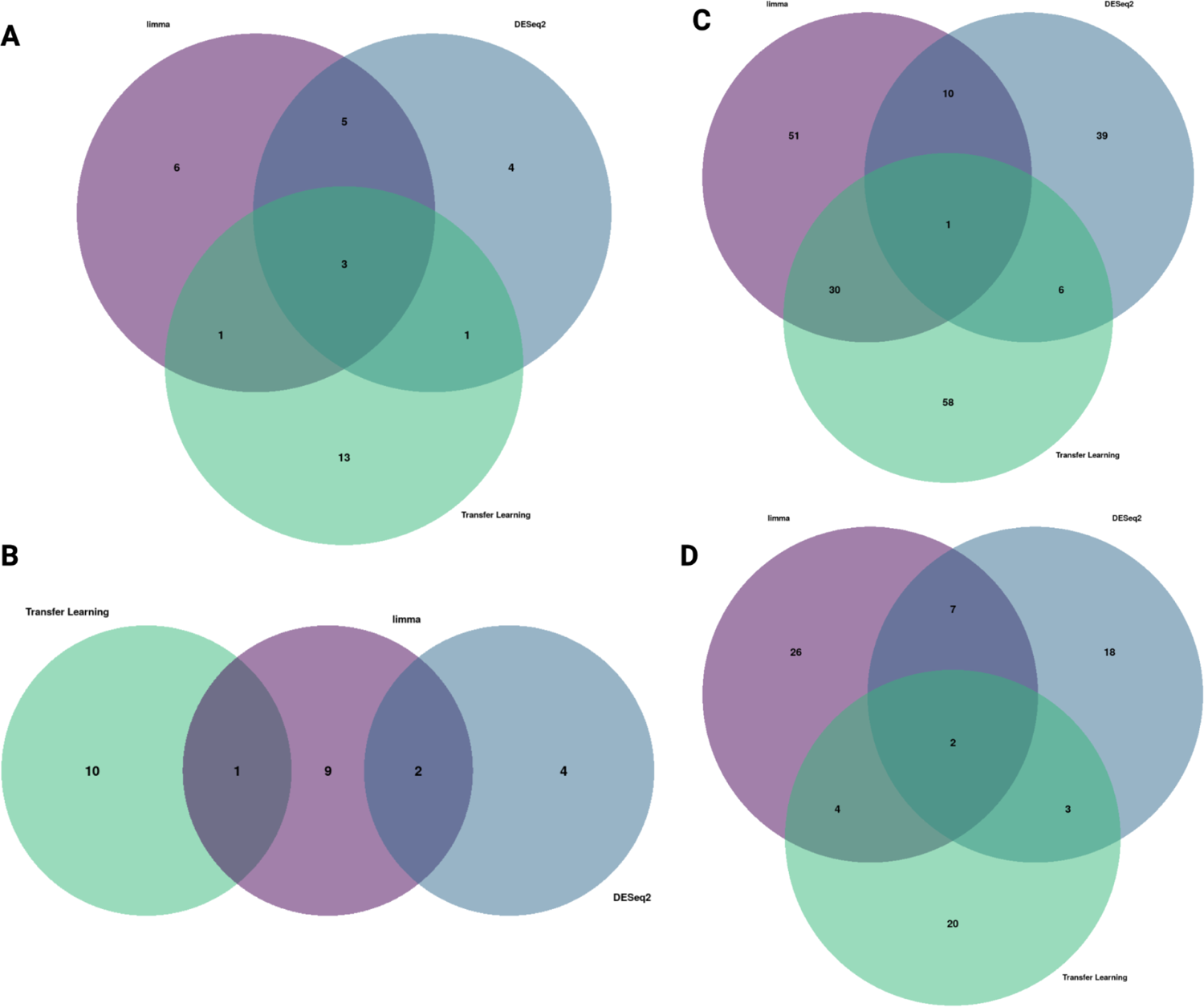
**A-D)** Venn diagrams of the overlapping drug candidates between disease-associated signatures from each method with a negative normalized connectivity score and a tau value less than −80 for GBM, LIHC, LUAD, and PAAD, respectively.

**Supplemental Figure 21:**
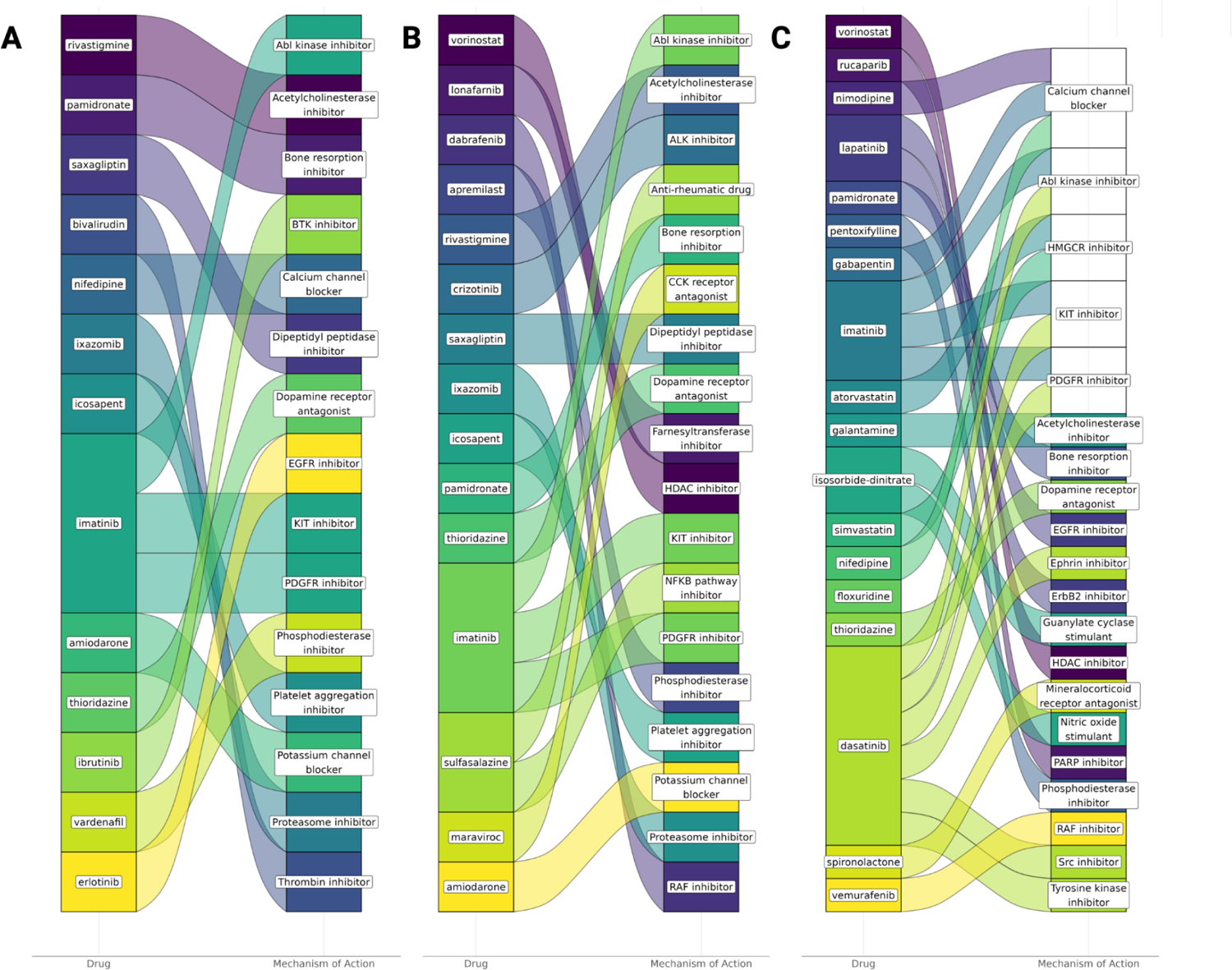
Alluvial plots of the top mechanism of action for identified GBM drug candidates. **A-C)** Alluvial plots for GBM drug candidates identified from the DESeq2, limma, or transfer learning disease-associated signatures, respectively. If the mechanism of action is colored white, this mechanism of action is shared by multiple drugs while the colored mechanism of action matches the drug with the same color. Note that these colors do not correspond to the same drug across disease-associated signatures or across cancer figures.

**Supplemental Figure 22:**
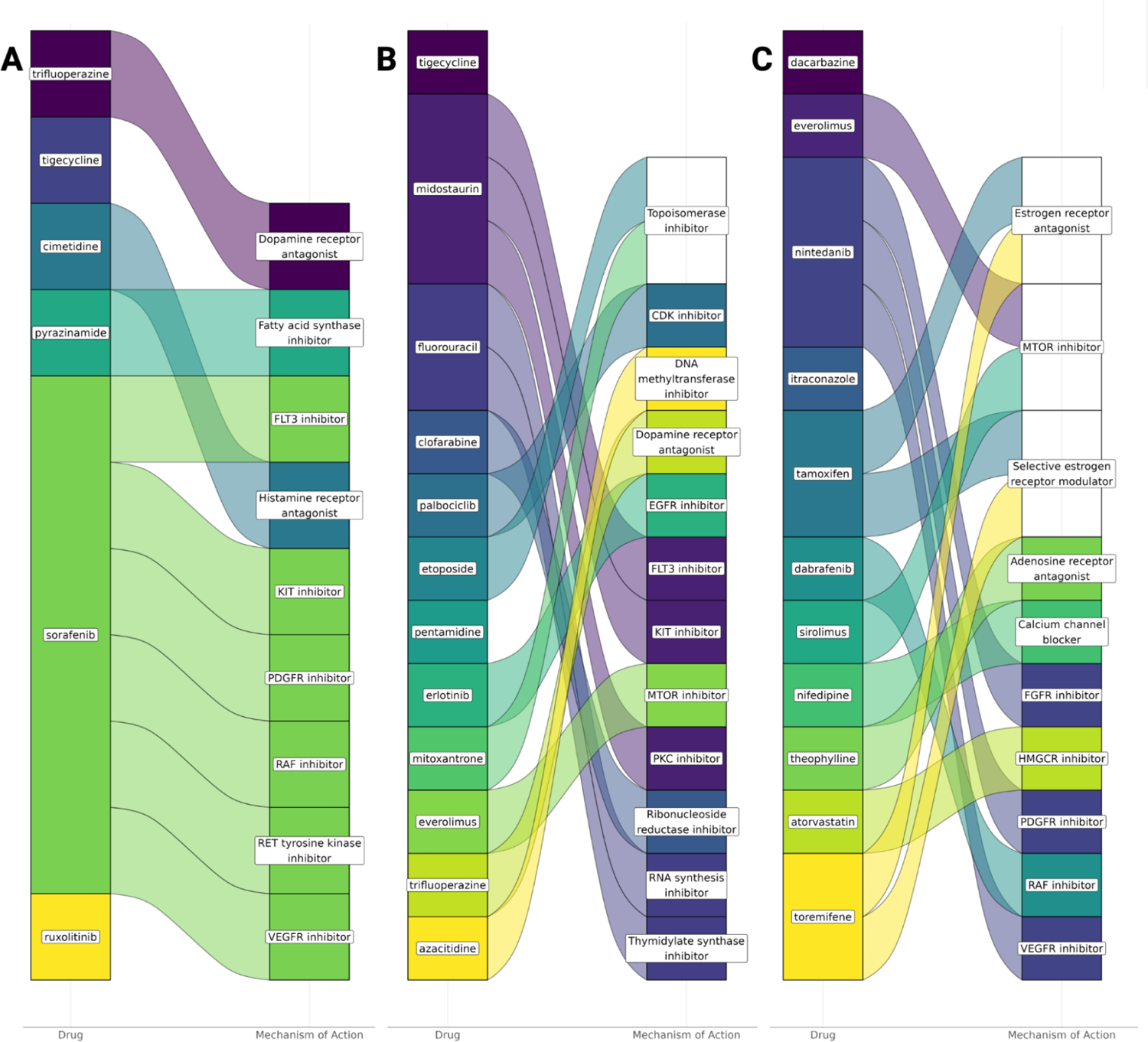
Alluvial plots of the top mechanism of action for identified LIHC drug candidates. **A-C)** Alluvial plots for GBM drug candidates identified from the DESeq2, limma, or transfer learning disease-associated signatures, respectively. If the mechanism of action is colored white, this mechanism of action is shared by multiple drugs while the colored mechanism of action matches the drug with the same color. Note that these colors do not correspond to the same drug across disease-associated signatures or across cancer figures.

**Supplemental Figure 23:**
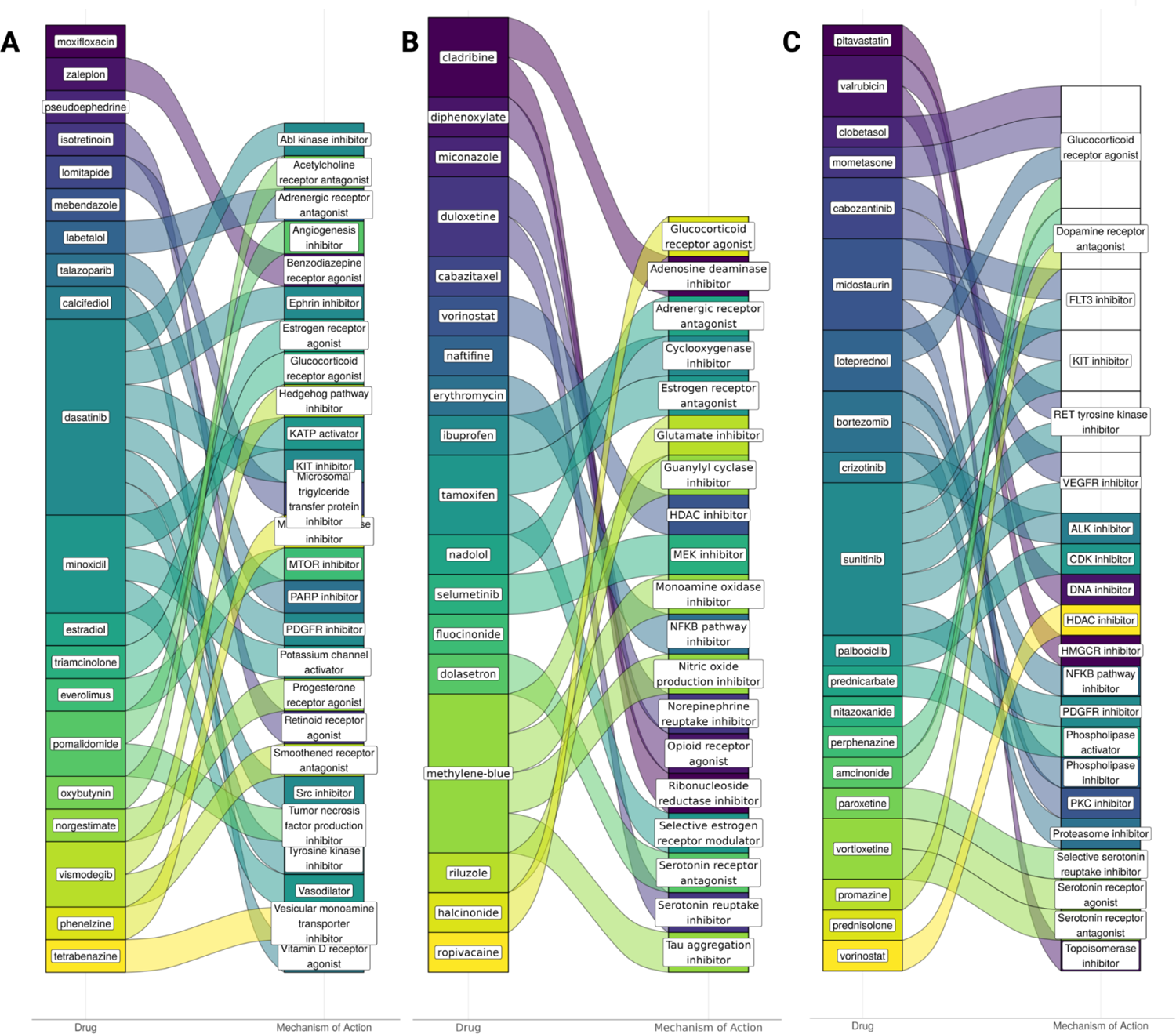
Alluvial plots of the top mechanism of action for identified LUAD drug candidates. **A-C)** Alluvial plots for GBM drug candidates identified from the DESeq2, limma, or transfer learning disease-associated signatures, respectively. If the mechanism of action is colored white, this mechanism of action is shared by multiple drugs while the colored mechanism of action matches the drug with the same color. Note that these colors do not correspond to the same drug across disease-associated signatures or across cancer figures.

**Supplemental Figure 24:**
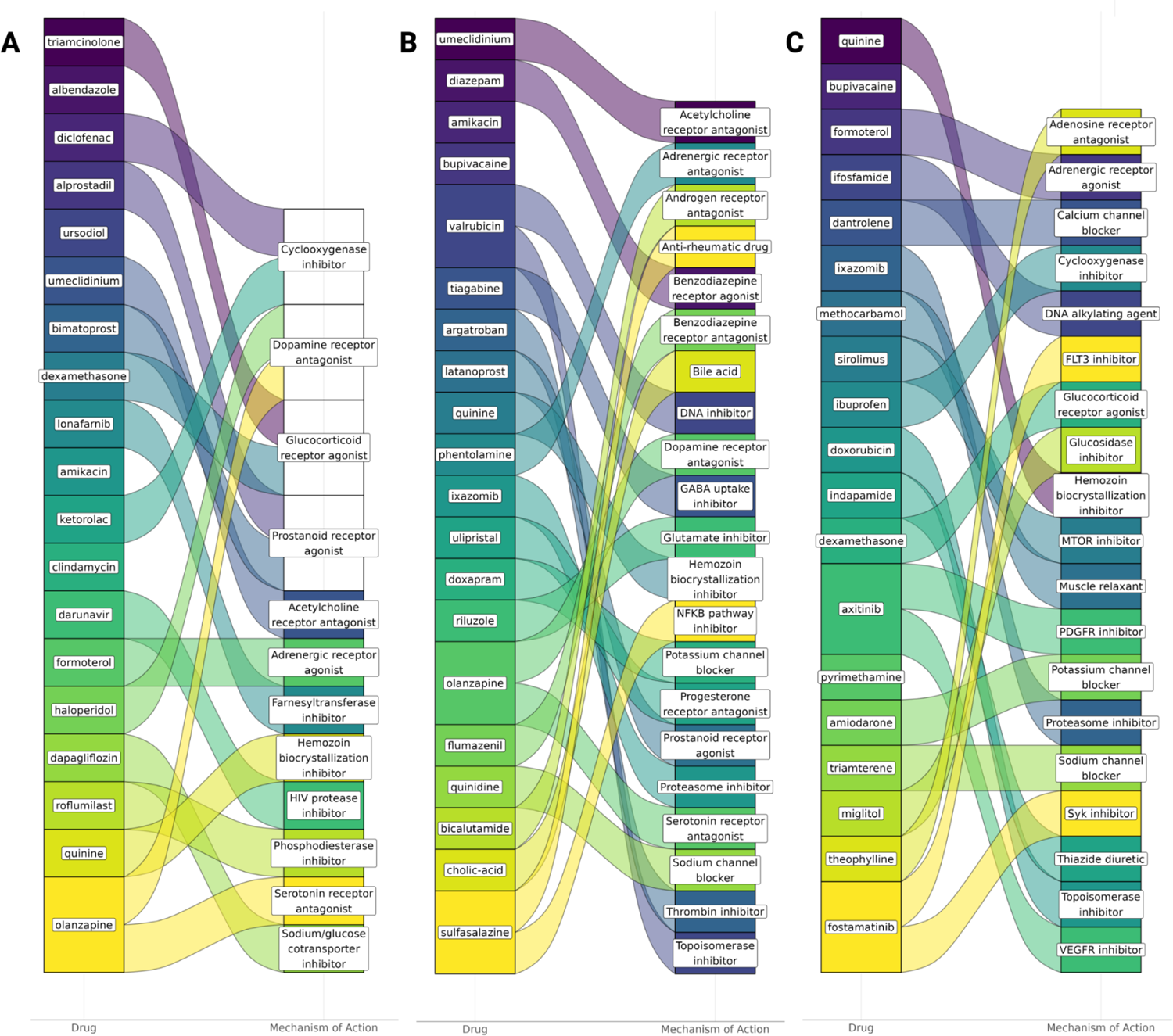
Alluvial plots of the top mechanism of action for identified PAAD drug candidates. **A-C)** Alluvial plots for GBM drug candidates identified from the DESeq2, limma, or transfer learning disease-associated signatures, respectively. If the mechanism of action is colored white, this mechanism of action is shared by multiple drugs while the colored mechanism of action matches the drug with the same color. Note that these colors do not correspond to the same drug across disease-associated signatures or across cancer figures.

**Supplemental Figure 25:**
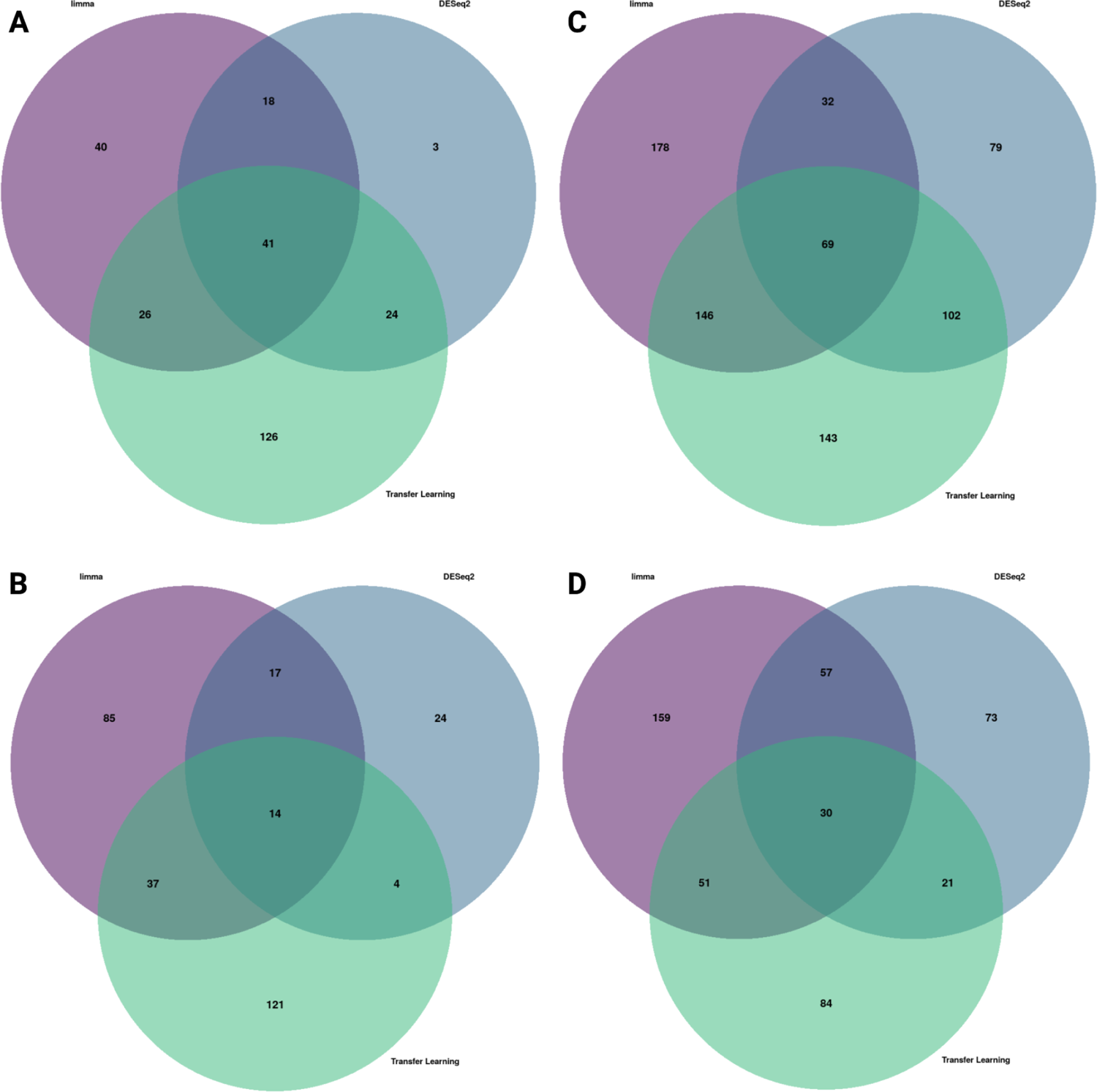
A-D) Venn diagrams of the overlap between the known drug targets of the top identified drug candidates for GBM, LIHC, LUAD, and PAAD, respectively.

**Supplemental Figure 26:**
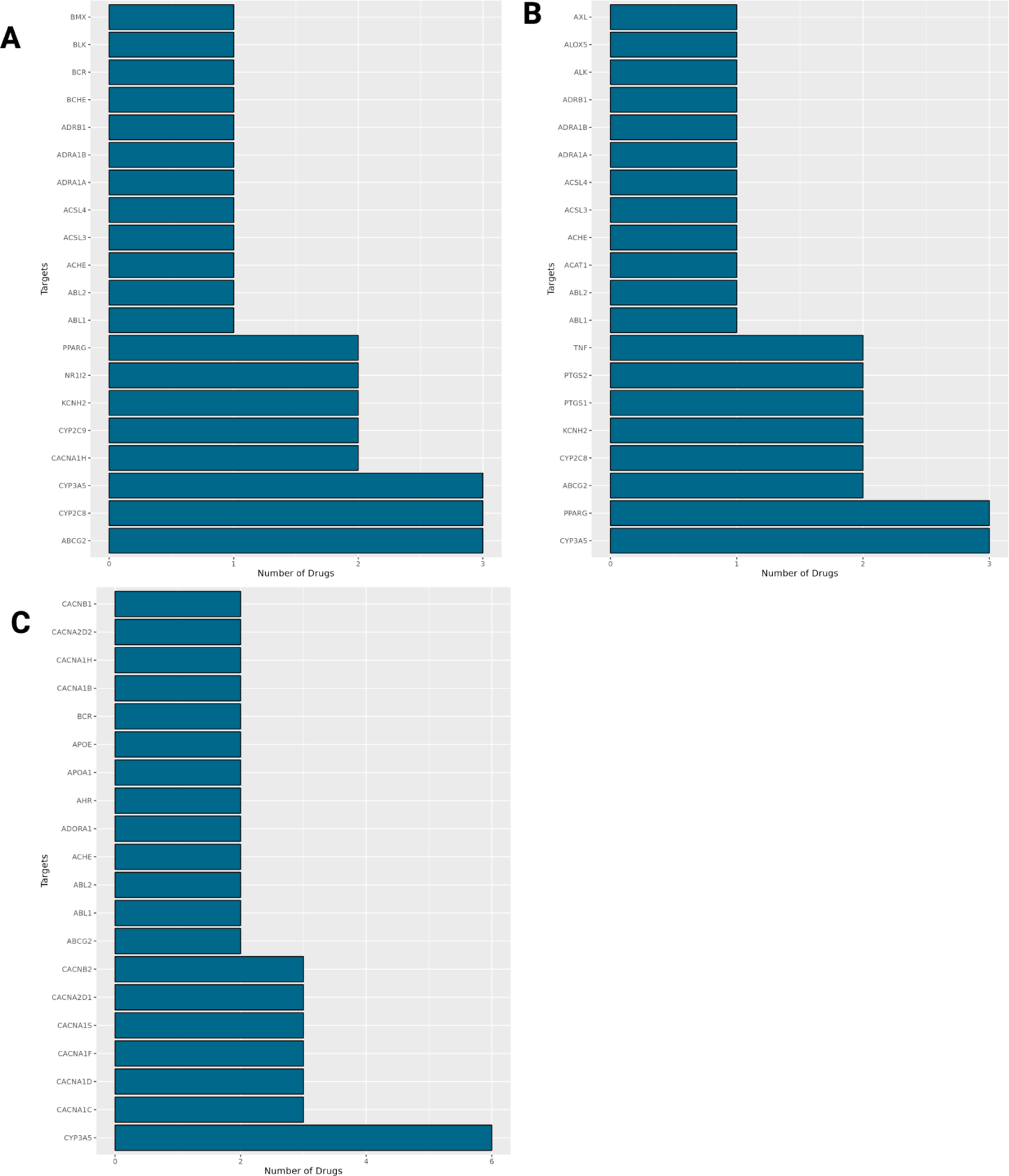
Bar plots of the top drug targets by disease-associated gene signature for GBM. **A)** Bar plots of the top known drug targets for identified GBM drug candidates from the DESeq2, limma, and transfer learning disease-associated gene signatures, respectively.

**Supplemental Figure 27:**
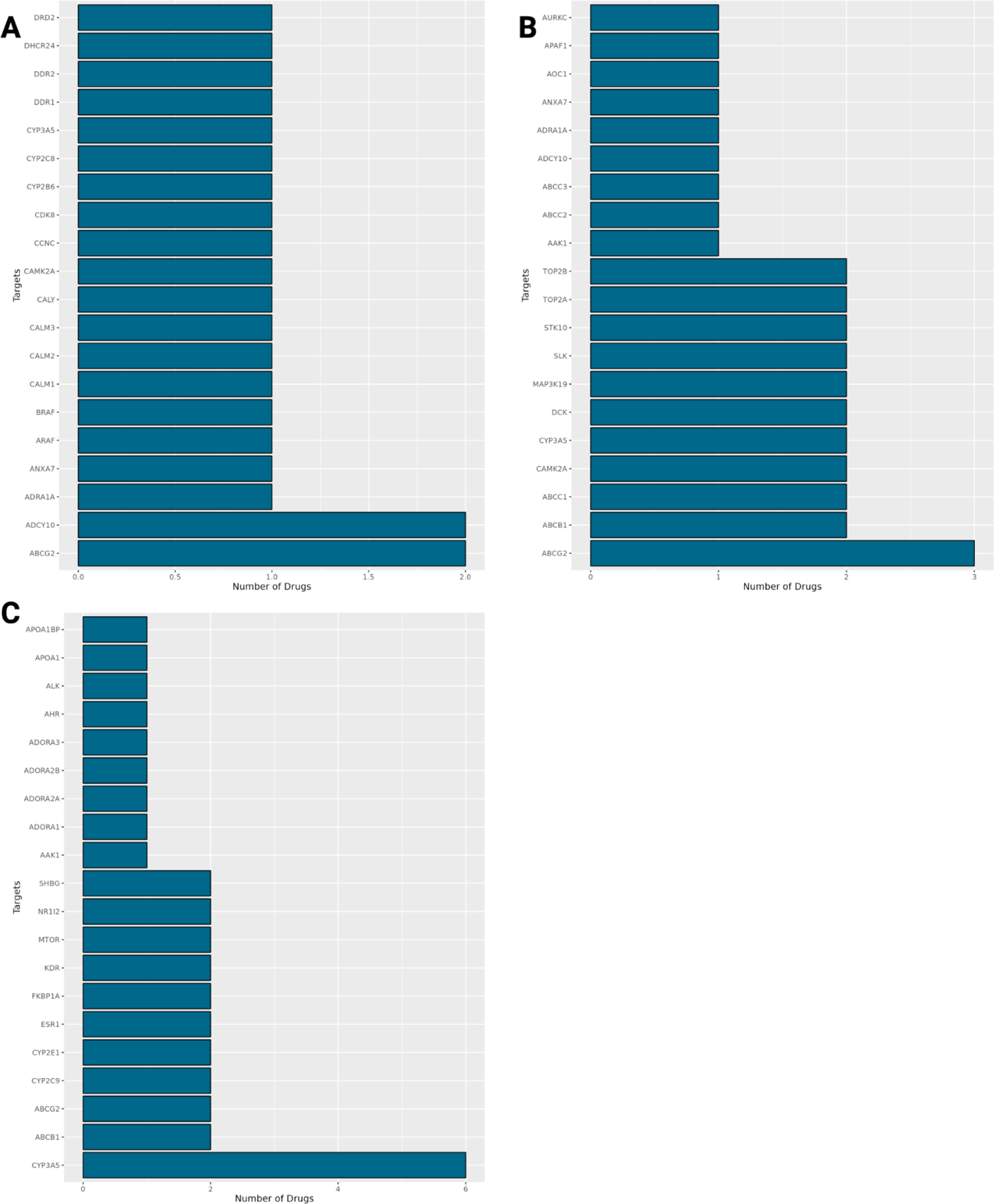
Bar plots of the top drug targets by disease-associated gene signature for LIHC. **A)** Bar plots of the top known drug targets for identified LIHC drug candidates from the DESeq2, limma, and transfer learning disease-associated gene signatures, respectively.

**Supplemental Figure 28:**
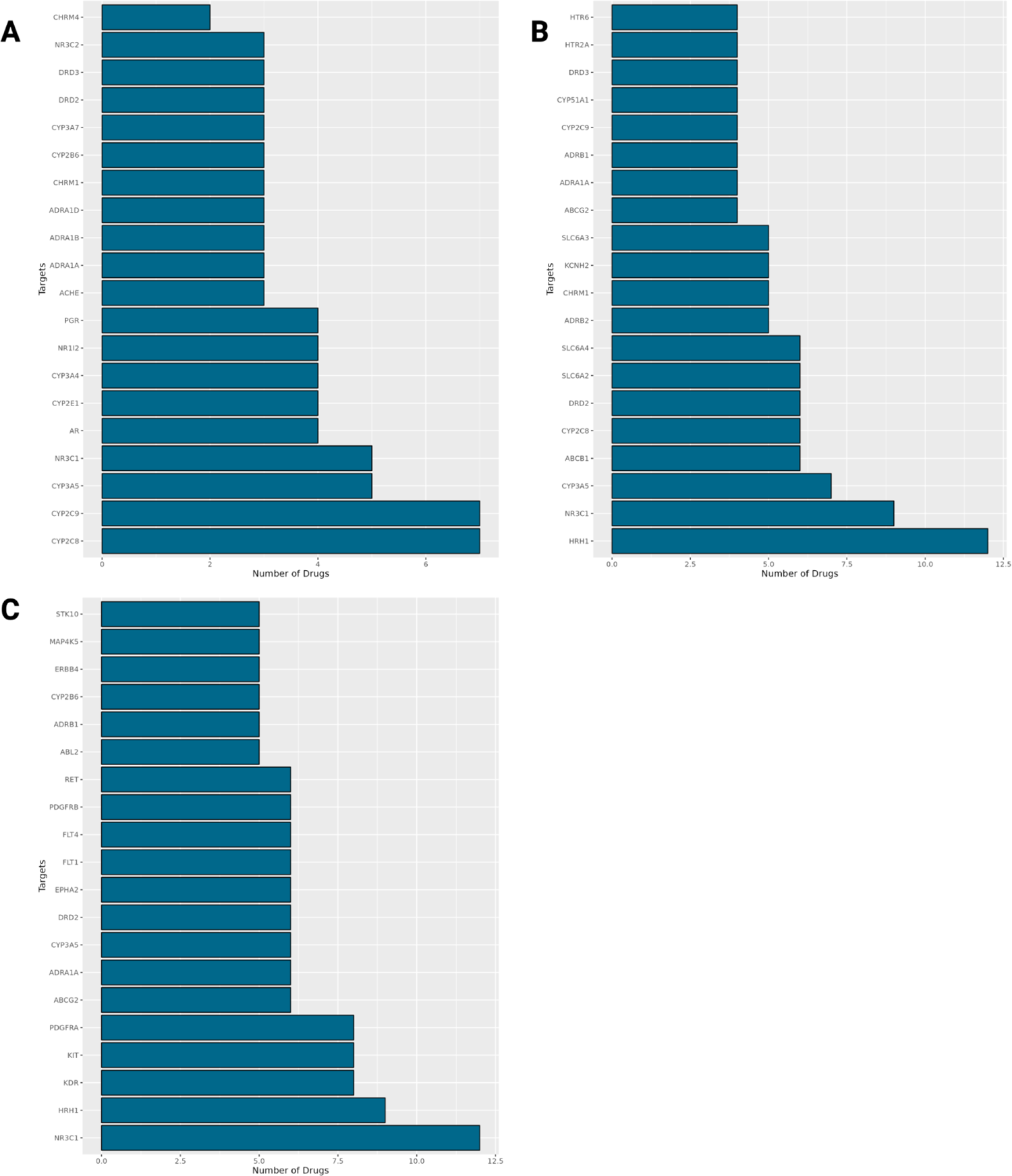
Bar plots of the top drug targets by disease-associated gene signature for LUAD. **A)** Bar plots of the top known drug targets for identified LUAD drug candidates from the DESeq2, limma, and transfer learning disease-associated gene signatures, respectively.

**Supplemental Figure 29:**
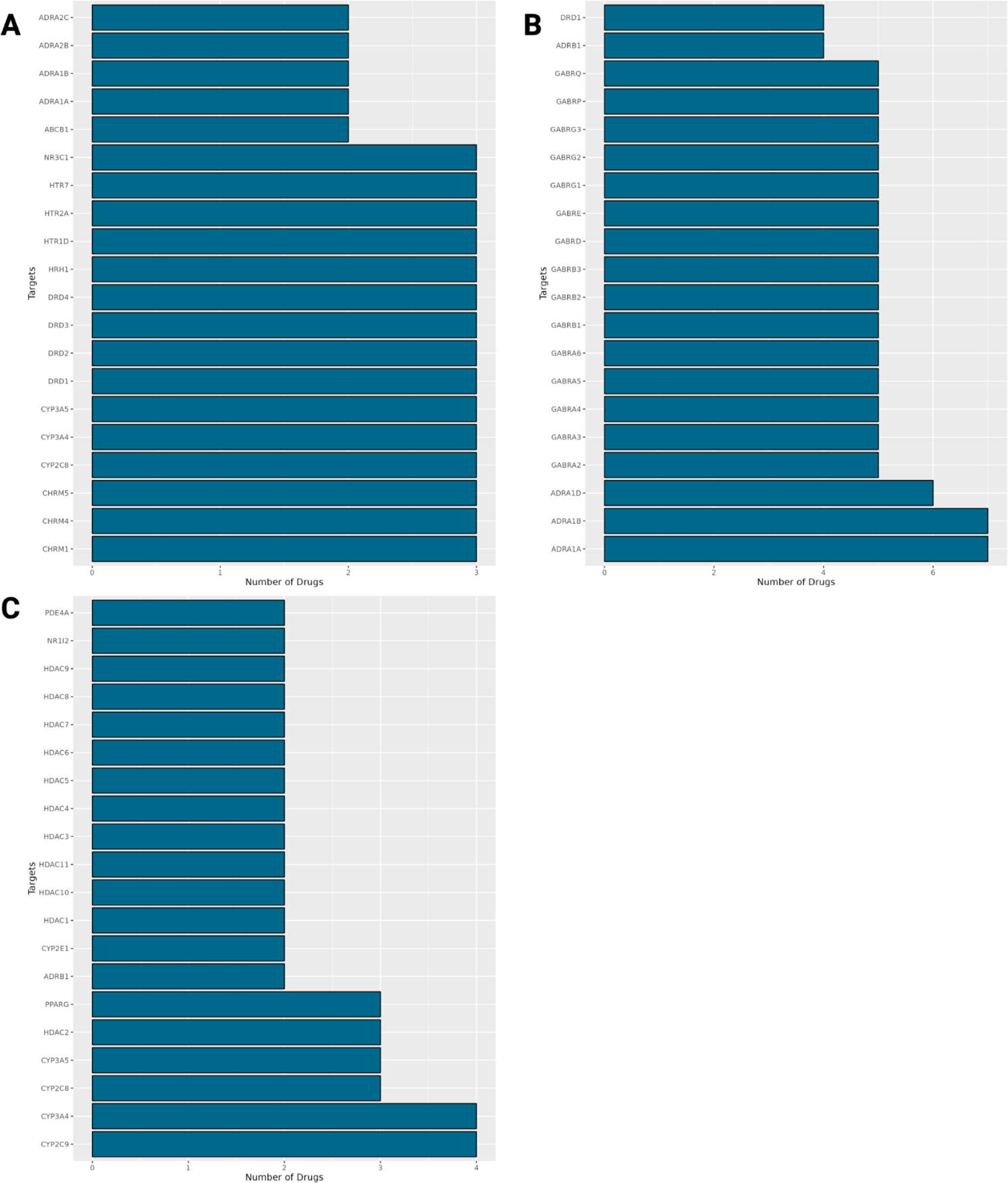
Bar plots of the top drug targets by disease-associated gene signature for PAAD. **A)** Bar plots of the top known drug targets for identified PAAD drug candidates from the DESeq2, limma, and transfer learning disease-associated gene signatures, respectively.

**Supplemental Figure 30:**
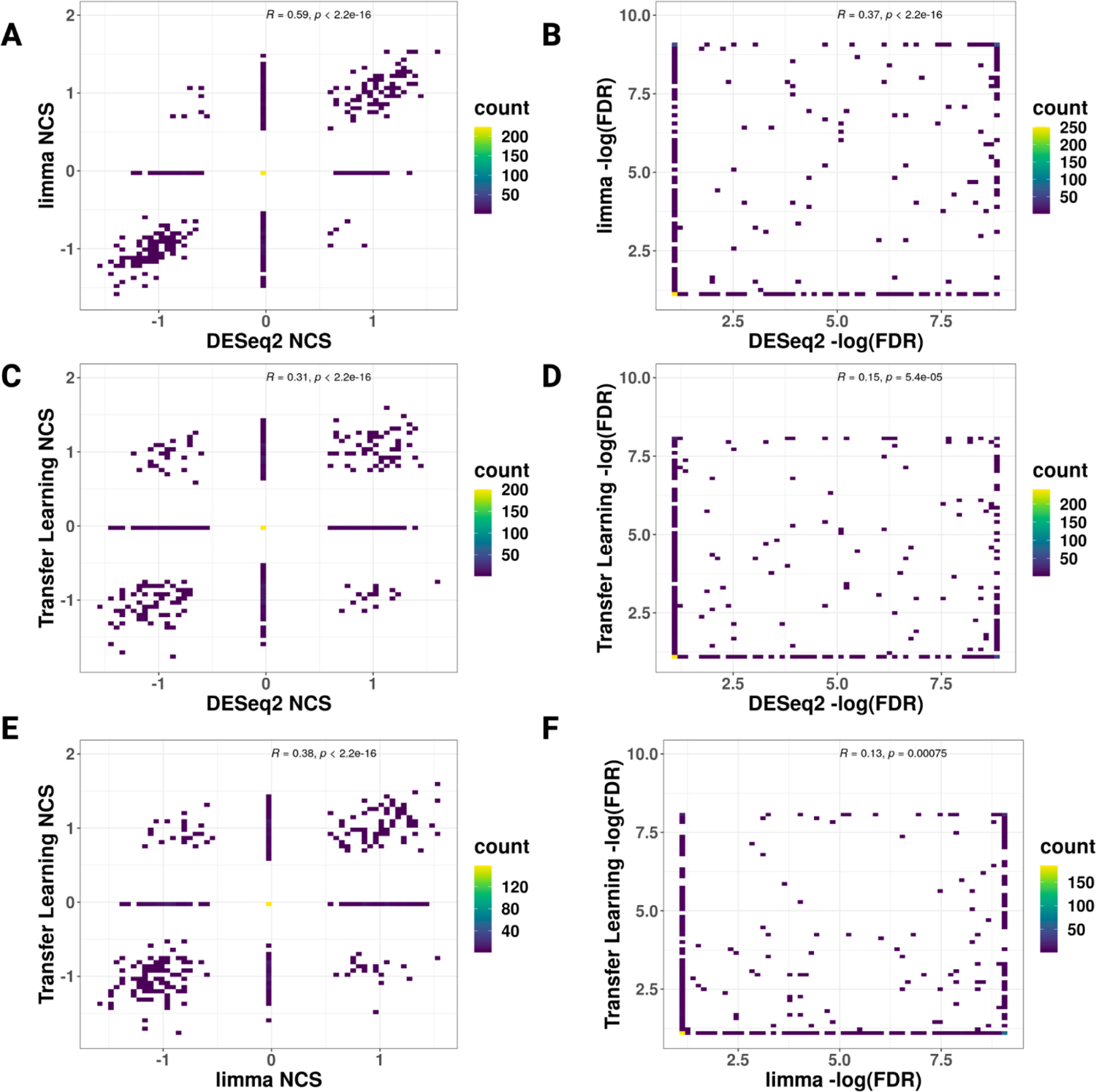
Signature reversion NCS and FDR scatter plots from each disease-associated gene signature for GBM. **A)** DESeq2 vs limma disease-associated gene signature reversion NCS score. **B)** DESeq2 vs limma disease-associated gene signature reversion FDR **C)** DESeq2 vs transfer learning disease-associated gene signature reversion NCS score **D)** DESeq2 vs transfer learning disease-associated gene signature reversion FDR **E)** limma vs. transfer learning disease-associated gene signature reversion NCS score **F)** limma vs. transfer learning disease-associated gene signature reversion FDR. Spearman correlation and p-value from linear regression models are also plotted on each panel.

**Supplemental Figure 31:**
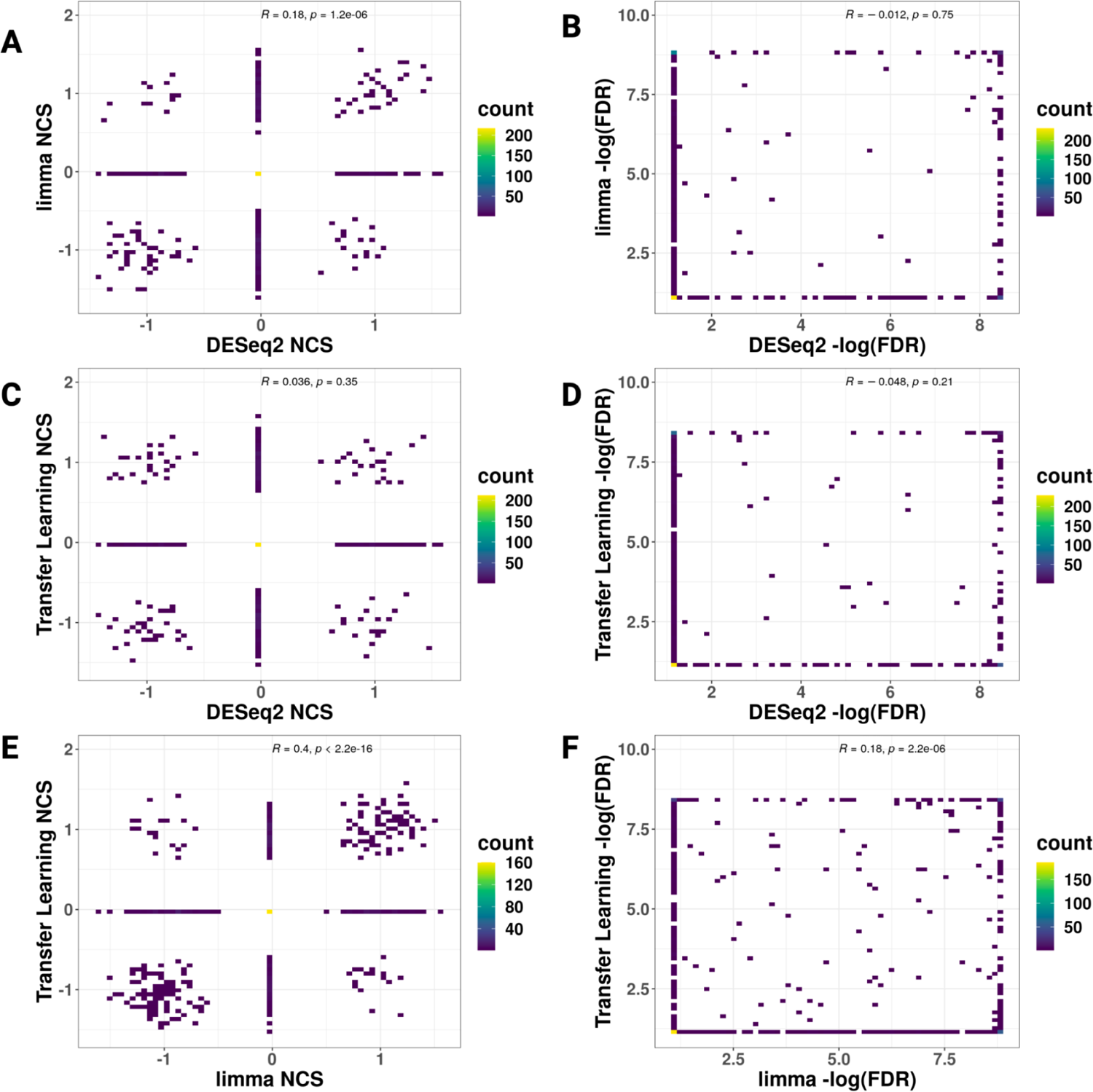
Signature reversion’s NCS and FDR scatter plots for all methods for LIHC. A) DESeq2 vs limma disease-associated gene signature reversion NCS score. **B)** DESeq2 vs limma disease-associated gene signature reversion FDR **C)** DESeq2 vs transfer learning disease-associated gene signature reversion NCS score **D)** DESeq2 vs transfer learning disease-associated gene signature reversion FDR **E)** limma vs. transfer learning disease-associated gene signature reversion NCS score **F)** limma vs. transfer learning disease-associated gene signature reversion FDR. Spearman correlation and p-value from linear regression models are also plotted on each panel.

**Supplemental Figure 32:**
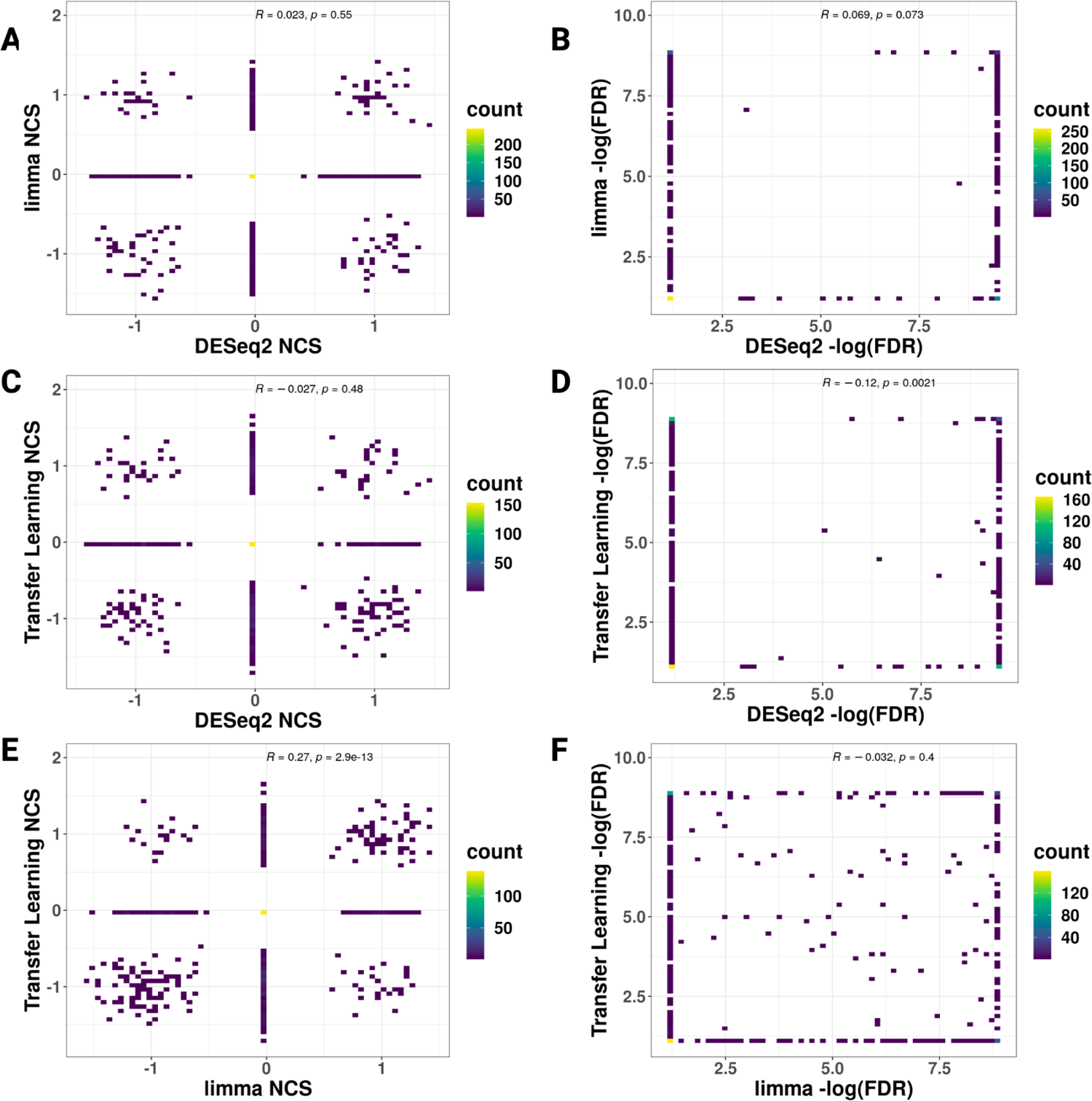
Signature reversion’s NCS and FDR scatter plots for all methods. **A)** DESeq2 vs limma disease-associated gene signature reversion NCS score. **B)** DESeq2 vs limma disease-associated gene signature reversion FDR **C)** DESeq2 vs transfer learning disease-associated gene signature reversion NCS score **D)** DESeq2 vs transfer learning disease-associated gene signature reversion FDR **E)** limma vs. transfer learning disease-associated gene signature reversion NCS score **F)** limma vs. transfer learning disease-associated gene signature reversion FDR. Spearman correlation and p-value from linear regression models are also plotted on each panel.

**Supplemental Figure 33:**
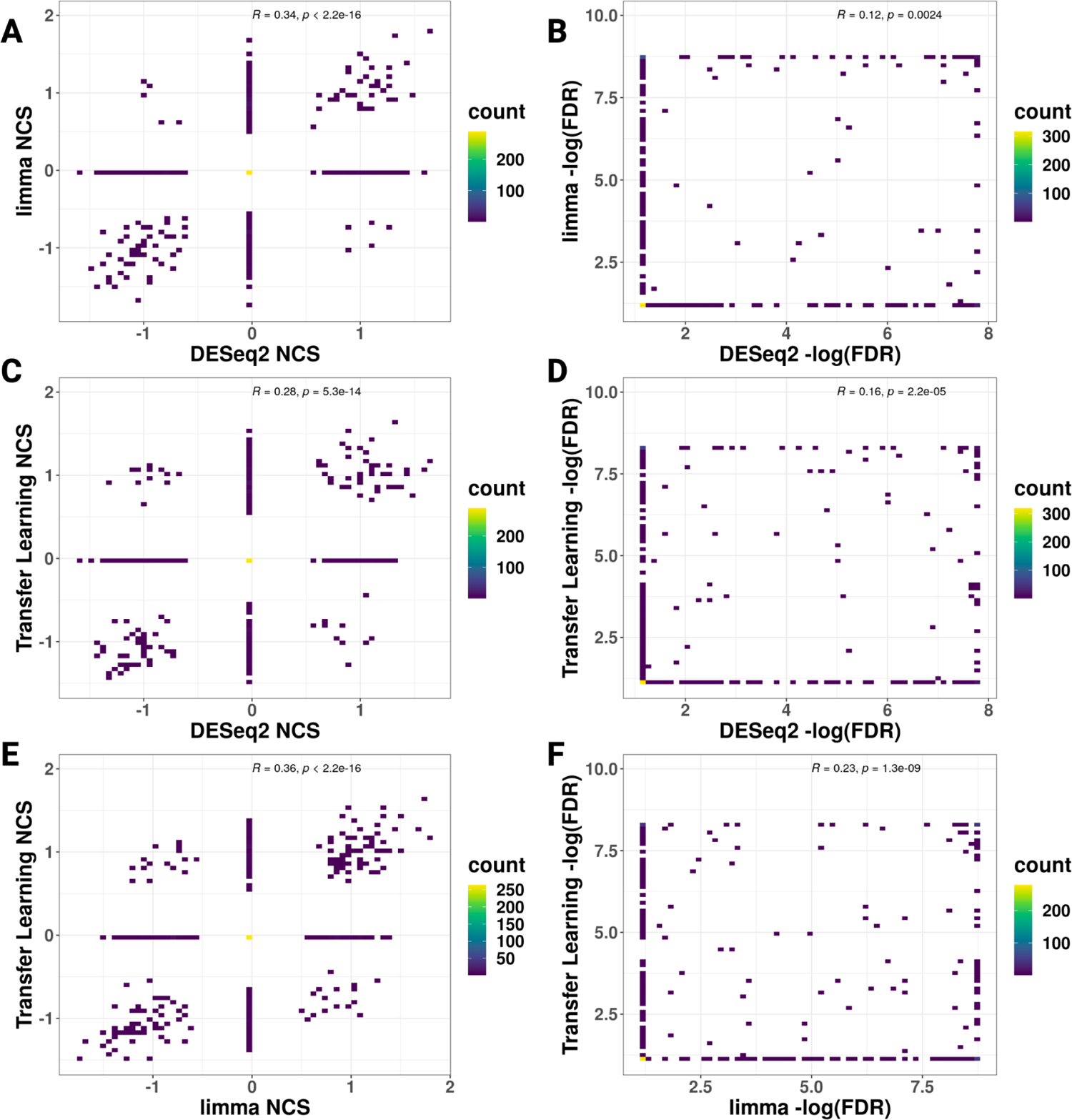
Signature reversion’s NCS and FDR scatter plots for all methods. **A)** DESeq2 vs limma disease-associated gene signature reversion NCS score. **B)** DESeq2 vs limma disease-associated gene signature reversion FDR **C)** DESeq2 vs transfer learning disease-associated gene signature reversion NCS score **D)** DESeq2 vs transfer learning disease-associated gene signature reversion FDR **E)** limma vs. transfer learning disease-associated gene signature reversion NCS score **F)** limma vs. transfer learning disease-associated gene signature reversion FDR. Spearman correlation and p-value from linear regression models are also plotted on each panel.

**Supplemental Figure 34:**
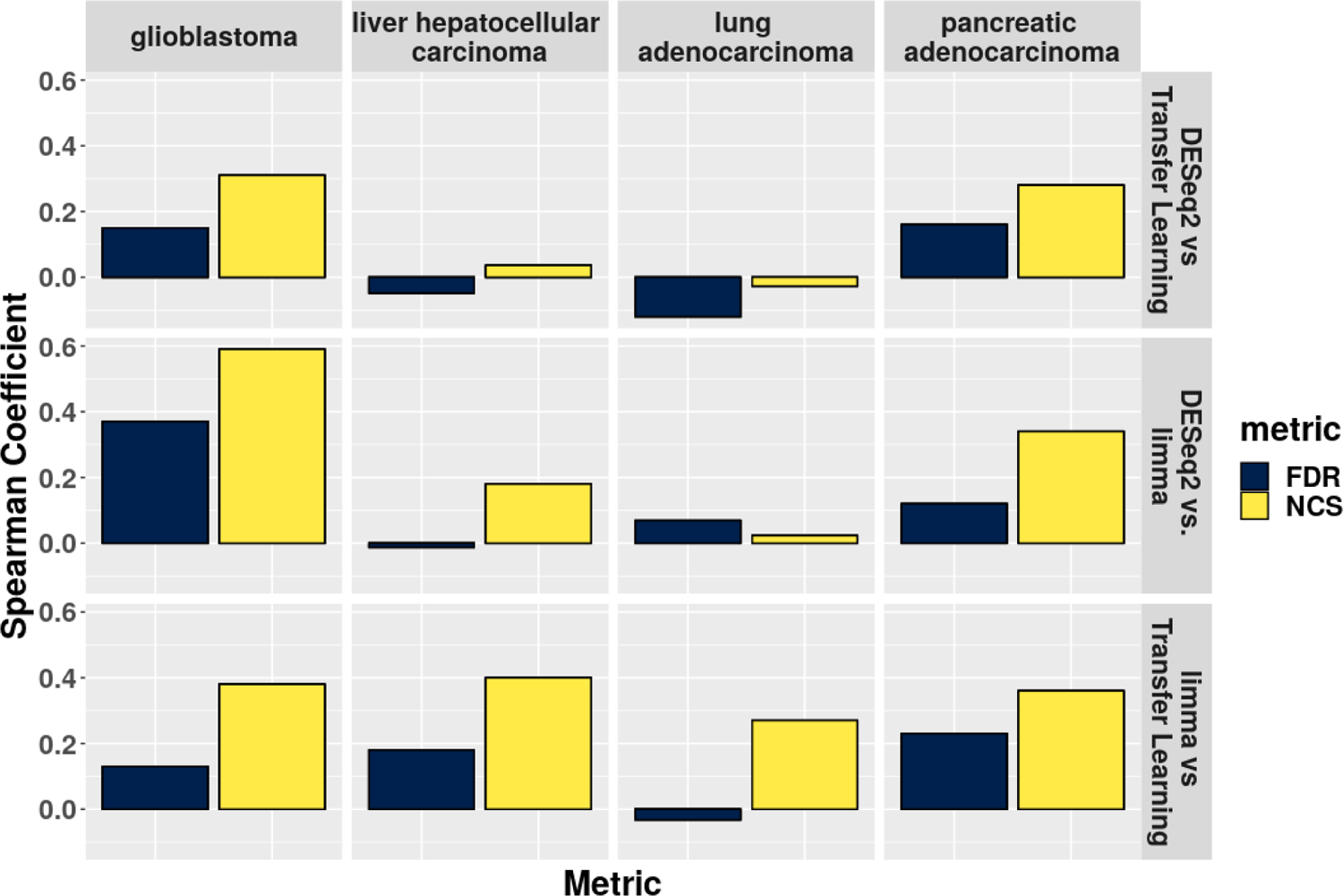
A bar plot of the Spearman correlation of the normalized connectivity score (NCS) and the false discovery (FDR) between the different disease-associated gene signature reversion results across the different cancers.

**Supplemental Figure 35:**
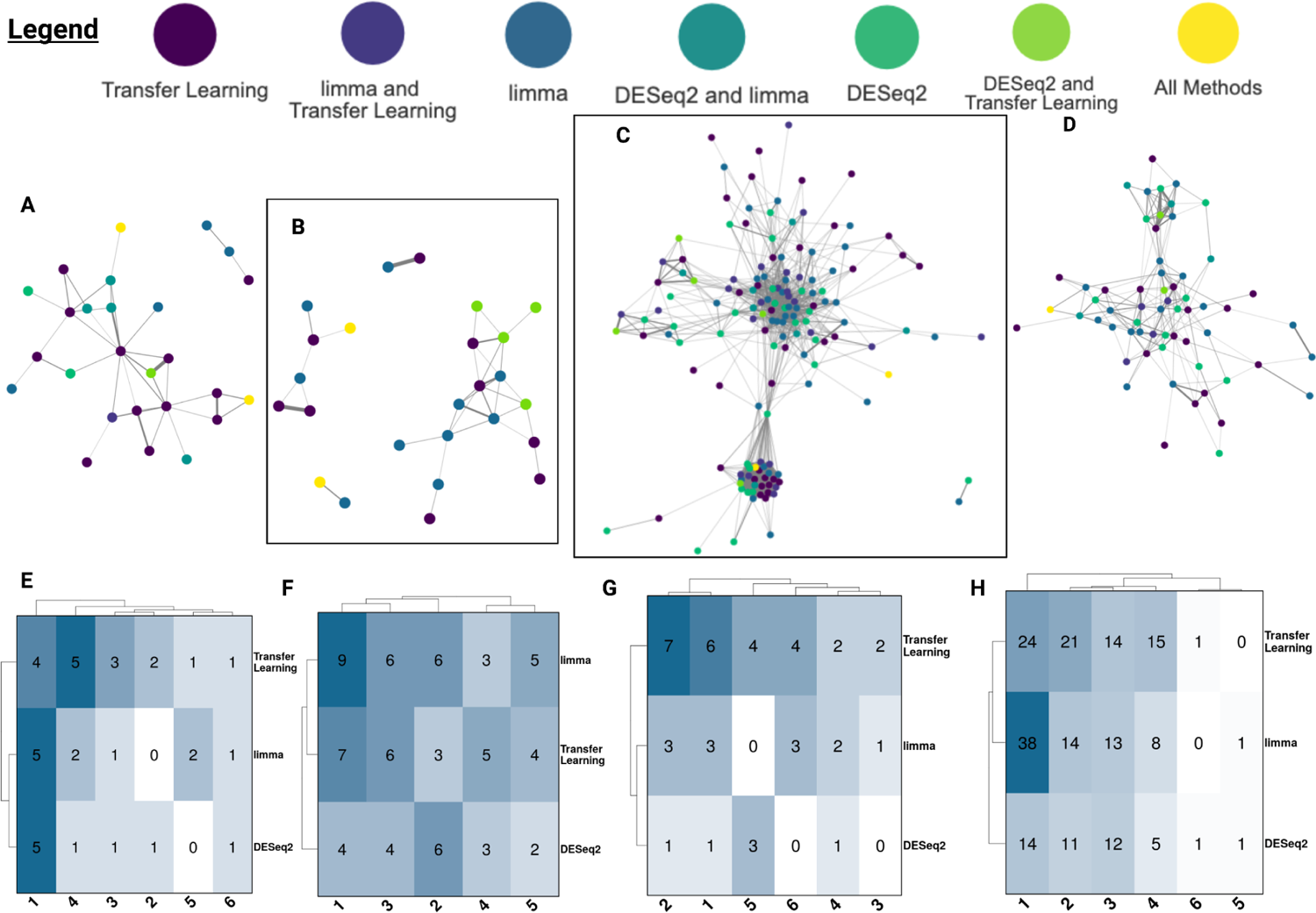
Drug-drug similarity networks - drug structure. **A-D)** Drug-drug similarity networks based on the Tanimoto coefficient of SMILES drug structures where each node is a candidate colored by the disease-associated gene signature used to identify that candidate and the top 90% of edges displayed and weighted by cosine similarity for **A)** GBM candidates (GI1 profiles), **B)** LIHC cancer candidates (HEPG2 profiles), **C)** LUAD candidates (A529 LINCS profiles), and **D)** PAAD candidates (YPAC profiles). Heatmaps of the composition of the Leiden communities in the drug-drug similarity networks for **E)** GBM candidates (GI1 profiles), **F)** LIHC candidates (HEPG2 profiles), **G)** LUAD candidates (A529 LINCS profiles), and **H)** PAAD candidates (YPAC profiles).

**Supplemental Figure 36:**
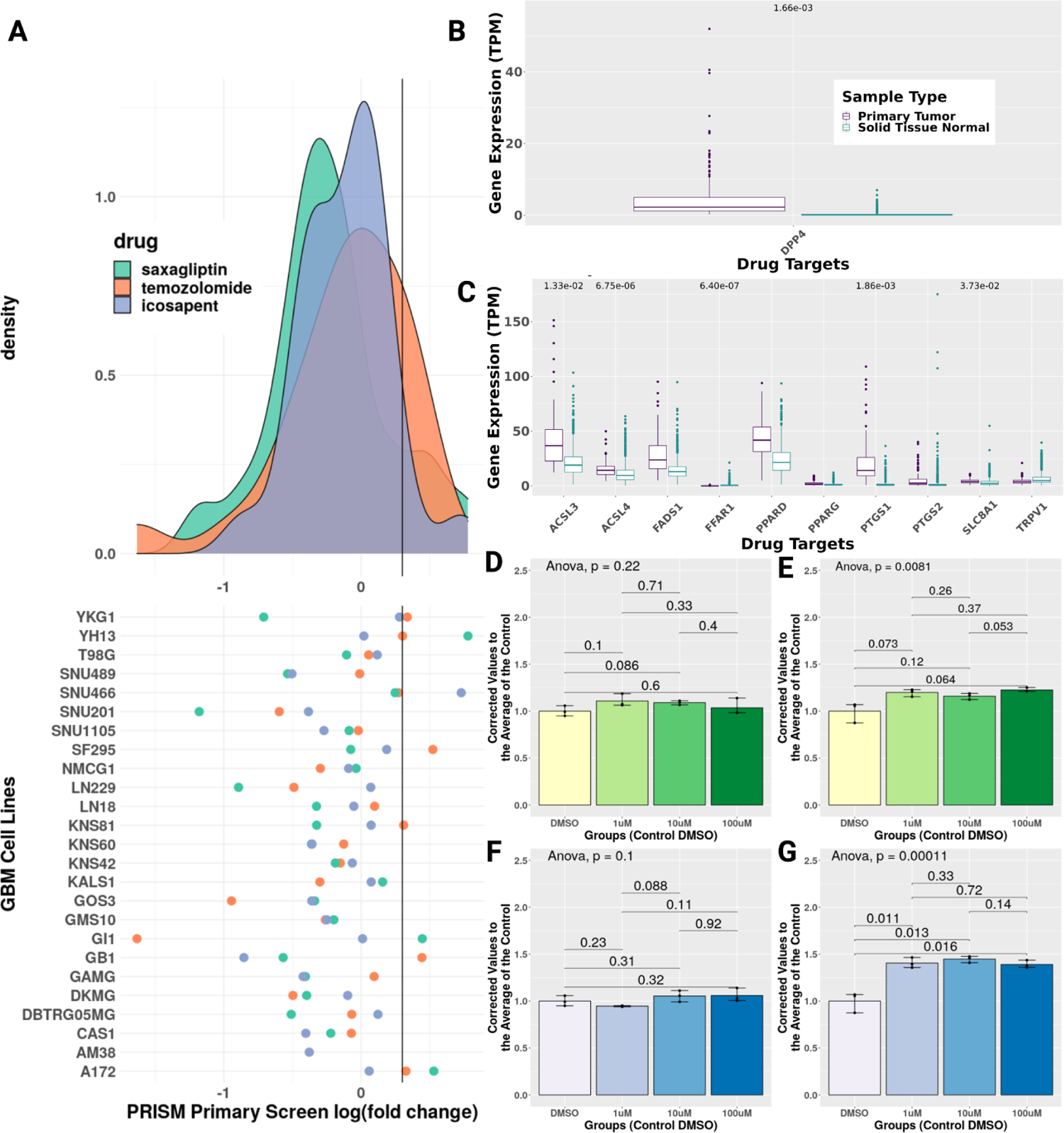
Round 1 results for saxagliptin and icosapent. **A)** Density plot of candidates and TMZ log fold change from PRISM primary screen and dot plot of primary screen results for GBM cell lines. **B+C)** Boxplot of candidate’s drug target expression in tumor tissue and control brain tissue for saxagliptin **B** and icosapent **C**. **D + E)** CellTiter-Glo growth assay results for saxagliptin in U251 GBM and NHA cell lines, respectively. **F + G)** CellTiter-Glo growth assay results for icosapent in U251 GBM and NHA cell lines, respectively.

**Supplemental Figure 37:**
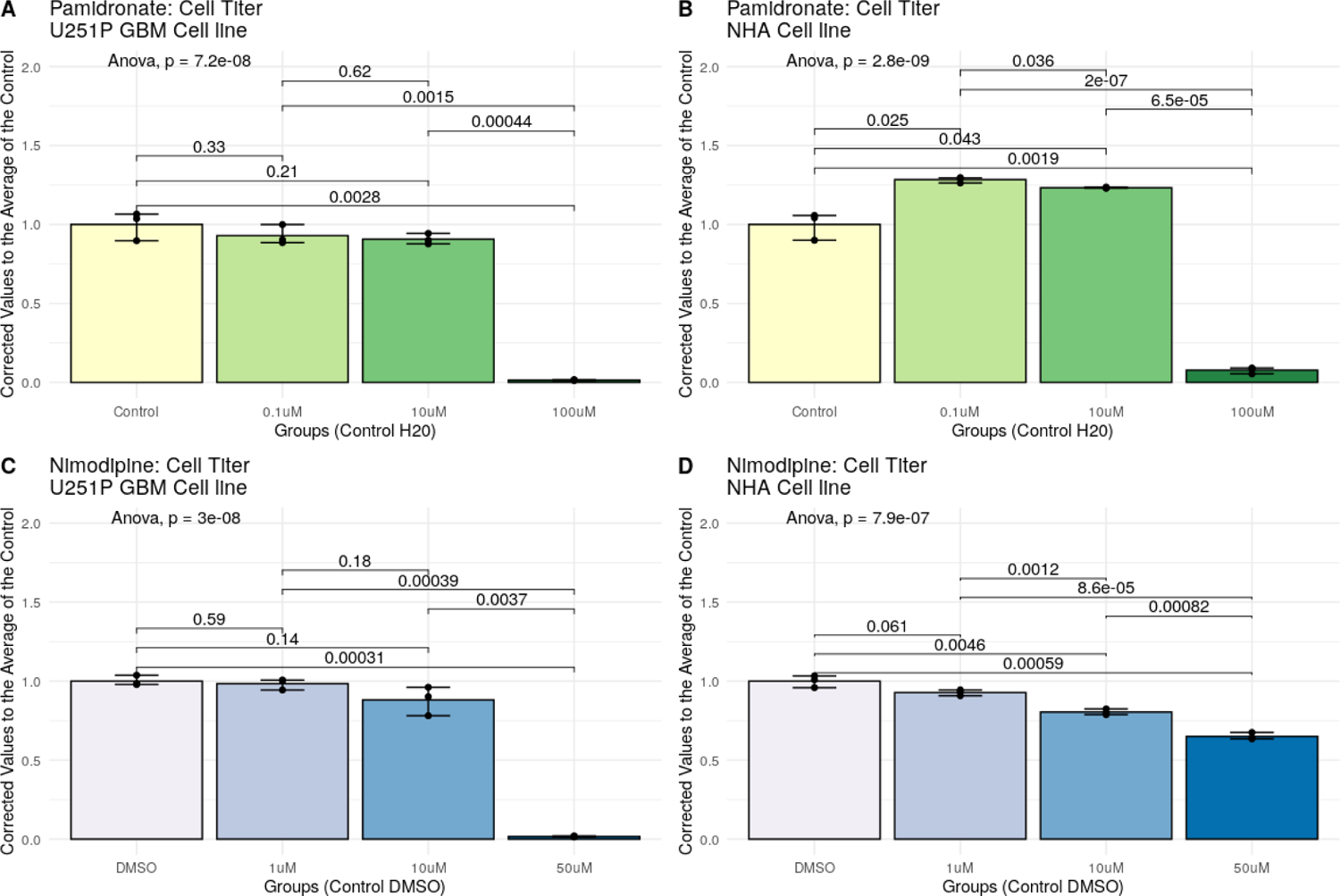
CellTiter-Glo Round 1 results for pamidronate and nimodipine. **A + B)** CellTiter-Glo viability assay results for pamidronate in U251 GBM and NHA cell lines, respectively. **C + D)** CellTiter-Glo viability assay results for nimodipine in U251 GBM and NHA cell lines, respectively.

**Supplemental Figure 38:**
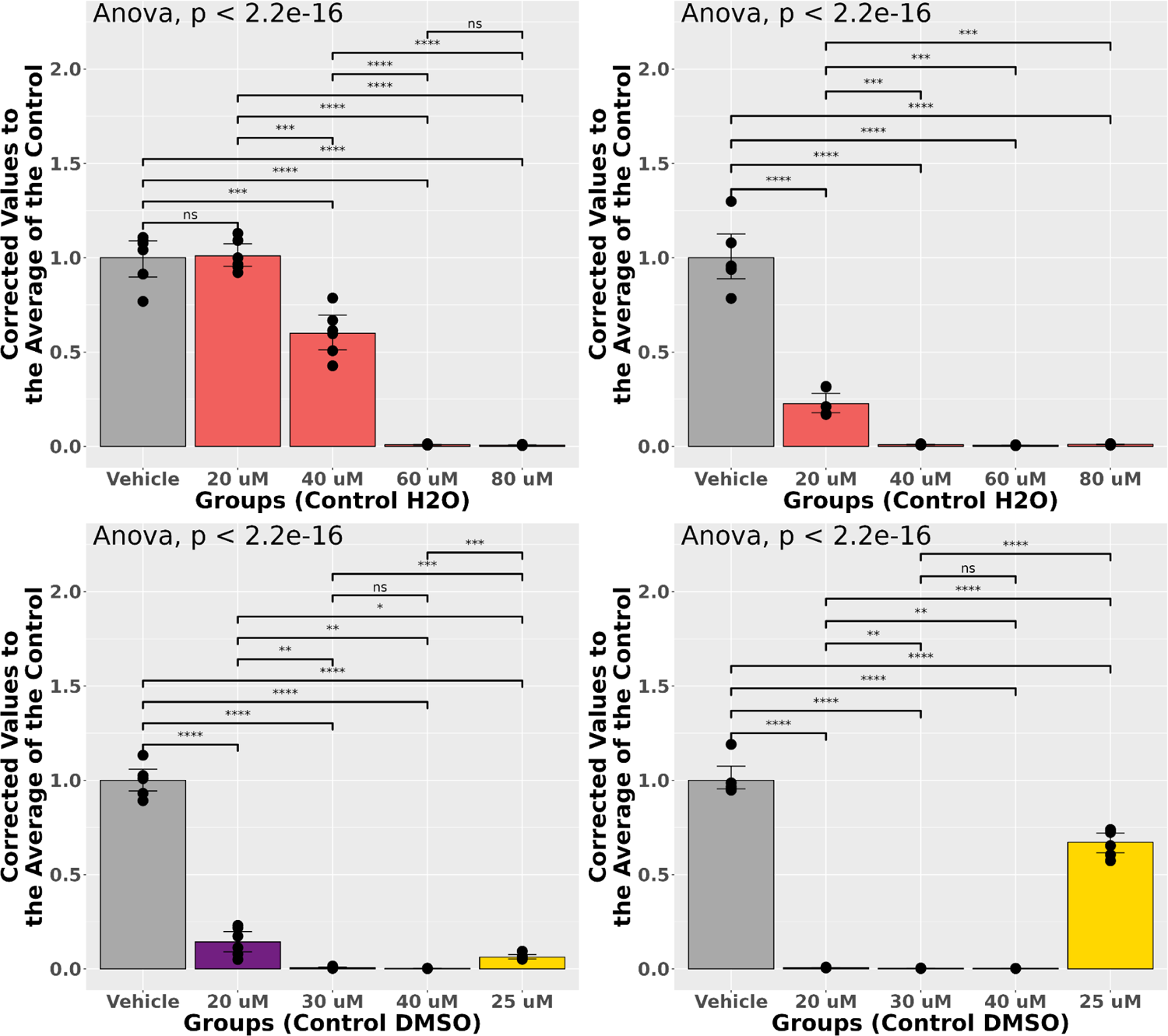
CellTiter-Glo Round 3 results for pamidronate and nimodipine. **A + B)** CellTiter-Glo viability assay results for pamidronate in U251 GBM and JX39 cell lines, respectively. **C + D)** CellTiter-Glo viability assay results for nimodipine in U251 GBM and JX39 cell lines, respectively. *P < 0.05, **P < 0.01, ***P < 0.001, ****P < 0.0001 ANOVA with Bonferroni corrected t-tests for pairwise comparisons.

**Supplemental Figure 39:**
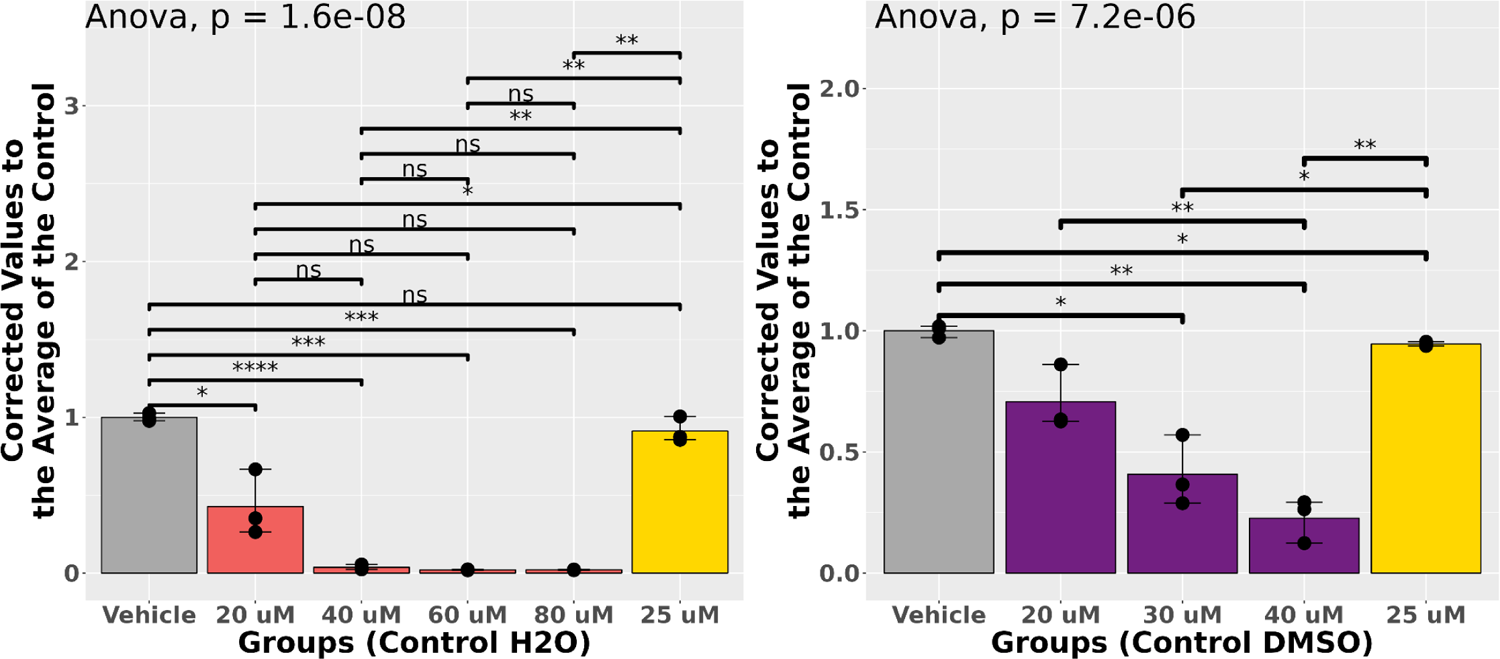
CellTiter-Glo round 2 results for pamidronate and nimodipine in NHA cell line, respectively. *P < 0.05, **P < 0.01, ***P < 0.001, ****P < 0.0001 ANOVA with Bonferroni corrected t-tests for pairwise comparisons.

**Supplemental Figure 40:**
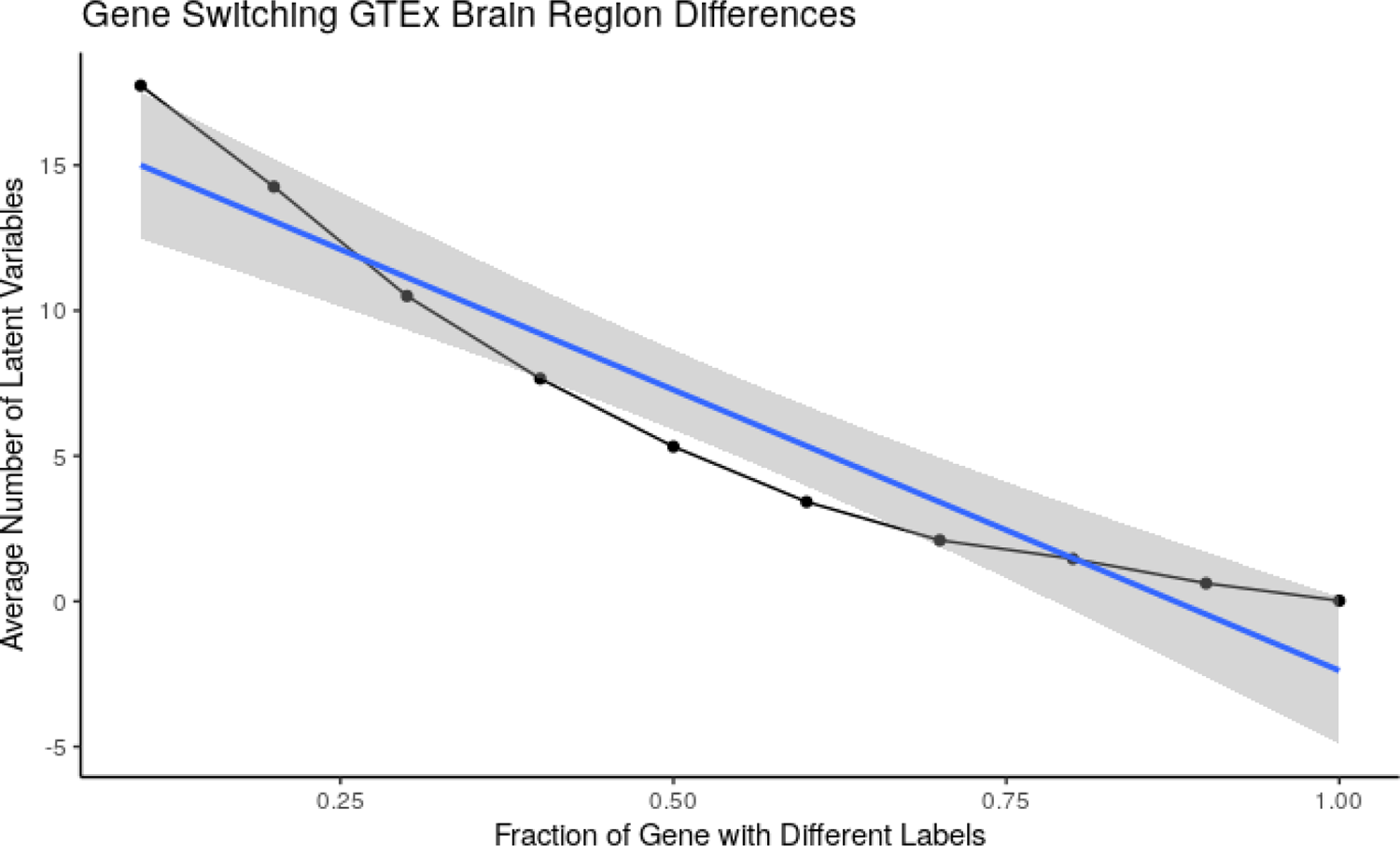
Gene label switching test for transfer learning approach. GTEx brain frontal cortex and cerebellar hemisphere regions were compared with the transfer learning approach by determining latent variables that were different between these two regions. A fraction of the genes had their labels switched with another gene. This was done in 10% intervals starting at 10% and ending at 100% of all the genes. This process was done 50 times at each interval. The average number of latent variables significant for each interval was calculated (adj. p-value < 0.05; fold change of 0.05). A linear regression model was used to determine if there was an influence of the gene label and the number of latent variables (R^2^ = 0.9066 & p-value = 1.346e-05).

