## Supplementary material for "Signature reversion of three disease-associated gene signatures prioritizes cancer drug repurposing candidates": FILE4_computer_system_and_packages_info_dockers_conda

### 221221_rstudio_tf_dr_v3_singularity


##### Jennifer Fisher


### 12/21/2022

```
GitHub Repo:
Transfer_Learning_R03

Docker:
rstudio_tf_dr_v3

Directory of operations:
/data/project/lasseigne_lab/JLF_scratch/Transfer_Learning_R03 (singularity)
```

```
sessionInfo()
```

```
## R version 4.1.2 (2021-11-01)
## Platform: x86_64-pc-linux-gnu (64-bit)
## Running under: Ubuntu 20.04.3 LTS
##
## Matrix products: default
## BLAS/LAPACK: /usr/lib/x86_64-linux-gnu/openblas-pthread/libopenblasp-r0.3.8.so
##
## locale:
##  [1] LC_CTYPE=en_US.UTF-8       LC_NUMERIC=C
##  [3] LC_TIME=en_US.UTF-8        LC_COLLATE=en_US.UTF-8
##  [5] LC_MONETARY=en_US.UTF-8    LC_MESSAGES=C
##  [7] LC_PAPER=en_US.UTF-8       LC_NAME=C
##  [9] LC_ADDRESS=C               LC_TELEPHONE=C
## [11] LC_MEASUREMENT=en_US.UTF-8 LC_IDENTIFICATION=C
##
## attached base packages:
## [1] stats     graphics  grDevices utils     datasets  methods   base
##
## loaded via a namespace (and not attached):
##  [1] digest_0.6.29   R6_2.5.1        jsonlite_1.7.2  magrittr_2.0.1
##  [5] evaluate_0.14   rlang_0.4.12    stringi_1.7.6   jquerylib_0.1.4
##  [9] bslib_0.3.1     rmarkdown_2.11  tools_4.1.2     stringr_1.4.0
## [13] xfun_0.28       yaml_2.2.1      fastmap_1.1.0   compiler_4.1.2
## [17] htmltools_0.5.2 knitr_1.36      sass_0.4.0
```

```
# Listing packages
installed.packages()[,c(1,3)]
```

```
##                         Package                  Version
## abind                   "abind"                  "1.4-5"
## affy                    "affy"                   "1.72.0"
## affyio                  "affyio"                 "1.64.0"
## amap                    "amap"                   "0.8-18"
## annotate                "annotate"               "1.72.0"
## AnnotationDbi           "AnnotationDbi"          "1.56.2"
## AnnotationFilter        "AnnotationFilter"       "1.18.0"
## AnnotationHub           "AnnotationHub"          "3.0.2"
## AnVIL                   "AnVIL"                  "1.6.0"
## apcluster               "apcluster"              "1.4.8"
## ape                     "ape"                    "5.5"
## aplot                   "aplot"                  "0.1.1"
## askpass                 "askpass"                "1.1"
## assertthat              "assertthat"             "0.2.1"
## backports               "backports"              "1.4.0"
## base64enc               "base64enc"              "0.1-3"
## BgeeDB                  "BgeeDB"                 "2.20.0"
## BH                      "BH"                     "1.75.0-0"
## Biobase                 "Biobase"                "2.54.0"
## BiocFileCache           "BiocFileCache"          "2.2.0"
## BiocGenerics            "BiocGenerics"           "0.40.0"
## BiocIO                  "BiocIO"                 "1.4.0"
## BiocManager             "BiocManager"            "1.30.16"
## BiocParallel            "BiocParallel"           "1.28.2"
## BiocVersion             "BiocVersion"            "3.14.0"
## biomaRt                 "biomaRt"                "2.50.1"
## Biostrings              "Biostrings"             "2.62.0"
## bit                     "bit"                    "4.0.4"
## bit64                   "bit64"                  "4.0.5"
## bitops                  "bitops"                 "1.0-7"
## biwt                    "biwt"                   "1.0"
## blob                    "blob"                   "1.2.2"
## brew                    "brew"                   "1.0-6"
## brio                    "brio"                   "1.1.2"
## broom                   "broom"                  "0.7.10"
## bslib                   "bslib"                  "0.3.1"
## cachem                  "cachem"                 "1.0.6"
## callr                   "callr"                  "3.7.0"
## car                     "car"                    "3.0-12"
## carData                 "carData"                "3.0-4"
## caret                   "caret"                  "6.0-90"
## caTools                 "caTools"                "1.18.2"
## cellranger              "cellranger"             "1.1.0"
## circlize                "circlize"               "0.4.13"
## cli                     "cli"                    "3.1.0"
## clipr                   "clipr"                  "0.7.1"
## clue                    "clue"                   "0.3-60"
## clusterProfiler         "clusterProfiler"        "4.0.5"
## CoGAPS                  "CoGAPS"                 "3.14.0"
## cogena                  "cogena"                 "1.28.0"
## colorspace              "colorspace"             "2.0-2"
```

```
## commonmark            "commonmark"            "1.7"
## ComplexHeatmap        "ComplexHeatmap"        "2.10.0"
## ComplexUpset          "ComplexUpset"          "1.3.1"
## conquer               "conquer"               "1.2.1"
## corrplot              "corrplot"              "0.92"
## cowplot               "cowplot"               "1.1.1"
## cpp11                 "cpp11"                 "0.4.2"
## crayon                "crayon"                "1.4.2"
## credentials           "credentials"           "1.3.1"
## crosstalk             "crosstalk"             "1.2.0"
## curl                  "curl"                  "4.3.2"
## data.table            "data.table"            "1.14.2"
## DBI                   "DBI"                   "1.1.1"
## dbplyr                "dbplyr"                "1.3.0"
## DelayedArray          "DelayedArray"          "0.20.0"
## dendextend            "dendextend"            "1.15.2"
## DEoptimR              "DEoptimR"              "1.0-9"
## desc                  "desc"                  "1.4.0"
## DESeq2                "DESeq2"                "1.34.0"
## devtools              "devtools"              "2.4.3"
## diffobj               "diffobj"               "0.3.5"
## digest                "digest"                "0.6.29"
## DO.db                 "DO.db"                 "2.9"
## docopt                "docopt"                "0.7.1"
## doParallel            "doParallel"            "1.0.16"
## DOSE                  "DOSE"                  "3.18.3"
## downloader            "downloader"            "0.4"
## dplyr                 "dplyr"                 "1.0.7"
## DT                    "DT"                    "0.20"
## dtplyr                "dtplyr"                "1.2.0"
## e1071                 "e1071"                 "1.7-9"
## ellipse               "ellipse"               "0.4.2"
## ellipsis              "ellipsis"              "0.3.2"
## enrichplot            "enrichplot"            "1.12.3"
## EnsDb.Hsapiens.v75    "EnsDb.Hsapiens.v75"    "2.99.0"
## ensembldb             "ensembldb"             "2.18.2"
## evaluate              "evaluate"              "0.14"
## ExperimentHub         "ExperimentHub"         "2.0.0"
## factoextra            "factoextra"            "1.0.7"
## FactoMineR            "FactoMineR"            "2.4"
## fansi                 "fansi"                 "0.5.0"
## farver                "farver"                "2.1.0"
## fastcluster           "fastcluster"           "1.2.3"
## fastmap               "fastmap"               "1.1.0"
## fastmatch             "fastmatch"             "1.1-3"
## fgsea                 "fgsea"                 "1.18.0"
## filelock              "filelock"              "1.0.2"
## flashClust            "flashClust"            "1.01-2"
## fontawesome           "fontawesome"           "0.2.2"
## forcats               "forcats"               "0.5.1"
## foreach               "foreach"               "1.5.1"
## formatR               "formatR"               "1.11"
```

```
## fs                     "fs"                     "1.5.2"
## futile.logger          "futile.logger"          "1.4.3"
## futile.options         "futile.options"         "1.0.1"
## future                 "future"                 "1.23.0"
## future.apply           "future.apply"           "1.8.1"
## gargle                 "gargle"                 "1.2.0"
## genefilter             "genefilter"             "1.76.0"
## geneplotter            "geneplotter"            "1.72.0"
## generics               "generics"               "0.1.1"
## GenomeInfoDb           "GenomeInfoDb"           "1.30.0"
## GenomeInfoDbData       "GenomeInfoDbData"       "1.2.7"
## GenomicAlignments      "GenomicAlignments"      "1.30.0"
## GenomicFeatures        "GenomicFeatures"        "1.46.1"
## GenomicRanges          "GenomicRanges"          "1.46.1"
## GEOquery               "GEOquery"               "2.62.1"
## gert                   "gert"                   "1.4.3"
## GetoptLong             "GetoptLong"             "1.0.5"
## ggalluvial             "ggalluvial"             "0.12.3"
## ggforce                "ggforce"                "0.3.3"
## ggfun                  "ggfun"                  "0.0.4"
## ggplot2                "ggplot2"                "3.3.5"
## ggplotify              "ggplotify"              "0.1.0"
## ggpubr                 "ggpubr"                 "0.4.0"
## ggraph                 "ggraph"                 "2.0.5"
## ggrepel                "ggrepel"                "0.9.1"
## ggsci                  "ggsci"                  "2.9"
## ggsignif               "ggsignif"               "0.6.3"
## ggtree                 "ggtree"                 "3.0.4"
## gh                     "gh"                     "1.3.0"
## gitcreds               "gitcreds"               "0.1.1"
## glmnet                 "glmnet"                 "4.1-3"
## GlobalOptions          "GlobalOptions"          "0.1.2"
## globals                "globals"                "0.14.0"
## glue                   "glue"                   "1.5.1"
## GO.db                  "GO.db"                  "3.14.0"
## googledrive            "googledrive"            "2.0.0"
## googlesheets4          "googlesheets4"          "1.0.0"
## GOSemSim               "GOSemSim"               "2.18.1"
## gower                  "gower"                  "0.2.2"
## gplots                 "gplots"                 "3.1.1"
## gprofiler2             "gprofiler2"             "0.2.1"
## graph                  "graph"                  "1.72.0"
## graphlayouts           "graphlayouts"           "0.7.2"
## gridBase               "gridBase"               "0.4-7"
## gridExtra              "gridExtra"              "2.3"
## gridGraphics           "gridGraphics"           "0.5-1"
## GSEABase               "GSEABase"               "1.54.0"
## gtable                 "gtable"                 "0.3.0"
## gtools                 "gtools"                 "3.9.2"
## haven                  "haven"                  "2.4.3"
## HDF5Array              "HDF5Array"              "1.20.0"
## highr                  "highr"                  "0.9"
```

```
## hms                    "hms"                    "1.1.1"
## htmltools              "htmltools"              "0.5.2"
## htmlwidgets            "htmlwidgets"            "1.5.4"
## httpuv                 "httpuv"                 "1.6.3"
## httr                   "httr"                   "1.4.2"
## ids                    "ids"                    "1.0.1"
## igraph                 "igraph"                 "1.2.9"
## ini                    "ini"                    "0.3.1"
## interactiveDisplayBase "interactiveDisplayBase" "1.30.0"
## ipred                  "ipred"                  "0.9-12"
## IRanges                "IRanges"                "2.28.0"
## isoband                "isoband"                "0.2.5"
## iterators              "iterators"              "1.0.13"
## jquerylib              "jquerylib"              "0.1.4"
## jsonlite               "jsonlite"               "1.7.2"
## KEGGREST               "KEGGREST"               "1.34.0"
## knitr                  "knitr"                  "1.36"
## kohonen                "kohonen"                "3.0.10"
## labeling               "labeling"               "0.4.2"
## lambda.r               "lambda.r"               "1.2.4"
## later                  "later"                  "1.3.0"
## lava                   "lava"                   "1.6.10"
## lazyeval               "lazyeval"               "0.2.2"
## leaps                  "leaps"                  "3.1"
## lifecycle              "lifecycle"              "1.0.1"
## limma                  "limma"                  "3.50.0"
## listenv                "listenv"                "0.8.0"
## littler                "littler"                "0.3.14"
## lme4                   "lme4"                   "1.1-27.1"
## locfit                 "locfit"                 "1.5-9.4"
## lubridate              "lubridate"              "1.8.0"
## magrittr               "magrittr"               "2.0.1"
## maptools               "maptools"               "1.1-2"
## markdown               "markdown"               "1.1"
## MatrixGenerics         "MatrixGenerics"         "1.6.0"
## MatrixModels           "MatrixModels"           "0.5-0"
## matrixStats            "matrixStats"            "0.61.0"
## mclust                 "mclust"                 "5.4.8"
## memoise                "memoise"                "2.0.1"
## mime                   "mime"                   "0.12"
## minqa                  "minqa"                  "1.2.4"
## ModelMetrics           "ModelMetrics"           "1.2.2.2"
## modelr                 "modelr"                 "0.1.8"
## munsell                "munsell"                "0.5.0"
## mvtnorm                "mvtnorm"                "1.1-3"
## nloptr                 "nloptr"                 "1.2.2.3"
## NMF                    "NMF"                    "0.23.0"
## numDeriv               "numDeriv"               "2016.8-1.1"
## openssl                "openssl"                "1.4.5"
## org.Hs.eg.db           "org.Hs.eg.db"           "3.14.0"
## parallelly             "parallelly"             "1.29.0"
## pasilla                "pasilla"                "1.22.0"
```

```
## patchwork            "patchwork"            "1.1.1"
## pbkrtest             "pbkrtest"             "0.5.1"
## pcaPP                "pcaPP"                "1.9-74"
## pheatmap             "pheatmap"             "1.0.12"
## pillar               "pillar"               "1.6.4"
## pkgbuild             "pkgbuild"             "1.2.0"
## pkgconfig            "pkgconfig"            "2.0.3"
## pkgload              "pkgload"              "1.2.3"
## pkgmaker             "pkgmaker"             "0.32.2"
## PLIER                "PLIER"                "0.99.0"
## plogr                "plogr"                "0.2.0"
## plotly               "plotly"               "4.10.0"
## plyr                 "plyr"                 "1.8.6"
## png                  "png"                  "0.1-7"
## polyclip             "polyclip"             "1.10-0"
## polynom              "polynom"              "1.4-0"
## praise               "praise"               "1.0.0"
## preprocessCore       "preprocessCore"       "1.56.0"
## prettyunits          "prettyunits"          "1.1.1"
## pROC                 "pROC"                 "1.18.0"
## processx             "processx"             "3.5.2"
## prodlim              "prodlim"              "2019.11.13"
## progress             "progress"             "1.2.2"
## progressr            "progressr"            "0.9.0"
## projectR             "projectR"             "1.10.0"
## ProliferativeIndex   "ProliferativeIndex"   "1.0.1"
## promises             "promises"             "1.2.0.1"
## ProtGenerics         "ProtGenerics"         "1.26.0"
## proxy                "proxy"                "0.4-26"
## ps                   "ps"                   "1.6.0"
## purrr                "purrr"                "0.3.4"
## quantreg             "quantreg"             "5.86"
## qvalue               "qvalue"               "2.24.0"
## R.methodsS3          "R.methodsS3"          "1.8.1"
## R.oo                 "R.oo"                 "1.24.0"
## R.utils              "R.utils"              "2.11.0"
## R6                   "R6"                   "2.5.1"
## rapiclient           "rapiclient"           "0.1.3"
## rappdirs             "rappdirs"             "0.3.3"
## rcmdcheck            "rcmdcheck"            "1.4.0"
## RColorBrewer         "RColorBrewer"         "1.1-2"
## Rcpp                 "Rcpp"                 "1.0.7"
## RcppArmadillo        "RcppArmadillo"        "0.10.7.3.0"
## RcppEigen            "RcppEigen"            "0.3.3.9.1"
## RCurl                "RCurl"                "1.98-1.5"
## reactome.db          "reactome.db"          "1.76.0"
## readr                "readr"                "2.1.1"
## readxl               "readxl"               "1.3.1"
## recipes              "recipes"              "0.1.17"
## recount3             "recount3"             "1.4.0"
## registry             "registry"             "0.5-1"
## rematch              "rematch"              "1.0.1"
```

```
## rematch2               "rematch2"               "2.1.2"
## remotes                "remotes"                "2.4.1"
## reprex                 "reprex"                 "2.0.1"
## reshape2               "reshape2"               "1.4.4"
## restfulr               "restfulr"               "0.0.13"
## rhdf5                  "rhdf5"                  "2.38.0"
## rhdf5filters           "rhdf5filters"           "1.6.0"
## Rhdf5lib               "Rhdf5lib"               "1.16.0"
## Rhtslib                "Rhtslib"                "1.26.0"
## rjson                  "rjson"                  "0.2.20"
## rlang                  "rlang"                  "0.4.12"
## rmarkdown              "rmarkdown"              "2.11"
## rngtools               "rngtools"               "1.5.2"
## robustbase             "robustbase"             "0.93-9"
## ROCR                   "ROCR"                   "1.0-11"
## roxygen2               "roxygen2"               "7.1.2"
## rprojroot              "rprojroot"              "2.0.2"
## rrcov                  "rrcov"                  "1.6-0"
## Rsamtools              "Rsamtools"              "2.10.0"
## RSQLite                "RSQLite"                "2.2.9"
## rstatix                "rstatix"                "0.7.0"
## rstudioapi             "rstudioapi"             "0.13"
## rsvd                   "rsvd"                   "1.0.5"
## rtracklayer            "rtracklayer"            "1.54.0"
## rversions              "rversions"              "2.1.1"
## rvest                  "rvest"                  "1.0.2"
## S4Vectors              "S4Vectors"              "0.32.3"
## sass                   "sass"                   "0.4.0"
## scales                 "scales"                 "1.1.1"
## scatterpie             "scatterpie"             "0.1.7"
## scatterplot3d          "scatterplot3d"          "0.3-41"
## selectr                "selectr"                "0.4-2"
## sessioninfo            "sessioninfo"            "1.2.1"
## shadowtext             "shadowtext"             "0.0.9"
## shape                  "shape"                  "1.4.6"
## shiny                  "shiny"                  "1.7.1"
## signatureSearch        "signatureSearch"        "1.9.2"
## signatureSearchData    "signatureSearchData"    "1.8.2"
## SingleCellExperiment   "SingleCellExperiment"   "1.16.0"
## snow                   "snow"                   "0.4-4"
## sourcetools            "sourcetools"            "0.1.7"
## sp                     "sp"                     "1.4-6"
## SparseM                "SparseM"                "1.81"
## SQUAREM                "SQUAREM"                "2021.1"
## stringi                "stringi"                "1.7.6"
## stringr                "stringr"                "1.4.0"
## SummarizedExperiment   "SummarizedExperiment"   "1.24.0"
## sys                    "sys"                    "3.4"
## testthat               "testthat"               "3.1.0"
## TFEA.ChIP              "TFEA.ChIP"              "1.14.0"
## tibble                 "tibble"                 "3.1.6"
## tidygraph              "tidygraph"              "1.2.0"
```

```
## tidyr                "tidyr"               "1.1.4"
## tidyselect           "tidyselect"          "1.1.1"
## tidytree             "tidytree"            "0.3.6"
## tidyverse            "tidyverse"           "1.3.1"
## timeDate             "timeDate"            "3043.102"
## tinytex              "tinytex"             "0.35"
## topGO                "topGO"               "2.46.0"
## treeio               "treeio"              "1.16.2"
## tweenr               "tweenr"              "1.0.2"
## tzdb                 "tzdb"                "0.2.0"
## usethis              "usethis"             "2.1.3"
## utf8                 "utf8"                "1.2.2"
## uuid                 "uuid"                "1.0-3"
## vctrs                "vctrs"               "0.3.8"
## VennDiagram          "VennDiagram"         "1.7.1"
## viridis              "viridis"             "0.6.2"
## viridisLite          "viridisLite"         "0.4.0"
## visNetwork           "visNetwork"          "2.1.0"
## vroom                "vroom"               "1.5.7"
## waldo                "waldo"               "0.3.1"
## whisker              "whisker"             "0.4"
## withr                "withr"               "2.4.3"
## xfun                 "xfun"                "0.28"
## XML                  "XML"                 "3.99-0.8"
## xml2                 "xml2"                "1.3.2"
## xopen                "xopen"               "1.0.0"
## xtable               "xtable"              "1.8-4"
## XVector              "XVector"             "0.34.0"
## yaml                 "yaml"                "2.2.1"
## yulab.utils          "yulab.utils"         "0.0.4"
## zip                  "zip"                 "2.2.0"
## zlibbioc             "zlibbioc"            "1.40.0"
## base                 "base"                "4.1.2"
## boot                 "boot"                "1.3-28"
## class                "class"               "7.3-19"
## cluster              "cluster"             "2.1.2"
## codetools            "codetools"           "0.2-18"
## compiler             "compiler"            "4.1.2"
## datasets             "datasets"            "4.1.2"
## foreign              "foreign"             "0.8-81"
## graphics             "graphics"            "4.1.2"
## grDevices            "grDevices"           "4.1.2"
## grid                 "grid"                "4.1.2"
## KernSmooth           "KernSmooth"          "2.23-20"
## lattice              "lattice"             "0.20-45"
## MASS                 "MASS"                "7.3-54"
## Matrix               "Matrix"              "1.3-4"
## methods              "methods"             "4.1.2"
## mgcv                 "mgcv"                "1.8-38"
## nlme                 "nlme"                "3.1-153"
## nnet                 "nnet"                "7.3-16"
## parallel             "parallel"            "4.1.2"
```

```
## rpart                  "rpart"                 "4.1-15"
## spatial                "spatial"               "7.3-14"
## splines                "splines"               "4.1.2"
## stats                  "stats"                 "4.1.2"
## stats4                 "stats4"                "4.1.2"
## survival               "survival"              "3.2-13"
## tcltk                  "tcltk"                 "4.1.2"
## tools                  "tools"                 "4.1.2"
## utils                  "utils"                 "4.1.2"
```

```
R version 4.0.5 (2021-03-31)
Platform: x86_64-conda-linux-gnu (64-bit)
Running under: Red Hat Enterprise Linux

Matrix products: default
BLAS/LAPACK: /data/user/jfisher7/.conda/envs/SR_TAU_CELL/lib/
libopenblasp-r0.3.18.so

locale:
[1] C

attached base packages:
[1] stats     graphics  grDevices utils     datasets  methods   base

loaded via a namespace (and not attached):
[1] compiler_4.0.5
[1] "Listing packages"
                       Package                  Version
AnnotationDbi          "AnnotationDbi"          "1.52.0"
AnnotationHub          "AnnotationHub"          "2.22.1"
BH                     "BH"                     "1.78.0-0"
Biobase                "Biobase"                "2.50.0"
BiocFileCache          "BiocFileCache"          "1.14.0"
BiocGenerics           "BiocGenerics"           "0.36.1"
BiocManager            "BiocManager"            "1.30.16"
BiocParallel           "BiocParallel"           "1.24.1"
BiocVersion            "BiocVersion"            "3.12.0"
DBI                    "DBI"                    "1.1.2"
DO.db                  "DO.db"                  "2.9"
DOSE                   "DOSE"                   "3.16.0"
DT                     "DT"                     "0.20"
DelayedArray           "DelayedArray"           "0.16.3"
ExperimentHub          "ExperimentHub"          "1.16.1"
GO.db                  "GO.db"                  "3.12.1"
GOSemSim               "GOSemSim"               "2.16.1"
GSEABase               "GSEABase"               "1.52.1"
GenomeInfoDb           "GenomeInfoDb"           "1.26.7"
GenomeInfoDbData       "GenomeInfoDbData"       "1.2.4"
GenomicRanges          "GenomicRanges"          "1.42.0"
HDF5Array              "HDF5Array"              "1.18.1"
IRanges                "IRanges"                "2.24.1"
MASS                   "MASS"                   "7.3-55"
Matrix                 "Matrix"                 "1.4-0"
MatrixGenerics         "MatrixGenerics"         "1.2.1"
R.methodsS3            "R.methodsS3"            "1.8.1"
R.oo                   "R.oo"                   "1.24.0"
R.utils                "R.utils"                "2.11.0"
R6                     "R6"                     "2.5.1"
RColorBrewer           "RColorBrewer"           "1.1-2"
RCurl                  "RCurl"                  "1.98-1.5"
```

```
RSQLite                "RSQLite"                "2.2.9"
Rcpp                   "Rcpp"                   "1.0.8"
RcppArmadillo          "RcppArmadillo"          "0.10.8.1.0"
RcppEigen              "RcppEigen"              "0.3.3.9.1"
Rhdf5lib               "Rhdf5lib"               "1.12.1"
S4Vectors              "S4Vectors"              "0.28.1"
SummarizedExperiment   "SummarizedExperiment"   "1.20.0"
XML                    "XML"                    "3.99-0.8"
XVector                "XVector"                "0.30.0"
affy                   "affy"                   "1.68.0"
affyio                 "affyio"                 "1.60.0"
annotate               "annotate"               "1.68.0"
askpass                "askpass"                "1.1"
assertthat             "assertthat"             "0.2.1"
backports              "backports"              "1.4.1"
base                   "base"                   "4.0.5"
base64enc              "base64enc"              "0.1-3"
bit                    "bit"                    "4.0.4"
bit64                  "bit64"                  "4.0.5"
bitops                 "bitops"                 "1.0-7"
blob                   "blob"                   "1.2.2"
brio                   "brio"                   "1.1.3"
broom                  "broom"                  "0.7.12"
bslib                  "bslib"                  "0.3.1"
cachem                 "cachem"                 "1.0.6"
callr                  "callr"                  "3.7.0"
cellranger             "cellranger"             "1.1.0"
cli                    "cli"                    "3.1.1"
clipr                  "clipr"                  "0.7.1"
clusterProfiler        "clusterProfiler"        "3.18.1"
colorspace             "colorspace"             "2.0-2"
commonmark             "commonmark"             "1.7"
compiler               "compiler"               "4.0.5"
cowplot                "cowplot"                "1.1.1"
cpp11                  "cpp11"                  "0.4.2"
crayon                 "crayon"                 "1.4.2"
crosstalk              "crosstalk"              "1.2.0"
curl                   "curl"                   "4.3.2"
data.table             "data.table"             "1.14.2"
datasets               "datasets"               "4.0.5"
dbplyr                 "dbplyr"                 "2.1.1"
desc                   "desc"                   "1.4.0"
diffobj                "diffobj"                "0.3.5"
digest                 "digest"                 "0.6.29"
downloader             "downloader"             "0.4"
dplyr                  "dplyr"                  "1.0.7"
dtplyr                 "dtplyr"                 "1.2.1"
ellipsis               "ellipsis"               "0.3.2"
enrichplot             "enrichplot"             "1.10.2"
evaluate               "evaluate"               "0.14"
```

```
fansi                  "fansi"                  "1.0.2"
farver                 "farver"                 "2.1.0"
fastmap                "fastmap"                "1.1.0"
fastmatch              "fastmatch"              "1.1-3"
fgsea                  "fgsea"                  "1.16.0"
fontawesome            "fontawesome"            "0.2.2"
forcats                "forcats"                "0.5.1"
formatR                "formatR"                "1.11"
fs                     "fs"                     "1.5.2"
futile.logger          "futile.logger"          "1.4.3"
futile.options         "futile.options"         "1.0.1"
gargle                 "gargle"                 "1.2.0"
generics               "generics"               "0.1.2"
ggforce                "ggforce"                "0.3.3"
ggfun                  "ggfun"                  "0.0.5"
ggplot2                "ggplot2"                "3.3.5"
ggraph                 "ggraph"                 "2.0.5"
ggrepel                "ggrepel"                "0.9.1"
glue                   "glue"                   "1.6.1"
googledrive            "googledrive"            "2.0.0"
googlesheets4          "googlesheets4"          "1.0.0"
grDevices              "grDevices"              "4.0.5"
graph                  "graph"                  "1.68.0"
graphics               "graphics"               "4.0.5"
graphlayouts           "graphlayouts"           "0.8.0"
grid                   "grid"                   "4.0.5"
gridExtra              "gridExtra"              "2.3"
gtable                 "gtable"                 "0.3.0"
haven                  "haven"                  "2.4.3"
highr                  "highr"                  "0.9"
hms                    "hms"                    "1.1.1"
htmltools              "htmltools"              "0.5.2"
htmlwidgets            "htmlwidgets"            "1.5.4"
httpuv                 "httpuv"                 "1.6.5"
httr                   "httr"                   "1.4.2"
ids                    "ids"                    "1.0.1"
igraph                 "igraph"                 "1.2.11"
interactiveDisplayBase "interactiveDisplayBase" "1.28.0"
isoband                "isoband"                "0.2.5"
jquerylib              "jquerylib"              "0.1.4"
jsonlite               "jsonlite"               "1.7.3"
knitr                  "knitr"                  "1.37"
labeling               "labeling"               "0.4.2"
lambda.r               "lambda.r"               "1.2.4"
later                  "later"                  "1.3.0"
lattice                "lattice"                "0.20-45"
lazyeval               "lazyeval"               "0.2.2"
lifecycle              "lifecycle"              "1.0.1"
limma                  "limma"                  "3.46.0"
lubridate              "lubridate"              "1.8.0"
```

```
magrittr            "magrittr"            "2.0.2"
matrixStats         "matrixStats"         "0.61.0"
memoise             "memoise"             "2.0.1"
methods             "methods"             "4.0.5"
mgcv                "mgcv"                "1.8-38"
mime                "mime"                "0.12"
modelr              "modelr"              "0.1.8"
munsell             "munsell"             "0.5.0"
nlme                "nlme"                "3.1-155"
openssl             "openssl"             "1.4.6"
parallel            "parallel"            "4.0.5"
pillar              "pillar"              "1.7.0"
pkgconfig           "pkgconfig"           "2.0.3"
pkgload             "pkgload"             "1.2.4"
plogr               "plogr"               "0.2.0"
plyr                "plyr"                "1.8.6"
polyclip            "polyclip"            "1.10-0"
praise              "praise"              "1.0.0"
preprocessCore      "preprocessCore"      "1.52.1"
prettyunits         "prettyunits"         "1.1.1"
processx            "processx"            "3.5.2"
progress            "progress"            "1.2.2"
promises            "promises"            "1.2.0.1"
ps                  "ps"                  "1.6.0"
purrr               "purrr"               "0.3.4"
qvalue              "qvalue"              "2.22.0"
rappdirs            "rappdirs"            "0.3.3"
reactome.db         "reactome.db"         "1.74.0"
readr               "readr"               "2.1.2"
readxl              "readxl"              "1.3.1"
rematch             "rematch"             "1.0.1"
rematch2            "rematch2"            "2.1.2"
reprex              "reprex"              "2.0.1"
reshape2            "reshape2"            "1.4.4"
rhdf5               "rhdf5"               "2.34.0"
rhdf5filters        "rhdf5filters"        "1.2.1"
rlang               "rlang"               "1.0.0"
rmarkdown           "rmarkdown"           "2.11"
rprojroot           "rprojroot"           "2.0.2"
rstudioapi          "rstudioapi"          "0.13"
rvcheck             "rvcheck"             "0.2.1"
rvest               "rvest"               "1.0.2"
sass                "sass"                "0.4.0"
scales              "scales"              "1.1.1"
scatterpie          "scatterpie"          "0.1.7"
selectr             "selectr"             "0.4-2"
shadowtext          "shadowtext"          "0.1.1"
shiny               "shiny"               "1.7.1"
signatureSearch     "signatureSearch"     "1.4.6"
signatureSearchData "signatureSearchData" "1.4.0"
```

| | | |
|---|---|---|
| snow | "snow" | "0.4-4" |
| sourcetools | "sourcetools" | "0.1.7" |
| splines | "splines" | "4.0.5" |
| stats | "stats" | "4.0.5" |
| stats4 | "stats4" | "4.0.5" |
| stringi | "stringi" | "1.7.6" |
| stringr | "stringr" | "1.4.0" |
| sys | "sys" | "3.4" |
| tcltk | "tcltk" | "4.0.5" |
| testthat | "testthat" | "3.1.2" |
| tibble | "tibble" | "3.1.6" |
| tidygraph | "tidygraph" | "1.2.0" |
| tidyr | "tidyr" | "1.2.0" |
| tidyselect | "tidyselect" | "1.1.1" |
| tidyverse | "tidyverse" | "1.3.1" |
| tinytex | "tinytex" | "0.36" |
| tools | "tools" | "4.0.5" |
| tweenr | "tweenr" | "1.0.2" |
| tzdb | "tzdb" | "0.2.0" |
| utf8 | "utf8" | "1.2.2" |
| utils | "utils" | "4.0.5" |
| uuid | "uuid" | "1.0-3" |
| vctrs | "vctrs" | "0.3.8" |
| viridis | "viridis" | "0.6.2" |
| viridisLite | "viridisLite" | "0.4.0" |
| visNetwork | "visNetwork" | "2.1.0" |
| vroom | "vroom" | "1.5.7" |
| waldo | "waldo" | "0.3.1" |
| withr | "withr" | "2.4.3" |
| xfun | "xfun" | "0.29" |
| xml2 | "xml2" | "1.3.3" |
| xtable | "xtable" | "1.8-4" |
| yaml | "yaml" | "2.2.2" |
| yulab.utils | "yulab.utils" | "0.0.4" |
| zlibbioc | "zlibbioc" | "1.36.0" |

### 221221_rstudio_cancer_dr_info

Jennifer Fisher

2022-12-21

```
GitHub Repo:
Transfer_Leanring_R03

Docker:
rstudio_cancer_dr

Directory of operations:
/home/rstudio (Docker)
```

```
sessionInfo()
```

```
## R version 4.1.3 (2022-03-10)
## Platform: x86_64-pc-linux-gnu (64-bit)
## Running under: Ubuntu 20.04.4 LTS
##
## Matrix products: default
## BLAS/LAPACK: /usr/lib/x86_64-linux-gnu/openblas-pthread/libopenblasp-r0.3.8.so
##
## locale:
##  [1] LC_CTYPE=en_US.UTF-8       LC_NUMERIC=C
##  [3] LC_TIME=en_US.UTF-8        LC_COLLATE=en_US.UTF-8
##  [5] LC_MONETARY=en_US.UTF-8    LC_MESSAGES=en_US.UTF-8
##  [7] LC_PAPER=en_US.UTF-8       LC_NAME=C
##  [9] LC_ADDRESS=C               LC_TELEPHONE=C
## [11] LC_MEASUREMENT=en_US.UTF-8 LC_IDENTIFICATION=C
##
## attached base packages:
## [1] stats     graphics  grDevices utils     datasets  methods   base
##
## loaded via a namespace (and not attached):
##  [1] digest_0.6.29   R6_2.5.1        jsonlite_1.8.0  magrittr_2.0.3
##  [5] evaluate_0.16   stringi_1.7.8   cachem_1.0.6    rlang_1.0.6
##  [9] cli_3.4.1       rstudioapi_0.13 jquerylib_0.1.4 bslib_0.4.0
## [13] rmarkdown_2.16  tools_4.1.3     stringr_1.4.1   xfun_0.33
## [17] yaml_2.3.5      fastmap_1.1.0   compiler_4.1.3  htmltools_0.5.3
## [21] knitr_1.40      sass_0.4.2
```

```
# Listing packages
installed.packages()[,c(1,3)]
```

```
##                         Package                   Version
## abind                   "abind"                   "1.4-5"
## affy                    "affy"                    "1.72.0"
## affyio                  "affyio"                  "1.64.0"
## amap                    "amap"                    "0.8-18"
## annotate                "annotate"                "1.72.0"
## AnnotationDbi           "AnnotationDbi"           "1.56.2"
## AnnotationFilter        "AnnotationFilter"        "1.18.0"
## AnnotationForge         "AnnotationForge"         "1.36.0"
## AnnotationHub           "AnnotationHub"           "3.2.2"
## AnVIL                   "AnVIL"                   "1.6.6"
## apcluster               "apcluster"               "1.4.10"
## ape                     "ape"                     "5.6-2"
## apeglm                  "apeglm"                  "1.16.0"
## aplot                   "aplot"                   "0.1.7"
## ashr                    "ashr"                    "2.2-55"
## askpass                 "askpass"                 "1.1"
## assertthat              "assertthat"              "0.2.1"
## backports               "backports"               "1.4.1"
## base64                  "base64"                  "2.0.1"
## base64enc               "base64enc"               "0.1-3"
## base64url               "base64url"               "1.4"
## bayestestR              "bayestestR"              "0.13.0"
## bbmle                   "bbmle"                   "1.0.25"
## bdsmatrix               "bdsmatrix"               "1.3-6"
## beanplot                "beanplot"                "1.3.1"
## BgeeDB                  "BgeeDB"                  "2.20.1"
## BH                      "BH"                      "1.78.0-0"
## Biobase                 "Biobase"                 "2.54.0"
## BiocFileCache           "BiocFileCache"           "2.2.1"
## BiocGenerics            "BiocGenerics"            "0.40.0"
## BiocIO                  "BiocIO"                  "1.4.0"
## BiocManager             "BiocManager"             "1.30.16"
## BiocParallel            "BiocParallel"            "1.28.3"
## BiocVersion             "BiocVersion"             "3.14.0"
## biomaRt                 "biomaRt"                 "2.50.3"
## Biostrings              "Biostrings"              "2.62.0"
## bit                     "bit"                     "4.0.4"
## bit64                   "bit64"                   "4.0.5"
## bitops                  "bitops"                  "1.0-7"
## biwt                    "biwt"                    "1.0.1"
## blob                    "blob"                    "1.2.3"
## brew                    "brew"                    "1.0-7"
## brio                    "brio"                    "1.1.3"
## broom                   "broom"                   "1.0.1"
## broom.mixed             "broom.mixed"             "0.2.9.4"
## bslib                   "bslib"                   "0.4.0"
## bumphunter              "bumphunter"              "1.36.0"
## cachem                  "cachem"                  "1.0.6"
## callr                   "callr"                   "3.7.0"
## car                     "car"                     "3.1-0"
## carData                 "carData"                 "3.0-5"
```

```
## caret                "caret"                "6.0-93"
## Category             "Category"             "2.60.0"
## caTools              "caTools"              "1.18.2"
## cellranger           "cellranger"           "1.1.0"
## ChemmineR            "ChemmineR"            "3.46.0"
## chron                "chron"                "2.3-57"
## circlize             "circlize"             "0.4.15"
## classInt             "classInt"             "0.4-7"
## cli                  "cli"                  "3.4.1"
## clipr                "clipr"                "0.8.0"
## clue                 "clue"                 "0.3-61"
## clusterProfiler      "clusterProfiler"      "4.2.2"
## coda                 "coda"                 "0.19-4"
## CoGAPS               "CoGAPS"               "3.14.0"
## cogena               "cogena"               "1.28.0"
## colorspace           "colorspace"           "2.0-3"
## combinat             "combinat"             "0.0-8"
## commonmark           "commonmark"           "1.8.0"
## ComplexHeatmap       "ComplexHeatmap"       "2.10.0"
## ComplexUpset         "ComplexUpset"         "1.3.3"
## conflicted           "conflicted"           "1.1.0"
## corrplot             "corrplot"             "0.92"
## cowplot              "cowplot"              "1.1.1"
## cpp11                "cpp11"                "0.4.2"
## crayon               "crayon"               "1.5.2"
## credentials          "credentials"          "1.3.2"
## crosstalk            "crosstalk"            "1.2.0"
## curl                 "curl"                 "4.3.2"
## customCMPdb          "customCMPdb"          "1.4.0"
## data.table           "data.table"           "1.14.2"
## datawizard           "datawizard"           "0.6.1"
## DBI                  "DBI"                  "1.1.3"
## dbplyr               "dbplyr"               "2.2.1"
## DelayedArray         "DelayedArray"         "0.20.0"
## DelayedMatrixStats   "DelayedMatrixStats"   "1.16.0"
## dendextend           "dendextend"           "1.16.0"
## DEoptimR             "DEoptimR"             "1.0-11"
## desc                 "desc"                 "1.4.1"
## DESeq2               "DESeq2"               "1.34.0"
## devtools             "devtools"             "2.4.4"
## dials                "dials"                "1.0.0"
## DiceDesign           "DiceDesign"           "1.9"
## diffobj              "diffobj"              "0.3.5"
## digest               "digest"               "0.6.29"
## discrim              "discrim"              "1.0.0"
## DO.db                "DO.db"                "2.9"
## docopt               "docopt"               "0.7.1"
## doParallel           "doParallel"           "1.0.17"
## doRNG                "doRNG"                "1.8.2"
## DOSE                 "DOSE"                 "3.20.1"
## dotwhisker           "dotwhisker"           "0.7.4"
## downlit              "downlit"              "0.4.2"
```

```
## downloader            "downloader"            "0.4"
## dplyr                 "dplyr"                 "1.0.10"
## drugbankR             "drugbankR"             "1.5"
## DT                    "DT"                    "0.25"
## dtplyr                "dtplyr"                "1.2.2"
## e1071                 "e1071"                 "1.7-11"
## earth                 "earth"                 "5.3.1"
## edgeR                 "edgeR"                 "3.36.0"
## ellipse               "ellipse"               "0.4.3"
## ellipsis              "ellipsis"              "0.3.2"
## emdbook               "emdbook"               "1.3.12"
## emmeans               "emmeans"               "1.8.1-1"
## enrichplot            "enrichplot"            "1.14.2"
## EnsDb.Hsapiens.v75    "EnsDb.Hsapiens.v75"    "2.99.0"
## ensembldb             "ensembldb"             "2.18.4"
## estimability          "estimability"          "1.4.1"
## etrunct               "etrunct"               "0.1"
## evaluate              "evaluate"              "0.16"
## ExperimentHub         "ExperimentHub"         "2.2.1"
## factoextra            "factoextra"            "1.0.7"
## FactoMineR            "FactoMineR"            "2.6"
## fansi                 "fansi"                 "1.0.3"
## farver                "farver"                "2.1.1"
## fastcluster           "fastcluster"           "1.2.3"
## fastmap               "fastmap"               "1.1.0"
## fastmatch             "fastmatch"             "1.1-3"
## fgsea                 "fgsea"                 "1.20.0"
## filelock              "filelock"              "1.0.2"
## flashClust            "flashClust"            "1.01-2"
## fmcsR                 "fmcsR"                 "1.36.0"
## fontawesome           "fontawesome"           "0.3.0"
## forcats               "forcats"               "0.5.2"
## foreach               "foreach"               "1.5.2"
## formatR               "formatR"               "1.12"
## Formula               "Formula"               "1.2-4"
## fs                    "fs"                    "1.5.2"
## furrr                 "furrr"                 "0.3.1"
## futile.logger         "futile.logger"         "1.4.3"
## futile.options        "futile.options"        "1.0.1"
## future                "future"                "1.28.0"
## future.apply          "future.apply"          "1.9.1"
## gargle                "gargle"                "1.2.1"
## gbm                   "gbm"                   "2.1.8.1"
## genefilter            "genefilter"            "1.76.0"
## geneplotter           "geneplotter"           "1.72.0"
## generics              "generics"              "0.1.3"
## GenomeInfoDb          "GenomeInfoDb"          "1.30.1"
## GenomeInfoDbData      "GenomeInfoDbData"      "1.2.7"
## GenomicAlignments     "GenomicAlignments"     "1.30.0"
## GenomicFeatures       "GenomicFeatures"       "1.46.5"
## GenomicRanges         "GenomicRanges"         "1.46.1"
## GEOquery              "GEOquery"              "2.62.2"
```

```
## gert                   "gert"                   "1.5.0"
## GetoptLong             "GetoptLong"             "1.0.5"
## ggalluvial             "ggalluvial"             "0.12.3"
## ggdendro               "ggdendro"               "0.1.23"
## ggforce                "ggforce"                "0.3.4"
## ggfun                  "ggfun"                  "0.0.7"
## ggplot2                "ggplot2"                "3.3.6"
## ggplotify              "ggplotify"              "0.1.0"
## ggpubr                 "ggpubr"                 "0.4.0"
## ggraph                 "ggraph"                 "2.0.6"
## ggrepel                "ggrepel"                "0.9.1"
## ggsci                  "ggsci"                  "2.9"
## ggsignif               "ggsignif"               "0.6.3"
## ggstance               "ggstance"               "0.3.5"
## ggtree                 "ggtree"                 "3.2.1"
## gh                     "gh"                     "1.3.0"
## gitcreds               "gitcreds"               "0.1.1"
## glmnet                 "glmnet"                 "4.1-4"
## GlobalOptions          "GlobalOptions"          "0.1.2"
## globals                "globals"                "0.16.1"
## glue                   "glue"                   "1.6.2"
## GO.db                  "GO.db"                  "3.14.0"
## googledrive            "googledrive"            "2.0.0"
## googlesheets4          "googlesheets4"          "1.0.1"
## GOSemSim               "GOSemSim"               "2.20.0"
## GOstats                "GOstats"                "2.60.0"
## gower                  "gower"                  "1.0.0"
## GPfit                  "GPfit"                  "1.0-8"
## gplots                 "gplots"                 "3.1.3"
## gprofiler2             "gprofiler2"             "0.2.1"
## graph                  "graph"                  "1.72.0"
## graphlayouts           "graphlayouts"           "0.8.2"
## gridBase               "gridBase"               "0.4-7"
## gridExtra              "gridExtra"              "2.3"
## gridGraphics           "gridGraphics"           "0.5-1"
## GSEABase               "GSEABase"               "1.56.0"
## gsubfn                 "gsubfn"                 "0.7"
## gtable                 "gtable"                 "0.3.1"
## gtools                 "gtools"                 "3.9.3"
## hardhat                "hardhat"                "1.2.0"
## hash                   "hash"                   "2.2.6.2"
## haven                  "haven"                  "2.5.1"
## HDF5Array              "HDF5Array"              "1.22.1"
## here                   "here"                   "1.0.1"
## hexbin                 "hexbin"                 "1.28.2"
## highr                  "highr"                  "0.9"
## hms                    "hms"                    "1.1.2"
## htmltools              "htmltools"              "0.5.3"
## htmlwidgets            "htmlwidgets"            "1.5.4"
## httpuv                 "httpuv"                 "1.6.6"
## httr                   "httr"                   "1.4.4"
## ids                    "ids"                    "1.0.1"
```

```
## igraph                 "igraph"                 "1.3.5"
## illuminaio             "illuminaio"             "0.36.0"
## infer                  "infer"                  "1.0.3"
## ini                    "ini"                    "0.3.1"
## insight                "insight"                "0.18.4"
## interactiveDisplayBase "interactiveDisplayBase" "1.32.0"
## inum                   "inum"                   "1.0-4"
## invgamma               "invgamma"               "1.1"
## ipred                  "ipred"                  "0.9-13"
## IRanges                "IRanges"                "2.28.0"
## IRdisplay              "IRdisplay"              "1.1"
## IRkernel               "IRkernel"               "1.3"
## irlba                  "irlba"                  "2.3.5"
## isoband                "isoband"                "0.2.5"
## iterators              "iterators"              "1.0.14"
## jquerylib              "jquerylib"              "0.1.4"
## jsonlite               "jsonlite"               "1.8.0"
## KEGGREST               "KEGGREST"               "1.34.0"
## kernlab                "kernlab"                "0.9-31"
## klaR                   "klaR"                   "1.7-1"
## knitr                  "knitr"                  "1.40"
## kohonen                "kohonen"                "3.0.11"
## labeling               "labeling"               "0.4.2"
## labelled               "labelled"               "2.10.0"
## lambda.r               "lambda.r"               "1.2.4"
## later                  "later"                  "1.3.0"
## lava                   "lava"                   "1.6.10"
## lazyeval               "lazyeval"               "0.2.2"
## leaps                  "leaps"                  "3.1"
## lhs                    "lhs"                    "1.1.5"
## libcoin                "libcoin"                "1.0-9"
## LiblineaR              "LiblineaR"              "2.10-12"
## lifecycle              "lifecycle"              "1.0.2"
## limma                  "limma"                  "3.50.3"
## listenv                "listenv"                "0.8.0"
## littler                "littler"                "0.3.15"
## lme4                   "lme4"                   "1.1-30"
## locfit                 "locfit"                 "1.5-9.6"
## lubridate              "lubridate"              "1.8.0"
## magrittr               "magrittr"               "2.0.3"
## maptools               "maptools"               "1.1-4"
## margins                "margins"                "0.3.26"
## markdown               "markdown"               "1.4"
## mashr                  "mashr"                  "0.2.57"
## MASS                   "MASS"                   "7.3-58.1"
## Matrix                 "Matrix"                 "1.5-1"
## MatrixGenerics         "MatrixGenerics"         "1.6.0"
## MatrixModels           "MatrixModels"           "0.5-1"
## matrixStats            "matrixStats"            "0.62.0"
## mclust                 "mclust"                 "5.4.10"
## memoise                "memoise"                "2.0.1"
## mime                   "mime"                   "0.12"
```

```
## minfi                "minfi"                "1.40.0"
## miniUI               "miniUI"               "0.1.1.1"
## minqa                "minqa"                "1.2.4"
## mixsqp               "mixsqp"               "0.3-43"
## modeldata            "modeldata"            "1.0.1"
## ModelMetrics         "ModelMetrics"         "1.2.2.2"
## modelr               "modelr"               "0.1.9"
## multcompView         "multcompView"         "0.1-8"
## multtest             "multtest"             "2.50.0"
## munsell              "munsell"              "0.5.0"
## mvtnorm              "mvtnorm"              "1.1-3"
## naivebayes           "naivebayes"           "0.9.7"
## netZooR              "netZooR"              "1.1.15"
## nloptr               "nloptr"               "2.0.3"
## NLP                  "NLP"                  "0.2-1"
## NMF                  "NMF"                  "0.24.0"
## nnet                 "nnet"                 "7.3-17"
## nor1mix              "nor1mix"              "1.3-0"
## numDeriv             "numDeriv"             "2016.8-1.1"
## openssl              "openssl"              "2.0.3"
## org.Hs.eg.db         "org.Hs.eg.db"         "3.14.0"
## pamr                 "pamr"                 "1.56.1"
## pandaR               "pandaR"               "1.26.0"
## parallelly           "parallelly"           "1.32.1"
## parameters           "parameters"           "0.18.2"
## parsnip              "parsnip"              "1.0.1"
## partykit             "partykit"             "1.2-16"
## pasilla              "pasilla"              "1.22.0"
## patchwork            "patchwork"            "1.1.2"
## pbdZMQ               "pbdZMQ"               "0.3-7"
## pbkrtest             "pbkrtest"             "0.5.1"
## penalized            "penalized"            "0.9-52"
## permute              "permute"              "0.9-7"
## pheatmap             "pheatmap"             "1.0.12"
## pillar               "pillar"               "1.8.1"
## pkgbuild             "pkgbuild"             "1.3.1"
## pkgconfig            "pkgconfig"            "2.0.3"
## pkgdown              "pkgdown"              "2.0.6"
## pkgload              "pkgload"              "1.3.0"
## pkgmaker             "pkgmaker"             "0.32.2"
## PLIER                "PLIER"                "0.99.0"
## plogr                "plogr"                "0.2.0"
## plotly               "plotly"               "4.10.0"
## plotmo               "plotmo"               "3.6.2"
## plotrix              "plotrix"              "3.8-2"
## plyr                 "plyr"                 "1.8.7"
## png                  "png"                  "0.1-7"
## polyclip             "polyclip"             "1.10-0"
## polynom              "polynom"              "1.4-1"
## praise               "praise"               "1.0.0"
## prediction           "prediction"           "0.3.14"
## preprocessCore       "preprocessCore"       "1.56.0"
```

```
## prettyunits            "prettyunits"            "1.1.1"
## pROC                   "pROC"                   "1.18.0"
## processx               "processx"               "3.5.2"
## prodlim                "prodlim"                "2019.11.13"
## profvis                "profvis"                "0.3.7"
## progress               "progress"               "1.2.2"
## progressr              "progressr"              "0.11.0"
## projectR               "projectR"               "1.10.0"
## ProliferativeIndex     "ProliferativeIndex"     "1.0.1"
## promises               "promises"               "1.2.0.1"
## ProtGenerics           "ProtGenerics"           "1.26.0"
## proto                  "proto"                  "1.0.0"
## proxy                  "proxy"                  "0.4-27"
## ps                     "ps"                     "1.6.0"
## purrr                  "purrr"                  "0.3.4"
## quadprog               "quadprog"               "1.5-8"
## quantreg               "quantreg"               "5.94"
## quantro                "quantro"                "1.28.0"
## questionr              "questionr"              "0.7.7"
## qvalue                 "qvalue"                 "2.26.0"
## R.cache                "R.cache"                "0.16.0"
## R.methodsS3            "R.methodsS3"            "1.8.2"
## R.oo                   "R.oo"                   "1.25.0"
## R.utils                "R.utils"                "2.12.0"
## R6                     "R6"                     "2.5.1"
## ragg                   "ragg"                   "1.2.2"
## randomForest           "randomForest"           "4.7-1.1"
## ranger                 "ranger"                 "0.14.1"
## rapiclient             "rapiclient"             "0.1.3"
## rappdirs               "rappdirs"               "0.3.3"
## RBGL                   "RBGL"                   "1.70.0"
## rcmdcheck              "rcmdcheck"              "1.4.0"
## RColorBrewer           "RColorBrewer"           "1.1-3"
## Rcpp                   "Rcpp"                   "1.0.9"
## RcppArmadillo          "RcppArmadillo"          "0.11.2.4.0"
## RcppEigen              "RcppEigen"              "0.3.3.9.2"
## RcppGSL                "RcppGSL"                "0.3.11"
## RcppNumerical          "RcppNumerical"          "0.4-0"
## RcppTOML               "RcppTOML"               "0.1.7"
## RCurl                  "RCurl"                  "1.98-1.8"
## RCy3                   "RCy3"                   "2.14.2"
## reactome.db            "reactome.db"            "1.77.0"
## readr                  "readr"                  "2.1.2"
## readxl                 "readxl"                 "1.4.1"
## recipes                "recipes"                "1.0.1"
## recount3               "recount3"               "1.4.0"
## registry               "registry"               "0.5-1"
## rematch                "rematch"                "1.0.1"
## rematch2               "rematch2"               "2.1.2"
## remotes                "remotes"                "2.4.2"
## repr                   "repr"                   "1.1.4"
## reprex                 "reprex"                 "2.0.2"
```

```
## reshape               "reshape"               "0.8.9"
## reshape2              "reshape2"              "1.4.4"
## restfulr              "restfulr"              "0.0.15"
## reticulate            "reticulate"            "1.26"
## Rgraphviz             "Rgraphviz"             "2.38.0"
## rhdf5                 "rhdf5"                 "2.38.1"
## rhdf5filters          "rhdf5filters"          "1.6.0"
## Rhdf5lib              "Rhdf5lib"              "1.16.0"
## Rhtslib               "Rhtslib"               "1.26.0"
## rJava                 "rJava"                 "1.0-6"
## rjson                 "rjson"                 "0.2.21"
## RJSONIO               "RJSONIO"               "1.3-1.6"
## rlang                 "rlang"                 "1.0.6"
## rmarkdown             "rmarkdown"             "2.16"
## rmeta                 "rmeta"                 "3.0"
## rngtools              "rngtools"              "1.5.2"
## robustbase            "robustbase"            "0.95-0"
## ROCR                  "ROCR"                  "1.0-11"
## roxygen2              "roxygen2"              "7.2.1"
## rpart                 "rpart"                 "4.1.16"
## rprojroot             "rprojroot"             "2.0.3"
## rrvgo                 "rrvgo"                 "1.6.0"
## rsample               "rsample"               "1.1.0"
## Rsamtools             "Rsamtools"             "2.10.0"
## RSQLite               "RSQLite"               "2.2.17"
## rstatix               "rstatix"               "0.7.0"
## rstudioapi            "rstudioapi"            "0.13"
## rsvd                  "rsvd"                  "1.0.5"
## rsvg                  "rsvg"                  "2.3.1"
## rtracklayer           "rtracklayer"           "1.54.0"
## RUnit                 "RUnit"                 "0.4.32"
## rversions             "rversions"             "2.1.1"
## rvest                 "rvest"                 "1.0.3"
## RWeka                 "RWeka"                 "0.4-44"
## RWekajars             "RWekajars"             "3.9.3-2"
## S4Vectors             "S4Vectors"             "0.32.4"
## sass                  "sass"                  "0.4.2"
## scales                "scales"                "1.2.1"
## scatterpie            "scatterpie"            "0.1.8"
## scatterplot3d         "scatterplot3d"         "0.3-42"
## scrime                "scrime"                "1.3.5"
## selectr               "selectr"               "0.4-2"
## sessioninfo           "sessioninfo"           "1.2.2"
## shadowtext            "shadowtext"            "0.1.2"
## shape                 "shape"                 "1.4.6"
## shiny                 "shiny"                 "1.7.2"
## siggenes              "siggenes"              "1.68.0"
## signatureSearch       "signatureSearch"       "1.8.2"
## signatureSearchData   "signatureSearchData"   "1.8.4"
## SingleCellExperiment  "SingleCellExperiment"  "1.16.0"
## skimr                 "skimr"                 "2.1.4"
## slam                  "slam"                  "0.1-50"
```

```
## slider                "slider"                "0.2.2"
## snow                  "snow"                  "0.4-4"
## softImpute            "softImpute"            "1.4-1"
## sourcetools           "sourcetools"           "0.1.7"
## sp                    "sp"                    "1.5-0"
## SparseM               "SparseM"               "1.81"
## sparseMatrixStats     "sparseMatrixStats"     "1.6.0"
## sqldf                 "sqldf"                 "0.4-11"
## SQUAREM               "SQUAREM"               "2021.1"
## STRINGdb              "STRINGdb"              "2.6.5"
## stringi               "stringi"               "1.7.8"
## stringr               "stringr"               "1.4.1"
## styler                "styler"                "1.7.0"
## SummarizedExperiment  "SummarizedExperiment"  "1.24.0"
## sys                   "sys"                   "3.4"
## systemfonts           "systemfonts"           "1.0.4"
## TeachingDemos         "TeachingDemos"         "2.12"
## testthat              "testthat"              "3.1.4"
## textshaping           "textshaping"           "0.3.6"
## TFEA.ChIP             "TFEA.ChIP"             "1.14.0"
## tibble                "tibble"                "3.1.8"
## tidygraph             "tidygraph"             "1.2.2"
## tidymodels            "tidymodels"            "1.0.0"
## tidyr                 "tidyr"                 "1.2.1"
## tidyselect            "tidyselect"            "1.1.2"
## tidytree              "tidytree"              "0.4.1"
## tidyverse             "tidyverse"             "1.3.2"
## timeDate              "timeDate"              "4021.104"
## tinytex               "tinytex"               "0.41"
## tm                    "tm"                    "0.7-8"
## topGO                 "topGO"                 "2.46.0"
## treeio                "treeio"                "1.18.1"
## treemap               "treemap"               "2.4-3"
## truncnorm             "truncnorm"             "1.0-8"
## tune                  "tune"                  "1.0.0"
## tweenr                "tweenr"                "2.0.2"
## tzdb                  "tzdb"                  "0.3.0"
## uchardet              "uchardet"              "1.1.0"
## urlchecker            "urlchecker"            "1.0.1"
## usethis               "usethis"               "2.1.6"
## utf8                  "utf8"                  "1.2.2"
## uuid                  "uuid"                  "1.1-0"
## vctrs                 "vctrs"                 "0.4.2"
## vegan                 "vegan"                 "2.6-2"
## VennDiagram           "VennDiagram"           "1.7.3"
## viridis               "viridis"               "0.6.2"
## viridisLite           "viridisLite"           "0.4.1"
## visNetwork            "visNetwork"            "2.1.2"
## vroom                 "vroom"                 "1.5.7"
## waldo                 "waldo"                 "0.4.0"
## warp                  "warp"                  "0.2.0"
## whisker               "whisker"               "0.4"
```

```
## withr                "withr"             "2.5.0"
## wordcloud            "wordcloud"         "2.6"
## workflows            "workflows"         "1.1.0"
## workflowsets         "workflowsets"      "1.0.0"
## xfun                 "xfun"              "0.33"
## xgboost              "xgboost"           "1.6.0.1"
## XML                  "XML"               "3.99-0.10"
## xml2                 "xml2"              "1.3.3"
## xopen                "xopen"             "1.0.0"
## xtable               "xtable"            "1.8-4"
## XVector              "XVector"           "0.34.0"
## yaml                 "yaml"              "2.3.5"
## yardstick            "yardstick"         "1.1.0"
## yarn                 "yarn"              "1.20.0"
## yulab.utils          "yulab.utils"       "0.0.5"
## zip                  "zip"               "2.2.0"
## zlibbioc             "zlibbioc"          "1.40.0"
## base                 "base"              "4.1.3"
## boot                 "boot"              "1.3-28"
## class                "class"             "7.3-20"
## cluster              "cluster"           "2.1.2"
## codetools            "codetools"         "0.2-18"
## compiler             "compiler"          "4.1.3"
## datasets             "datasets"          "4.1.3"
## foreign              "foreign"           "0.8-82"
## graphics             "graphics"          "4.1.3"
## grDevices            "grDevices"         "4.1.3"
## grid                 "grid"              "4.1.3"
## KernSmooth           "KernSmooth"        "2.23-20"
## lattice              "lattice"           "0.20-45"
## MASS                 "MASS"              "7.3-55"
## Matrix               "Matrix"            "1.4-0"
## methods              "methods"           "4.1.3"
## mgcv                 "mgcv"              "1.8-39"
## nlme                 "nlme"              "3.1-155"
## nnet                 "nnet"              "7.3-17"
## parallel             "parallel"          "4.1.3"
## rpart                "rpart"             "4.1.16"
## spatial              "spatial"           "7.3-15"
## splines              "splines"           "4.1.3"
## stats                "stats"             "4.1.3"
## stats4               "stats4"            "4.1.3"
## survival             "survival"          "3.3-1"
## tcltk                "tcltk"             "4.1.3"
## tools                "tools"             "4.1.3"
## utils                "utils"             "4.1.3"
```
